# High-throughput affinity measurements of direct interactions between activation domains and co-activators

**DOI:** 10.1101/2024.08.19.608698

**Authors:** Nicole DelRosso, Peter H. Suzuki, Daniel Griffith, Jeffrey M. Lotthammer, Borna Novak, Selin Kocalar, Maya U. Sheth, Alex S. Holehouse, Lacramioara Bintu, Polly Fordyce

## Abstract

Sequence-specific activation by transcription factors is essential for gene regulation^1,2^. Key to this are activation domains, which often fall within disordered regions of transcription factors^3,4^ and recruit co-activators to initiate transcription^5^. These interactions are difficult to characterize via most experimental techniques because they are typically weak and transient^6,7^. Consequently, we know very little about whether these interactions are promiscuous or specific, the mechanisms of binding, and how these interactions tune the strength of gene activation. To address these questions, we developed a microfluidic platform for expression and purification of hundreds of activation domains in parallel followed by direct measurement of co-activator binding affinities (STAMMPPING, for Simultaneous Trapping of Affinity Measurements via a Microfluidic Protein-Protein INteraction Generator). By applying STAMMPPING to quantify direct interactions between eight co-activators and 204 human activation domains (>1,500 *K*_d_s), we provide the first quantitative map of these interactions and reveal 334 novel binding pairs. We find that the metazoan-specific co-activator P300 directly binds >100 activation domains, potentially explaining its widespread recruitment across the genome to influence transcriptional activation. Despite sharing similar molecular properties (*e.g.* enrichment of negative and hydrophobic residues), activation domains utilize distinct biophysical properties to recruit certain co-activator domains. Co-activator domain affinity and occupancy are well-predicted by analytical models that account for multivalency, and *in vitro* affinities quantitatively predict activation in cells with an ultrasensitive response. Not only do our results demonstrate the ability to measure affinities between even weak protein-protein interactions in high throughput, but they also provide a necessary resource of over 1,500 activation domain/co-activator affinities which lays the foundation for understanding the molecular basis of transcriptional activation.

## Introduction

Eukaryotic transcription factors (TFs) generally consist of structured DNA binding domains^8^ connected to larger, disordered regions^4,9–11^. TFs that promote gene expression contain activation domains (ADs), often overlapping these disordered regions and recruiting large, multi-domain or multi-subunit transcriptional co-activators (co-As)^5,12^ (**Fig. 1a**). Once recruited to DNA, co-A complexes promote transcription by changing the chromatin state of the gene to make it more permissive to transcription (*e.g.* the acetyltransferase domain^13^ of the chromatin regulator P300^14^), recruiting and/or regulating RNA Polymerase II activity (*e.g.* the TATA-binding protein complex TFIID^15^ and the Mediator complex^16^, which is recruited by the majority of yeast ADs^17^), or by regulating transcriptional elongation (*e.g.* bromodomain-containing proteins^18,19^).

**Figure 1.**
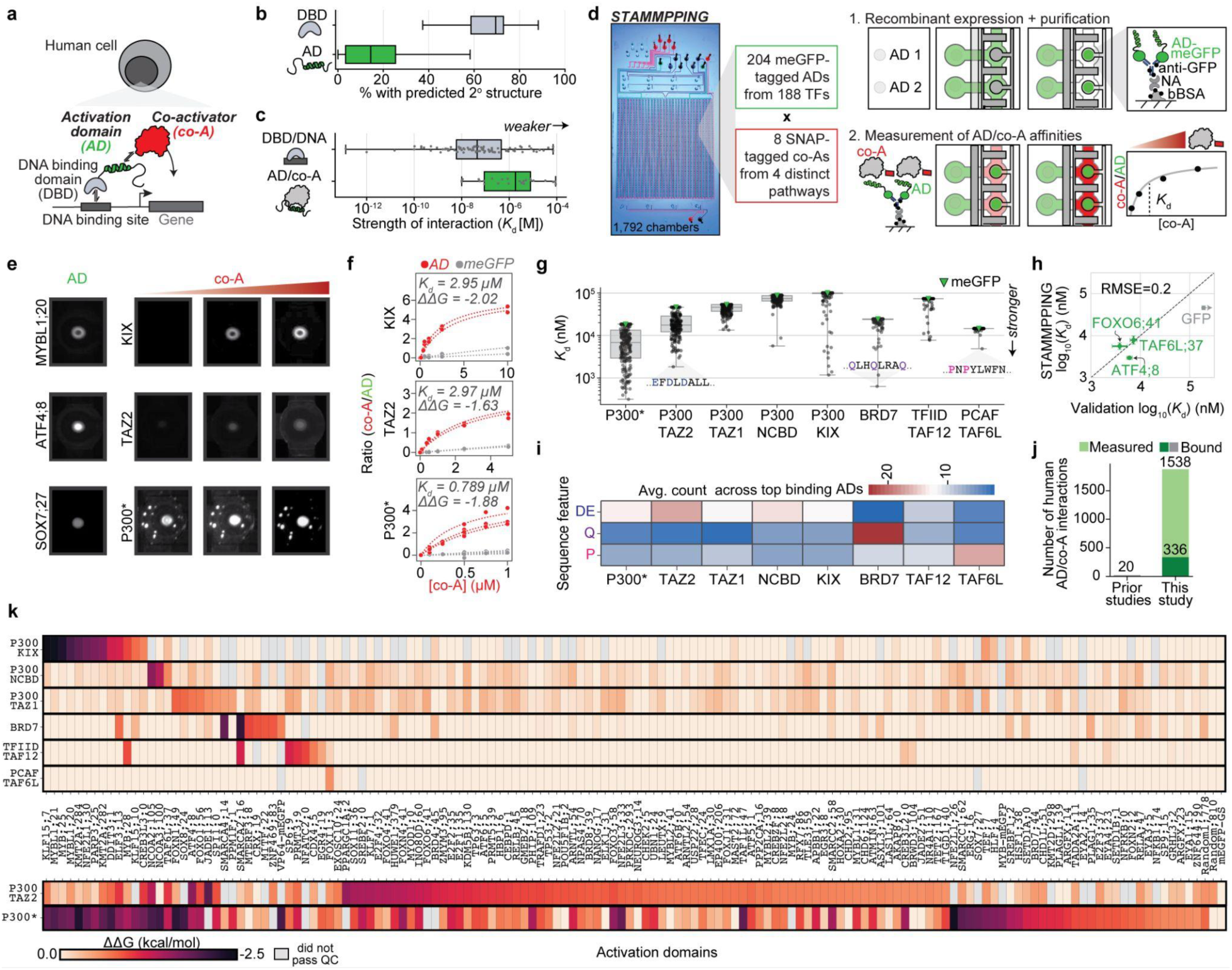
STAMMPPING enables large-scale binding affinity measurements of transcriptional protein complexes. **a,** Cartoon schematic of relevant molecular interactions for transcription. **b,** Fraction of residues predicted by AlphaFold to exist within helical or strand secondary structures for annotated ADs^20^ and DBDs (Uniprot). **c,** Distribution of previously measured affinities for DBD/DNA^47^ and AD/co-A interactions. **d,** Image of STAMMPPING microfluidic device filled with food dye (left) and cartoon overview of STAMMPPING assay (right). NA=neutravidin, bBSA=biotinylated bovine serum albumin. **e**, Example images of ADs after expression and purification (left) and after incubation with increasing concentrations of three different co-As. All images within each co-A have the same intensity scale, but not across different co-As. P300 shows formation of high intensity punctae at high concentrations. **f**, Example binding curves corresponding to the chamber images in **e** (red) and for meGFP and three different co-As (gray). Lines indicate Langmuir isotherm fits; ΔΔG values are calculated by subtracting each meGFP/co-A’s ΔG (ΔG=-RTln(*K*_d_)) from each AD/co-A interaction’s ΔG. P300’s *K*_d_s are marked with an asterisk to indicate the presence of punctae that may reduce the effective concentration of soluble protein such that the *K*_d_s represent an upper limit. **g**, Swarm plot showing individual *K*_d_s for each AD/co-A; box plots show median *K*_d_ and interquartile range for all interactions with a given co-A. Affinities for meGFP indicate apparent non-specific binding threshold (green); AD/co-A interactions with weaker affinities were set at this value and *K*_d_s were indicated as lower limits. Annotations indicate sequence features enriched within the highest affinity binders for P300 TAZ2 (ATF4;8), BRD7 (SMARCA2;16), and TAF6L (FOXI1;3). **h,** Comparison between measured STAMMPPING and microscale thermophoresis affinities for ATF4;8, FOXO6;41, TAF6L;37 binding to P300’s TAZ2. Mean ± standard deviation across two to three replicates shown. RMSE denotes mean squared error calculated only on the AD variants and based on log_10_-transformed *K*_d_ values in nM (1.1-fold). Arrow on meGFP indicates a lower limit of detection. **i,** Average number of acidic, glutamine, and proline amino acids within each co-A’s most strongly bound ADs. **j,** Number of interacting human AD/co-A pairs identified here and before this study. **k,** Heatmap of ΔΔG values for each co-A (rows) interacting with each AD (columns), ΔΔGs were calculated in reference to meGFP/co-A’s *K*_d_. ADs with no detectable binding to any co-A or domain tested (44) are not shown. AD ΔΔG values are ordered manually by each co-A’s *K*_d_ beginning with P300 KIX. Gray indicates measurements that did not pass quality control.

In prior work, we systematically identified 374 ADs within human TFs using a high-throughput reporter assay called HT-recruit^20^. Consistent with other work, these ADs shared common sequence characteristics: they tended to be disordered (**Fig. 1b**), negatively charged, and contained prolines, serines and large hydrophobic residues^2,3,17,20–27^. On the basis of these similarities, deep learning models have been developed to predict which sequences activate transcription with reasonable accuracy^17,27–29^. Nevertheless, the mechanisms by which Ads recognize and bind co-A partners remain unclear: efforts to organize these AD sequences into motifs have not been fruitful, with the only common grammar being clusters of hydrophobic residues surrounded by negative charge^22,24,28^. Given that ADs must physically interact with co-As in order to activate, perhaps efforts to analyze AD sequence preferences have failed because they group together heterogenous ADs with possible distinct co-A binding preferences.

Compared to our knowledge about AD sequence characteristics, we know relatively little about the molecular basis of AD/co-A interactions, outside of a few key examples^30–35^. This is likely because both AD/co-A affinities are weak (**Fig. 1c**), making them difficult to detect experimentally, and because bound-state complexes appear to retain a high degree of conformational heterogeneity^7,33,36–38^. The view that ADs are “negative noodles^21^” that leverage their acidic residues to promote exposure of nearby hydrophobic residues^22^ to bind co-A hydrophobic pockets implies that AD/co-A interactions are generally promiscuous. However, depletion studies in human cells have found certain co-As are only functionally required at specific enhancers^39,40^, implying that co-As have some degree of specificity. Mapping AD/co-A molecular specificities using a sensitive technique capable of quantifying even weak binding events in high-throughput would provide a mechanistic framework for AD sequence-to-function relationships.

### High-throughput protein-protein binding affinity measurements

To map and quantify affinities for direct interactions between thousands of AD/co-A pairs, we created STAMMPPING, for <u>S</u>imultaneous <u>T</u>rapping of <u>A</u>ffinity <u>M</u>easurements via a <u>M</u>icrofluidic <u>P</u>rotein-<u>P</u>rotein <u>IN</u>teraction <u>G</u>enerator (**Fig. 1d, Supplementary Figure 1**). STAMMPPING uses a 1,792 chamber valved microfluidic device^41–45^ to simultaneously express and purify many AD ‘bait’ proteins and quantify their binding affinities to co-A ‘prey’ proteins. To begin each experiment, we print a library of linear DNA templates encoding C-terminally monomeric enhanced green fluorescent protein (meGFP)-tagged ADs on a glass slide and align devices to arrayed DNA spots. Next, we introduce *in vitro* transcription/translation (IVTT) mix into each chamber to express all AD-meGFP ‘bait’ proteins in parallel, immobilize each AD onto an anti-GFP antibody-patterned spot within each chamber, and wash out excess protein and IVTT reagent from the device while a closed pneumatic valve protects the AD-patterned spots from flow. Next, we quantify affinities to co-A ‘prey’ proteins by introducing AlexaFluor647-labeled SNAP-tagged co-As at multiple concentrations, allowing interactions to come to equilibrium, and measuring fractional occupancies at each step by quantifying the ratio of bound Alexa647 (co-A) to meGFP (AD) intensities. Finally, we fit the resulting concentration-dependent binding curves to a Langmuir isotherm, enabling us to estimate 1,792 AD/coA binding affinities (*K*_d_) in parallel per experiment.

To select ‘bait’ ADs for further study, we attempted on-chip expression of 1,156 tiles (80 amino acids long) that overlap the 374 ADs previously shown to activate transcription via HT-recruit^20^ (**Extended Data Fig. 1a-b**, **Supplementary Table 1, Methods**). As AD/co-A interaction *K*_d_s are likely in the micromolar range and difficult to detect (**Fig. 1c, Supplementary Table 2**), we maximized expected signal-to-noise by selecting activating tiles with high on-chip expression for further characterization. Of the 805 tiles that were successfully cloned, 310 reproducibly expressed at high levels (four times the standard deviation above the mean of empty chambers) across two replicate devices (**Extended Data Fig. 1c-d, Supplementary Table 3**); we then selected the tile from each protein with the highest activated transcription in cells^20^ for further study (204 tiles, hereafter ADs).

To select ‘prey’ co-As, we attempted to express C-terminally SNAP-tagged full-length proteins (and, in some cases, individual folded domains) from five co-As that regulate transcription in diverse ways: the acetyltransferase P300/CBP (which has four annotated, well-folded activation binding domains (KIX, NCBD, TAZ1 and TAZ2 connected by long disordered linkers^6^); subunits from the TATA-binding protein complex TFIID; subunits from the structural complex Mediator; a subunit from P300/CBP-associated factor (PCAF) complex; and bromodomain-containing BRD proteins. Eleven of the 17 co-As expressed well *in vitro* (**Extended Data Fig. 1e, Supplementary Figure 2**). Two of the remaining co-As were commercially available as either FLAG-tagged or GST-tagged proteins, allowing detection of binding via co-introduction of fluorescent anti-FLAG or anti-GST. All co-As remained in solution at the highest concentrations profiled except for full-length P300 and BRD4’s BRD domain, which showed visible high-intensity punctae (**Fig. 1e, Supplementary Figure 2, Supplementary Note 1**). For these proteins, the concentration of soluble protein is likely lower than the quoted values such that fitted *K*_d_s represent an upper limit and are annotated as such.

We then applied STAMMPPING to directly quantify affinities between the final curated set of 204 ADs and 13 co-As and domains (full-length P300; the P300 domains KIX, TAZ2, NCBD, and TAZ1; the TFIID domains TAF9 and TAF12; full-length MED15 and its KIX domain along with MED9; PCAF subunit TAF6L; full-length BRD7 and the bromodomain from BRD4), yielding 2,652 concentration-dependent binding curves (**Fig. 1f**, **Supplementary Figure 2**). Each experiment was performed at least twice and included meGFP and two random control sequences as negative controls, the transcriptional activator MYB as a positive control for KIX-binding, the strong synthetic viral activator VP64, and 32 empty chambers to sensitively detect any cross-chamber contamination. Langmuir isotherm fits to measured concentration-dependent binding returned estimated apparent affinities (*K*_d_s) for each of these co-As to the 204-member AD library (**Fig. 1f**, **Supplementary Figure 2**). We observed reproducible and sequence-specific binding above background (median Pearson r^2^>0.56) for 8/13 co-As and domains (full-length P300; the P300 domains KIX, TAZ2, NCBD, and TAZ1; the TFIID domain TAF12; PCAF subunit TAF6L; and full-length BRD7) (**Extended Data Fig. 1f, Supplementary Figure 2**), yielding 1,538 high-confidence AD/co-A binding affinities (**Supplementary Table 4**). The remaining co-As either did not bind any ADs (TAF9, BRD4’s bromodomain), had variable binding across experiments (MED9, MED15), or appeared to bind relatively non-specifically (MED15 KIX).

AD/co-A affinities differed across co-As and domains (**Fig. 1g**), with full-length P300 binding most tightly (median *K*_d_s <1 µM for 19 ADs). Co-As were either promiscuous (full-length P300 and P300’s TAZ2) or specific (P300’s KIX, NCBD, TAZ1, full-length BRD7, TFIID’s TAF12, PCAF’s TAF6L). Forty-four ADs did not show significant binding to any tested co-A and were significantly less negatively charged than co-A-binding ADs (*p*-value=0.01, Mann-Whitney U test, one-tailed). Measurements of the labeled SNAP protein alone showed no detectable binding up to the highest concentrations, confirming a lack of binding between the two fluorescent tags (**Extended Data Fig. 1g**). STAMMPPING measurements (**Extended Data Fig. 1h**) agreed with previously published values^46^ and orthogonal measurements via microscale thermophoresis, a solution-phase label-free method (**Fig. 1h, Extended Data Fig. 1i**), confirming STAMMPPING accuracy and that fluorescence tagging and surface immobilization do not impact affinities.

To test if individual co-As and their domains preferentially bind ADs with particular sequence features, we quantified enrichment of amino acids within the most strongly bound ADs for each co-A (*K*_d_s below 10th percentile for promiscuous co-As and *K*_d_s three or more sigma below the mean for specific co-As, **Methods**). P300 preferentially bound most strongly to ADs relatively enriched in acidic residues (*p*-value=0.02 compared to BRD7, TAF12, TAF6L, Mann-Whitney U test, one-tailed), BRD7 bound most strongly to ADs relatively enriched in glutamine residues (*p*-value=0.01 compared to P300, TAF12, TAF6L), and TAF6L bound most strongly to an AD with relatively more prolines (*p*-value=0.05 compared to BRD7, P300, TAF12) (**Fig. 1i**). Together, these data suggest that co-As preferentially bind distinct subsets of ADs.

Out of the 1,538 AD/co-A interactions with high-confidence affinities (**Supplementary Table 4**), we classified 336 AD/co-A pairs as binding significantly above negative controls (*p*-value<=0.05, one-sided t-test, **Methods**). All but two of these pairs are novel, resulting in over 15x more interactions than have previously been reported (**Fig. 1j**). To visualize the network of AD/co-A interactions, we calculated differences in relative binding affinity (ΔΔGs) using each meGFP/co-A interaction as a reference point (**Fig. 1k**). As expected, measured ΔΔG values for full-length P300 correlate with the summed ΔΔG values for individual P300 subunits (Pearson r^2^=0.68, **Extended Data Fig. 2a-e**). Most interactions involve P300 and P300 TAZ2, each binding >80 ADs (**Fig. 1k** bottom). Other co-As were more specific, binding several ADs each (**Fig. 1k** top). For example, BRD7’s top binders were ADs from SMARCA2 and SMARCA4; P300’s NCBD domain, consistent with prior studies on CBP’s NCBD domain^35^, most tightly bound ADs from NCOA2 and NCOA3.

### Distinct binding mechanisms for two co-A domains

What physical features drive binding between largely disordered ADs and particular co-As (**Fig. 1k**)? <u>S</u>hort <u>Li</u>near <u>M</u>otifs (SLiMs) are minimal motifs within intrinsically disordered regions (IDRs) that mediate selective protein interactions across many eukaryotic proteins^49,50^, such as the SLiM in MYB’s AD (ELELLL) and the P300 co-A domain KIX^51,52^. In some cases (as for MYB/KIX), SLiMs lack a stable 3D structure but become ordered upon binding^53^; in other cases, AD/co-A interactions are shape-agnostic, ‘fuzzy’, and dynamic^33^ (as for yeast ADs that bind Mediator subunit 15^7,17,38,54^). The observed differences in co-A specificity (**Fig. 1k**) and amino acid enrichment amongst the most tightly bound ADs (**Fig. 1i**) suggest that different co-As employ distinct molecular strategies to specifically recognize their cognate ADs. To investigate these mechanisms, we turned to the KIX and TAZ2 domains from P300 (**Fig. 2a**). KIX is relatively specific, binding only 16 ADs, while TAZ2 is more promiscuous, binding 90 ADs with affinities tighter than meGFP alone (**Fig. 2b**, **Supplementary Table 4**).

**Figure 2.**
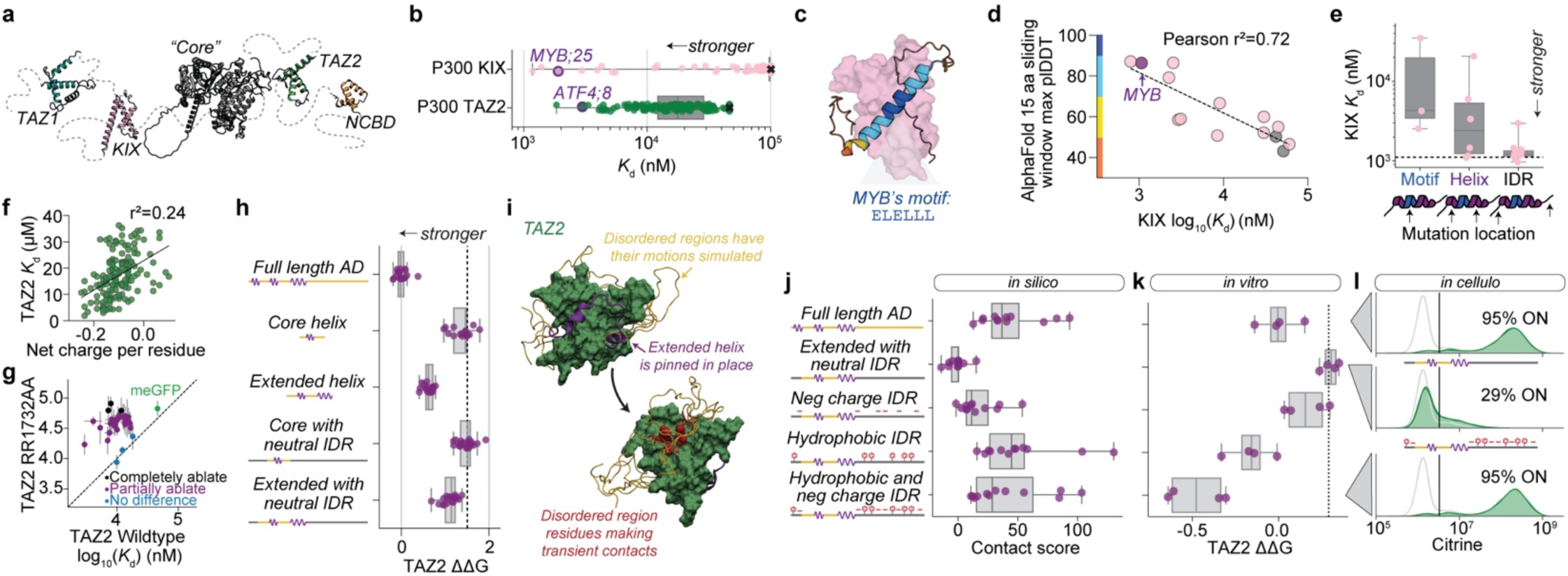
Binding determinants are distinct for promiscuous and specific co-A domains. **a,** Schematic showing AlphaFold-predicted structures for P300’s four activation binding domains and a catalytic core consisting of BRD, CH2, HAT, and ZZ domains connected by long disordered linkers^6^. **b**, Boxplots show median affinities and interquartile range for P300’s KIX and TAZ2 domains binding to different AD variants (see Fig. 1g). **c**, AlphaFold-Multimer model of MYB;25’s AD binding to P300 KIX. Colorbar depicts confidence score. **d**, Scatter plot showing AlphaFold-Multimer-predicted confidence scores *vs.* STAMMPPING affinities (*K*_d_s) for 12 KIX-binding ADs and two non-binding ADs (gray). **e**, Boxplots show median affinities and interquartile range for P300’s KIX domain binding to different rationally designed mutants of MYB;25. Helix indicates MYB’s predicted helix (as in 2c); IDR indicates intrinsically disordered region (see **Extended Data Fig. 3c** for sequences). **f**, Scatter plot showing each AD sequence’s net charge per residue *vs.* measured affinities with *K*_d_<40 µM for the P300 domain TAZ2; r^2^ indicates Spearman correlation coefficient. **g**, Pairwise comparison of AD affinities for RR1732AA mutant and WT TAZ2. Black markers indicate ADs with complete loss of TAZ2 binding (one-tailed t-test comparing mutant TAZ2 binding between each AD and meGFP); purple markers indicate ADs with partially ablated binding (two-tailed t-test comparing WT and mutant TAZ2); blue markers indicate ADs with no difference in their binding between WT and mutant TAZ2; green marker indicates meGFP. **h**, Boxplots show median relative affinities (ΔΔGs) and interquartile range for P300’s TAZ2 domain binding to different rationally designed mutants of ATF4;8. Core only, core with neutral IDR, extended core, and extended with neutral IDR sequence affinities differ significantly from the native AD with Mann-Whitney two-sided U test *p*-values of 1.2e-7, 1.62e-8, 7.7e-8, and 5.06e-8. **i**, Still shots from simulations of ATF4;8/TAZ2. Residues in yellow had their motions simulated, residues in purple were pinned in place, and residues in red are hydrophobic residues that were within eight angstroms of one another. **j**, Box plots showing *in silico* normalized contact scores between IDRs and folded TAZ2 domains from nine independent all-atom Monte Carlo simulation replicates for rationally designed ATF4;8 mutants (see **Extended Data Fig. 4e** for sequences) interacting with TAZ2. **k**, Box plots show median measured *in vitro* relative affinities (ΔΔGs) and interquartile range for core helix embedded within different IDR variants; *p*-values quantify statistical significance of differences from the WT IDR via Mann-Whitney two-sided U test (*p* = 0.029 for neutral IDR and neutral IDR with only charged residues replaced and 0.11 for IDRs with hydrophobic or hydrophobic and negative residues replaced). **l**, *In cellulo* flow cytometry distributions of a subset of the same ATF4;8 mutants (gray triangle indicates mutant identity) recruited to minCMV Citrine reporter gene integrated in K562 cells. Gray curves have no dox and serve as no recruitment negative controls; green curves have dox where the proteins are recruited to the reporter gene.

### KIX binding requires a SLiM displayed within an extended helical element

Binding of the AD within MYB to P300 KIX is essential for hematopoietic cell proliferation^55^, and aberrant expression of MYB is associated with leukemia and breast and colon cancers^56^. Prior efforts to develop small molecules that inhibit MYB/KIX binding^57^ and NMR structural data^58^ have revealed that KIX has two distinct binding surfaces that recognize ADs via a similar SLiM^52^ ([DEST][LMYI]..[LIF][LIV]^59^, a variant of the ΦXXΦΦ motif^60^, where Φ is a hydrophobic residue and X is any residue). Consistent with prior observations, most STAMMPPING-identified KIX-binding ADs contained this annotated SLiM (12/16 (75%), **Extended Data Fig. 3a**). However, many ADs with the same SLiM did not bind KIX (28/192 (15%), **Extended Data Fig. 3a**), instead binding TAZ2 here (*n*=20) and in prior studies^33^the SLiM alone is not sufficient to accurately predict KIX binding.

While only a few AD/co-A complexes have been structurally determined, AlphaFold-Multimer^61^ shows promise for predicting such complexes. Consistent with prior NMR structures^62^, predicted structures of the KIX/MYB complex using AlphaFold-Multimer revealed MYB’s ELELLL SLiM docked within the known binding pocket (**Fig. 2c**). AlphaFold-Multimer also docked 11 additional newly-identified KIX-binding ADs within the known binding pockets and confidence metrics correlated strongly with measured affinities (Pearson r^2^=0.72) (**Fig. 2d, Extended Data Fig. 3b**), suggesting that KIX recognition depends on specific interactions between helical SLiM residues and KIX’s binding pockets. To test if predicted high-confidence alpha helices extending beyond the annotated SLiM (**Fig. 2c**, light blue) impact binding, we generated 30 MYB mutants with sequence changes in either the ELELLL SLiM, this extended helical region, or surrounding IDRs (**Fig. 2e, Extended Data Fig. 3c, Supplementary Figure 3, Supplementary Table 5**). As expected, mutations to the known SLiM reduced affinity (with impacts ranging from 3- to 30-fold depending on the position mutated) and IDR mutations had minimal impacts on binding (with the exception of a sequence where the entire N-terminal IDR was shuffled). Mutations to single key residues (*e.g.* I295Y) or more extensive substitutions overlapping the extended helical region reduced affinity by up to 20-fold, confirming that KIX/AD binding specificity depends on a helical motif and extended secondary structure interactions that tune these affinities.

### Electrostatic interactions play a larger role in TAZ2-binding compared to KIX

In contrast to KIX, TAZ2-binding is promiscuous (90/204 ADs) and relatively uncharacterized. While TAZ2 and TAZ1 domains are highly similar^63,64^, fewer ADs bound TAZ1 above background levels; most TAZ1 binders also bound TAZ2 (21/37). Attempts to discover candidate SLiMs essential for TAZ2 binding from AD populations that either do or do not bind TAZ2 (using XSTREME^65^ and FIMO^66^) returned several motifs (DLDLDMF and EELDLAE). However, only 48% of TAZ2-binding ADs contained these motifs and they were also found in 11% of non-binding TAZ2 ADs (**Extended Data Fig. 3d**), indicating that SLiMs alone are not sufficient to determine TAZ2 recognition.

These candidate SLiMs contain multiple negatively-charged residues, suggesting that electrostatics may play an important role. Consistent with this, AD overall net charge was correlated with affinity for TAZ2-bound (but not KIX-bound) ADs (**Fig. 2f, Extended Data Fig. 3e-f**) and TAZ2 contains many positively-charged solvent-facing residues (**Extended Data Fig. 3g**). To test the degree to which these residues are crucial for TAZ2 AD recognition, we purified TAZ2 constructs in which two positively charged surface residues were mutated to neutral alanines (RR1732AA), individually shown to reduce binding to the P53 AD^67^); the mutant protein appeared folded when assessed via circular dichroism (**Extended Data Fig. 3h**). Measured concentration-dependent binding to 31 different TAZ2-binding ADs revealed reduced binding for most ADs (28/31, two-tailed t-test between WT and mutant TAZ2) (**Fig. 2g**, **Supplementary Figure 4, Supplementary Table 6**). Binding to three ADs was reduced to background (meGFP) levels, suggesting predominant binding at the RR1732 site (one-tailed t-test). An additional three ADs were unaffected, consistent with predominant binding at an alternate site or sites; the remainder were impacted to varying degrees, suggesting contributions to binding from multiple sites across TAZ2. To further confirm the importance of electrostatics, we purified KIX and TAZ2 in a salt-free buffer (**Extended Data Fig. 3i**) and carried out STAMMPPING experiments in varying salt concentrations from 100 to 440 mM NaCl (**Supplementary Figure 5, Supplementary Table 7**). TAZ2 formed visible aggregates after purification in salt-free buffer and high-intensity fluorescent punctae in STAMMPPING experiments (**Extended Data Fig. 3i-j**); nevertheless, concentration-dependent binding behavior could be fit by a single-site binding model across salt concentrations (**Supplementary Figure 5**). TAZ2 affinities were significantly more sensitive to changes in salt than their KIX counterparts (*p*-value=1.53e-4, two-sample Kolmogorov-Smirnov test, **Extended Data Fig. 3j**), confirming electrostatic interactions play a larger role in TAZ2 binding.

### Intrinsically disordered regions within TAZ2-binding ADs contribute to affinity

To further investigate determinants of TAZ2 binding specificity, we selected three ADs that strongly bound TAZ2 and full-length P300 (ATF4;8, FOXO1;56, and TAF6L;37, **Supplementary Table 4**), systematically deleted 10 amino acid (aa) fragments scanning across each AD, and quantified TAZ2 affinities via STAMMPPING (**Extended Data Fig. 4a-c**, **Supplementary Figure 6**, **Supplementary Table 8**). Deletion of multiple 10-aa regions within each AD weakened affinities (20/23, **Extended Data Fig. 4a-c**). Segments that overlapped short sections of AlphaFold-predicted helical structure (hereafter ‘core helices’) had some of the strongest effects, but loss of segments within IDRs also reduced affinities (**Extended Data Fig. 4a-c**). Consistent with a functional role for these IDRs, AD/TAZ2 AlphaFold-Multimer complex prediction confidence scores did not correlate with TAZ2 binding strength (r*^2^*=0.06, **Extended Data Fig. 4d**).

To quantify the importance of these IDRs for TAZ2 recognition, we measured TAZ2 affinities for ATF4, FOXO1, and TAF6L AD constructs comprised of the ‘core helix’ or an extended version of this helix (‘extended helix’) either alone, embedded within a neutral IDR of the same length, or within the native context (**Extended Data Fig. 4e**). For all three ADs, the ‘core helix’ alone was bound substantially more weakly than the full-length sequence and extending the helices only partially rescued binding (**Fig. 2h, Extended Data Fig. 4f**). Adding a surrounding neutral IDR of the same length either had no effect or slightly reduced affinities, suggesting that IDRs with specific charge properties are essential for proper TAZ2 binding.

IDR-mediated impacts on TAZ2/AD binding affinity could stem from either changes in the AD conformational ensemble or contacts between IDRs and the folded TAZ2 domain. To identify candidate IDR/TAZ2 interactions, we turned to all-atom simulations of the bound AD/co-A complex using the ABSINTH implicit solvent model^68^ in which the protein backbones of TAZ2 and the AD ‘extended helix’ were held fixed while the IDR backbone was allowed to freely sample (**Fig. 2i**). The native ATF4 IDR made frequent transient contacts with the TAZ2 surface that were not seen in simulations for ATF4 with a neutral IDR, nor for MYB/KIX interactions (**Extended Data Fig. 5a-d**). Additional simulations established that adding only the negatively charged or hydrophobic residues from the WT IDR to the neutral IDR sequence were sufficient to partially and fully restore WT-like AD/TAZ2 interactions, respectively (**Fig. 2j, Extended Data Fig. 5e**). Experimental STAMMPPING measurements of affinities for TAZ2 binding the same constructs *in vitro* showed highly similar patterns (**Fig. 2k, Supplementary Figure 7**, **Supplementary Table 9**); slight differences in the relative importances of adding negatively charged residues likely stem from either differences in electrostatic conditions between simulations and experiments or immobilization of the core interaction during simulations. Recruiting the same sequences *in cellulo* upstream of a minimal promoter and quantifying reporter expression showed the same pattern of activation (**Fig. 2l, Extended Data Fig. 5f-g**). This suite of *in silico*, *in vitro*, and *in cellulo* measurements establish that: (1) while both KIX- and TAZ2-binding ADs employ interactions within alpha helices to bind hydrophobic pockets of folded domains, TAZ2-binding ADs further rely on transient hydrophobic and electrostatic interactions between IDRs and folded domains for recognition (**Extended Data Fig. 5h**), and (2) that *in silico* and *in vitro* affinity measurements can accurately predict activation changes in cells.

### Multivalent wiring between ADs and co-As modulates affinity and occupancy

P300 and many of the ADs that bind it contain multiple binding sites (**Fig. 3a**, **Extended Data Fig. 6a**). P300 contains multiple protein-binding domains^6^ that each interact with multiple ADs (**Fig. 1k**). Within these domains, P300’s KIX domain contains two distinct hydrophobic binding pockets (MYB and MLL)^57,58^. Finally, most human ADs (>61%) contain more than one region necessary for activation^20^ (**Extended Data Fig. 6a**). A variety of models have previously been developed to describe how multivalency might impact macroscopic affinities and molecular recruitment^69–73^. To gain intuition as to the impact of multivalency within interactions between freely diffusing co-As and immobilized ADs (as in STAMMPPING experiments and for TFs bound to DNA regulatory sequences), we analytically modeled binding for ADs and co-As with either one or two sites each. AD polyvalency (a co-A with one site binding to ADs containing multiple homotypic sites) increases the maximum amount of co-A bound (*R_max_*) without altering affinity (**Fig. 3b** left, pink vs blue curves). By contrast, co-A polyvalency (a co-A with two sites binding to a single AD) increases affinity additively in *K_a_* space (**Fig. 3b** middle, orange vs blue curves). Avidity (pairs of cognate binding sites on ADs and co-As) increases affinity multiplicatively without altering *R_max_* until co-A concentrations greatly exceed the interaction *K_d_*^72^ (**Fig. 3b**, right, red vs blue curves, **Extended Data Fig. 6b**).

**Figure 3.**
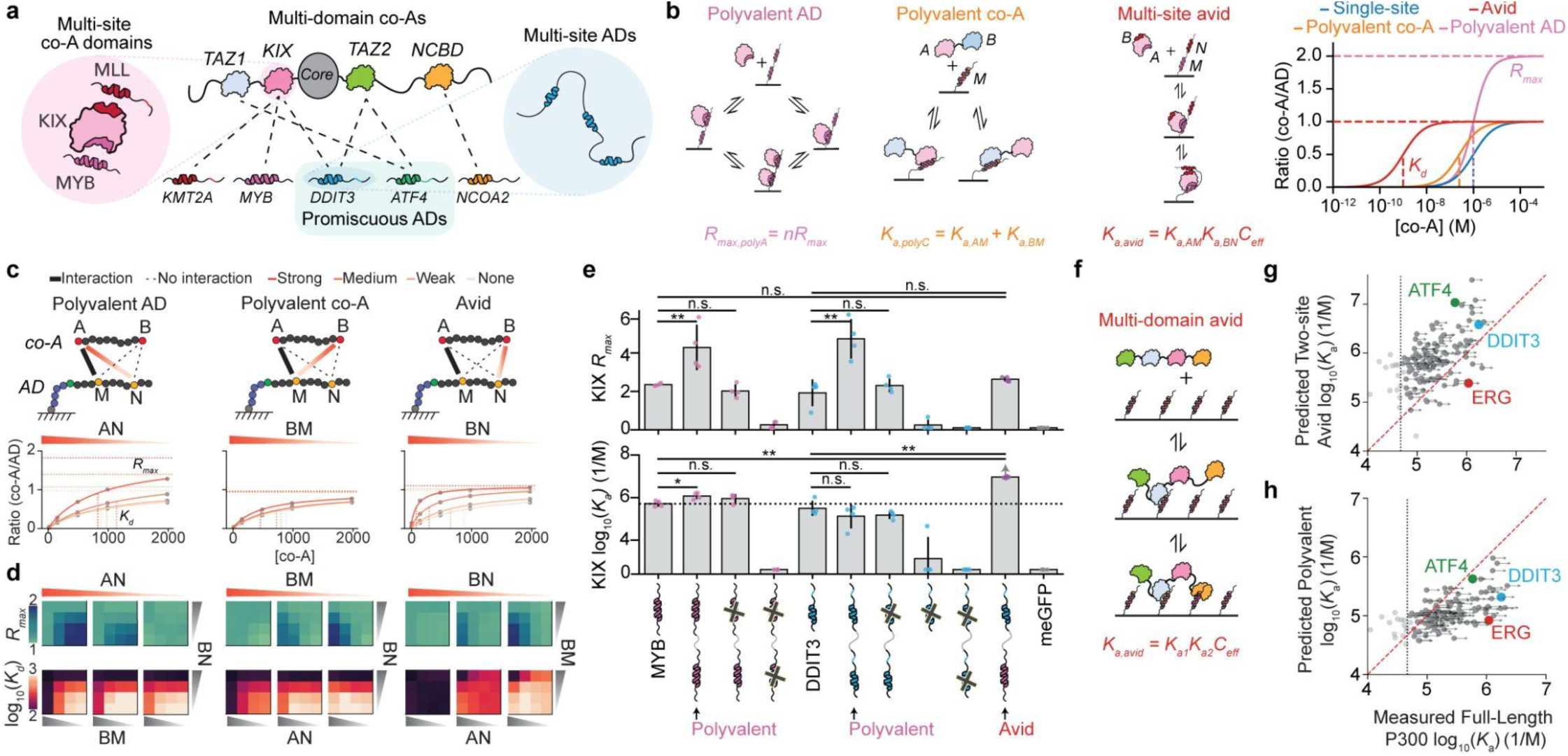
AD/co-A systems exploit multivalency in distinct ways. **a,** Cartoon schematic showing multivalency from multi-site ADs, multi-site folded domains within co-As, and multi-domain co-A proteins (*e.g.* P300). Dotted lines depict identified binding interactions (**Supplementary Table 4**). **b**, Cartoon schematics of polyvalent and avid binding between a multi-site AD and single-site co-A or multi-site co-A, respectively, paired with simulated binding curves (right, see **Methods**). **c**, PIMMS-simulated binding curves across increasing strengths of interactions between AN (left), BM (middle), and BN (right) while keeping AM interaction strength strong and turning off all other interactions. Dotted vertical lines indicate fitted *K_d_* and dotted horizontal lines indicate fitted *R_max_*. Light tan curve represents the single-site condition. **d**, Fitted affinity (*K_d_*) and occupancy (*R_max_*) as a function of interaction strengths between individual contacts within two site co-A/two site AD PIMMS simulations. **e**, Measured affinity (*K_a_*) and occupancy (*R_max_*) for P300 KIX binding synthetic multi-site ADs. n.s.: p>0.05, *:p<0.05, **:p<0.001 (two-sided t-test). Arrow denotes limit of detection; “x” marks denote L302A mutation for MYB constructs and sequence shuffle of residues 0-15 for DDIT3 constructs. **f**, Cartoon schematic of avid binding between a multi-domain co-A and multi-site AD. **g**, Predicted affinity from “two-site avid” model *vs.* measured affinity for full-length P300/AD binding interactions; “Two-site avid” prediction calculated by *K_a,avid_2site_* = *K_a1_ K_a2_ C_eff_*, where *K_a1_* and *K_a2_* are affinities of the two tightest-binding P300 subunits for a given AD and *C_eff_* estimated as shown in **Extended Data Fig. 8e**. Dashed line indicates average meGFP negative control; red dashed line indicates the identity line. Arrows indicate the presence and direction of measurement limits due to uncertainty in P300 soluble concentration. **h**, Predicted affinity from polyvalent model *vs.* measured affinity for full-length P300/AD binding interactions. Arrows indicate the presence and direction of measurement limits.

To validate and extend our analytical models to consider impacts of varying combinations of monomer interaction affinities, we used PIMMS (Polymer Interactions in Multicomponent MixtureS, a coarse-grained lattice-based Monte Carlo simulation engine, **Extended Data Fig. 6c**) to model 160 unique combinations of affinity microstates within a two-by-two site co-A/AD network. For each configuration, we quantified the number of co-As bound to surface-immobilized ADs at equilibrium as a function of concentration and extracted apparent affinities and occupancies (*K*_d_ and *R*_max_) by fitting these data to a Langmuir isotherm (**Methods**, **Extended Data Fig. 6d, Supplementary Table 10**). PIMMS predictions were highly similar to analytical model predictions (Pearson r^2^≥0.68, **Extended Data Fig. 6e-f**). However, results of PIMMS simulations (**Fig. 3c-d)**, additionally revealed that small differences in individual co-A/AD interaction affinities could yield large and nonlinear changes in macroscopic affinities in the avid binding case (**Fig. 3d**, bottom right heatmap), even when the total energy of network interactions is constant (**Extended Data Fig. 6g**). These simulations show that wiring between AD binding sites and co-A binding surfaces can significantly alter *K*_d_s, tuning between polyvalent-like and avid-like configurations (**Fig. 3d**, bottom heatmaps).

Next, we tested these model predictions by quantifying concentration-dependent binding for P300’s KIX domain binding to a set of surface-immobilized synthetic ADs consisting of one or two homo- or heterotypic pairs of the KIX-binding ADs MYB and DDIT3 (**Supplementary Figure 8, Supplementary Table 11**). Consistent with a polyvalent binding model (**Fig. 3b** left) and PIMMS simulations (**Extended Data Fig. 7a**), homotypic MYB-GS-MYB constructs (**Extended Data Fig. 7b**) increased KIX occupancy (*R*_max_) at saturation relative to single-site MYB (1.8+/-0.5-fold), with little or no increase in affinity (*K*_d_ change <1.1-fold; **Fig. 3e** left). Similar results were observed for DDIT3-GS-DDIT3 constructs (2.5+/-0.8-fold; *K*_d_ change <1.1-fold; **Fig. 3e** middle). Constructs with a mutation in one of the MYB or DDIT3 sites (MYB L302A^74^ or DDIT3 N-term shuffle, **Extended Data Fig. 7c**) bound similarly to a single AD alone (*p-*value>0.017), establishing that the remainder of the AD did not contribute to binding. These results are consistent with a model in which both MYB-GS-MYB and DDIT3-GS-DDIT3 bind in a polyvalent manner, implying that DDIT3 exclusively binds a single pocket on KIX, analogous to MYB^62^. By contrast, a heterotypic MYB-GS-DDIT3 synthetic AD significantly increased affinity (>1.7 kcal/mol), saturating at the first measured concentration of KIX such that estimated *K*_d_s represent an upper bound (**Supplementary Figure 8**) with little to no increase in *R_max_* (MYB *p*-value=0.01, DDIT3 *p*-value=0.09, **Fig. 3e**, far right). This result is consistent with an avid binding model in which MYB and DDIT3 simultaneously interact with distinct regions on the KIX domain, implying that DDIT3 is specific for the MLL pocket (or any non-MYB pocket on KIX). A similar trend in which heterotypic AD affinities were significantly tighter than for either AD alone was observed for MYB and KMT2A (**Extended Data Fig. 7d-e**, MYB *p*-value=2.2e-4, KMT2A *p*-value=1.7e-5). Overall, multiple homotypic AD sites recruited more KIX molecules while multiple heterotypic AD sites primarily increased KIX affinity.

Individual ADs could potentially interact with any of P300’s four annotated TF-binding domains and/or regions within the linkers or ‘core’. Given P300’s size, individual P300 molecules could bridge neighboring ADs on the same microfluidic spot surface and lead to avid binding (**Fig. 3f, Extended Data Fig. 8a-e**). However, measured *in vitro* P300 binding was mostly weaker than predictions of a simplified two-site avid model calculated using an effective concentration (*C*_eff_) estimated by considering the volume subtended by a single P300 molecule anchored at the tightest binding site and the surface density of displayed ADs (**Fig. 3g**, **Extended Data Fig. 8f, Supplementary Note 2**). While most ADs show approximately polyvalent binding to P300 (*e.g.* ATF4, **Fig. 3h**, **Extended Data Fig. 8g**), others bind up to 10-fold more tightly than expected by a polyvalent model (*e.g.* DDIT3, **Fig. 3h**), in line with previous observations in yeast’s Med15^7^. Thus, most AD/P300 interactions bind with affinities between the predictions of polyvalent and avid models without a need to include further complexity such as allovalency^69,70^ (**Supplementary Note 3**).

### P300’s KIX domain binding affinities *in vitro* predict activation strength *in cellulo*

Results thus far establish that most ADs directly bind P300 (**Fig. 4a**) when measured at similar AD densities as predicted at our HT-recruit reporter (**Extended Data Fig. 8c,h**). However, the relationship between AD/P300 affinities and levels of activation remains unknown. Across 200 ADs from 185 transcriptional proteins, ADs with the strongest P300 affinities (<1 µM) drove the strongest activation in HT-recruit activation measurements^20^, with median activation levels decreasing monotonically for medium (1-10 µM) and weak (>10 µM) affinity binders (**Fig. 4b**, strong *p*-value=4.64e-5, medium *p*-value=6.35e-4, Mann Whitney, one-sided). However, there is significant variation within each of these bins, consistent with a model in which ADs that activate transcription at higher levels than predicted from P300 binding measurements bind additional co-As to drive gene expression.

**Figure 4.**
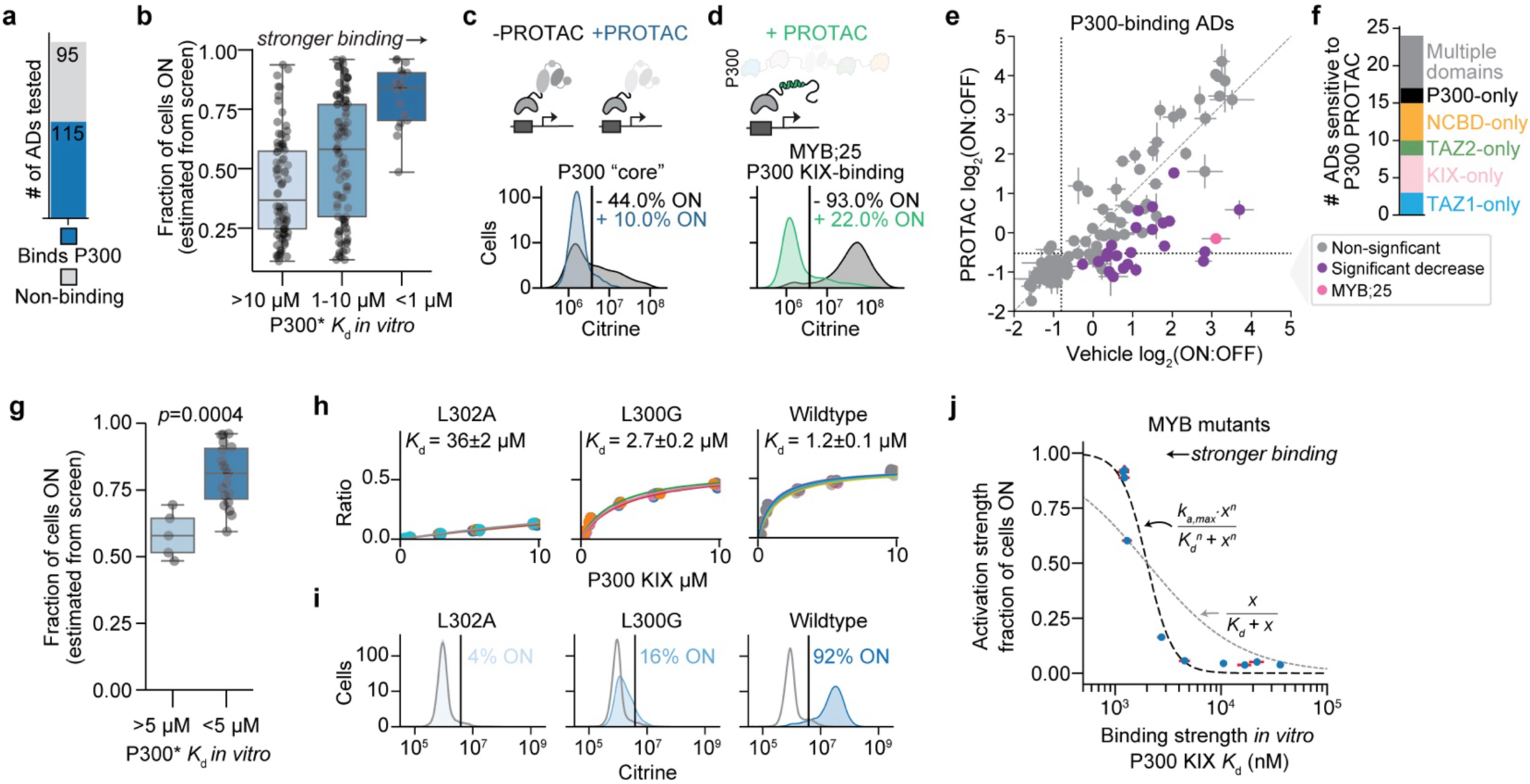
Binding strength to P300’s KIX domain *in vitro* is predictive of cellular activation strengths. **a**, Number of ADs that bind to full-length P300 (blue) in STAMMPPING experiments. **b**, Comparison between HT-recruit activation scores that were estimated from screen enrichment scores to fraction ON using a calibration curve based on flow cytometry individual measurements (**Methods**) from ref:^20^; 48 hours of recruitment and STAMMPPING-measured affinities for P300. P300 binding strength binned as strong (<1 µM), medium (1-10 µM), and weak (>10 µM). P300’s *K*_d_s are marked with an asterisk to indicate the presence of punctae that may reduce the effective concentration of soluble protein such that *K*_d_s represent an upper limit (**Supplementary Note 1**). **c**, Recruitment of P300’s “core” to minCMV promoter in the absence (black) or presence (colors) of 500 nM of dCBP-1, a PROTAC molecule that selectively degrades P300/CBP (*p*-value<10^-6^, two sample Kolmogorov-Smirnov test). **d**, Recruitment of MYB tile 25 to minCMV promoter in the absence (black) or presence (colors) of 500 nM of dCBP-1 (*p*-value<10^-6,^ two sample Kolmogorov-Smirnov test). **e,** Pairwise comparison between HT-recruit screen enrichment scores between vehicle- and drug-treated cells (18 hours of recruitment). P300-binding ADs with a *p*-value ≤ 0.05 and an average difference greater than 0.5 were considered significantly decreased (purple points). **f**, Distribution of P300 and P300 domains that PROTAC-sensitive ADs bind. **g**, Comparison between HT-recruit activation scores that were estimated from screen enrichment scores to fraction ON using a calibration curve based on flow cytometry individual measurements (**Methods**) from ref:^20^; 48 hours of recruitment and STAMMPPING-measured affinities for P300 for the 24 PROTAC-sensitive P300-binding domains. Mann Whitney U test, one-sided. **h**, STAMMPPING-measured concentration-dependent binding of P300’s KIX domain to individual MYB mutants (markers) and Langmuir isotherm fits (lines); each color indicates data from a different chamber; *K*_d_s are the average across at least four replicates and errors are the standard deviations. **i**, *In cellulo* flow cytometry distributions for individual MYB mutants recruited to minCMV Citrine reporter gene integrated in K562 cells. Gray curves have no dox and serve as no recruitment negative controls; blue curves have dox where the proteins are recruited to the reporter gene. **j**, Relationship between fraction of cells ON from individual recruitment measurements and STAMMPPING-measured affinities for P300’s KIX domain. Black dashed function plotted with *k_a,max_* set to 1 and a returned best-fit value for *n* 3.58; both black and gray functions are plotted with *x*=2e3 (**Extended Data Fig. 9f**).

To systematically identify ADs that only bind P300/CBP *vs.* those that likely bind multiple co-As, we carried out an HT-recruit activation screen for all 204 STAMMPPING-tested ADs in the presence or absence of a PROTAC molecule that selectively binds endogenous CBP and P300’s BRD domain within the catalytic core^77^. If an AD activates transcription via specific recruitment of P300/CBP alone, PROTAC-induced degradation of P300/CBP should drive a significant loss of activation. As expected, PROTAC treatment of cells in which we recruited either P300’s core catalytic domain or the MYB AD (which activates transcription via direct binding to P300 KIX) dramatically reduced activation (**Fig. 4c-d**). Within the 111 P300-binders recovered in HT-recruit (**Extended Data Fig. 9a**, **Supplementary Table 12**), 80 activated after 18 hours of recruitment; activation for 24 of them was significantly altered, suggesting these ADs recruit P300/CBP relatively specifically; the rest showed non-significant changes, suggesting these ADs recruit additional co-As beyond P300/CBP to drive their activation (**Fig. 4e**, **Extended Data Fig. 9b**). Individual tests of several PROTAC-insensitive ADs (*e.g.* P300 TAZ2-binding ATF4 and TAF6L) confirmed these results (**Extended Data Fig. 9c**). Most PROTAC-sensitive ADs specifically bound single P300 domains in STAMMPPING (**Fig. 4f**). Across the 24 PROTAC-sensitive P300-binding domains, stronger *in vitro* P300 binding was associated with stronger activation in cells (*p*-value=4e-3, Mann Whitney, one-sided, **Fig. 4g, Extended Data Fig. 9d**); however, because most PROTAC-sensitive ADs bind P300 tightly (*K*_d_s of 0.45-10 µM), the range of observed activation strengths was relatively small (from 48 to 96%).

To systematically assess how variations in P300 binding affinities across a wider dynamic range impact activation, we measured activation strengths for 10 MYB mutants shown to bind KIX with affinities (*K*_d_s) spanning from 1-40 µM in STAMMPPING assays (**Fig. 2e, Supplementary Figure 3**). Levels of transcriptional activation (percentage of cells “on”) for these MYB mutants varied from 4-92% (**Supplementary Figure 9**), with tighter-binding MYB mutants increasing both average levels of activation and the fraction of activated cells (**Fig. 4h-i, Extended Data Fig. 9e**). The input/output function between affinities and fraction of activated cells was best fit by an ultrasensitive binding model^78^ with a Hill coefficient of 3.6 such that mutants that only slightly reduce KIX binding *in vitro* drive large changes in activation (*e.g.* L300G, **Fig. 4h-j**, **Extended Data Fig. 9f**). We also observe an ultrasensitive relationship for another PROTAC-sensitive KIX-binding AD, KMT2A (**Supplementary Figure 10-11, Extended Data Fig. 9g-h**), suggesting ultrasensitivity between transcriptional strength and AD/co-A binding strength could be a general feature amongst specific P300-binding ADs.

## Discussion

Metazoan-specific P300^79^ is widely used as a marker for transcriptionally active enhancers in human cells^80–82^. Consistent with this central role in transcription, we find that full-length P300 directly binds most tested ADs (115/204, *K*_d_s=0.3-14 µM) via distinct mechanisms. While most ADs directly bind the TAZ2 domain, in part by electrostatic and IDR-mediated interactions, others only bind the KIX, NCBD, or TAZ1 domains. Some specifically contact more than one domain, providing additional opportunities for multivalent regulation by differentially tuning either affinity or occupancy. While observed patterns of amino acid enrichments^17,20,22,28^ provide a useful tool for identifying ADs^23,27,83,84^, universal models of AD/co-A recognition do not exist. Our integrated high-throughput *in silico, in vitro*, and *in cellulo* measurements yield a global view of AD/co-A specificity landscapes that establish weak and disordered interactions can encode extensive specificity and partner flexibility, consistent with low-throughput NMR studies^36,85^.

Transcription from a gene’s promoter is controlled in part by enhancers^86^, and disruption of specific enhancer-promoter contacts can result in developmental defects or cancer^87–89^. How do these enhancers know which promoters to specifically activate? One proposed model, ‘biochemical compatibility,’ posits that this specificity is encoded in specific protein-protein interactions connecting enhancers and promoters^90^. STAMMPPING measurements revealed strong specificity between some ADs and partner co-As (*e.g.* BRD7). This discovery of highly specific interactions within this network could open up new avenues for the development of small molecules designed to inhibit binding interactions^91^ or target disease-associated transcription factors for degradation^92,93^. Thus, the library of AD/co-A measurements, and extension of the methods detailed here to map the rest of the vast TF/cofactor network, will provide essential quantitative information for next-generation drug discovery.

The input/output function connecting AD/P300 affinities and transcription in human cells was unknown. Here, observed levels of transcription depend quantitatively and cooperatively on AD/co-A affinities. This observed ultrasensitivity could result from multiple rTetR-activators binding simultaneously to the nine TetO multivalent synthetic promoter. Alternatively, this cooperativity could arise from positive feedback loops resulting from either P300-mediated deposition and binding of H3K acetylation^94^, or cooperative homotypic assembly of many P300 molecules^95,96^. Future studies systematically varying single parameters could clarify these mechanisms.

Several models have been developed to explain the impacts of multivalency on binding^70^. Beyond the avid and polyvalent models described above, allovalent and “fuzzy binding” models are special cases of polyvalent binding that tune the local concentration of available binding partners and number of binding sites, respectively. While prior single-molecule tracking measurements in human cells have suggested P300 avidly binds ADs^97^, here we find P300 *in vitro* binding exists on a spectrum between polyvalent and avid predictions. In addition, we find that weak AD/co-A interactions enriched in disorder can non-linearly amplify macroscopic affinities if their multivalent wiring becomes avid^71^. In cells, co-As with multiple interaction domains, such as P300, could recruit additional soluble ADs to promote molecular chaining and nucleate large macromolecular self-assemblies^98,99^ that could have similar nonlinear impacts on occupancies. Finally, dynamic changes in disordered region lengths between neighboring binding sites^71,100^ could directly rewire multivalent networks and transcription in response to changing nuclear conditions^101–103^. Thus, the measurements and models presented here on IDR-containing AD/co-A multivalent interactions provide critical information required to directly and quantitatively predict multivalent phenomena observed in cells^104–108^.

Although transcription is the most highly enriched cellular process involving proteins with large disordered regions^50^, other cellular processes such as signaling^109^, circadian rhythms^110^, and cell cycle regulation^111^ all depend on communication between IDRs and folded domains. Moreover, an emerging paradigm suggests that IDR-mediated interactions with folded domains are driven by both sequence specificity (whereby linear motifs enable canonical molecular recognition) and chemical specificity (whereby complementary chemistry enables dynamic interactions without the requirement for a precise amino acid sequence)^73,112^. Both of these modes emerge here, demonstrating that STAMMPPING can reliably quantify 1000s of weak protein-protein interactions across different binding mechanisms. This, in turn, opens the door to the systematic investigation of dense regulatory networks dominated by weakly interacting nodes.

**Extended Data Figure 1.**
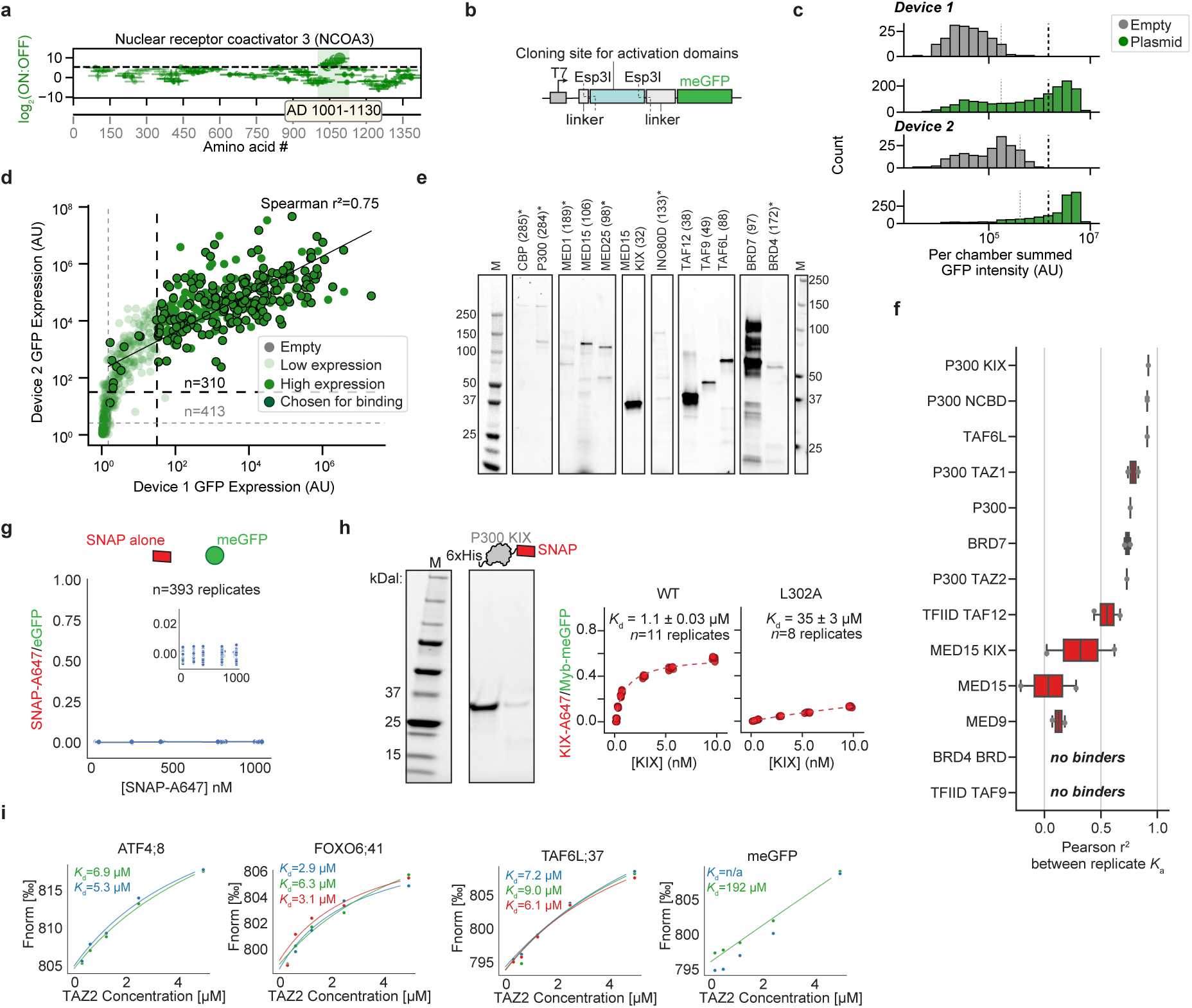
**Related to Figure 1 a**, Example tiling plot illustrating how the 1,156 member activating tile library was designed. For STAMMPPING measurements, we cloned all activating tiles identified in prior cell-based chromatin regulator and TF tiling screens (ref:^20^) (*e.g.* the six tiles within the green highlighted region shown here). **b**, Activating tiles were cloned as a pool into a common plasmid encoding expression of tiles fused to a C-terminal meGFP backbone under control of a T7 promoter; individual variants were isolated and sequence-validated via uPIC-M^48^. **c**, Distributions of summed GFP intensities for empty and plasmid-containing chambers across two STAMMPPING devices; light dashed lines indicate expression thresholds (calculated as mean ± three standard deviations above each device’s empty chambers; dark dashed line indicates threshold (1.5 x 10^6^) used to distinguish chambers that expressed strongly (four times the standard deviation above the mean of empty chambers). **d**, Pairwise comparison of summed GFP intensities for individual chambers across two STAMMPPING devices; light dashed lines indicate expression thresholds (calculated as mean ± three standard deviations above each device’s empty chambers; dark dashed line indicates threshold (1.5 x 10^6^) used to distinguish chambers that expressed strongly (four times the standard deviation above the mean of empty chambers). **e**, Example gel showing a subset of IVTT-expressed Alexa647-labeled SNAP-tagged proteins before purification. Stars indicate poorly expressed proteins. Markers on the right are for the BRD proteins and markers on the left are for all other proteins. **f**, Pearson r^2^ values between replicate *K*_a_ measurements (1/*K*_d_) across all ADs for each co-A. Box edges indicate lower and upper quartiles; whiskers indicate data range extents. **g**, Concentration-dependent binding curves for A647-labeled SNAP interacting with surface-immobilized meGFP. Markers indicate summed microfluidic well spot fluorescence ratios for 393 chambers; line indicates the average values. **h**, Example gel showing A647-labeled SNAP-tagged P300 KIX (left) and measured concentration-dependent binding curves for this protein interacting with the WT MYB AD (middle) and a single amino acid mutant (L302A, right). Markers indicate measured intensity ratios for individual chambers; dashed line indicates Langmuir isotherm fit to all data returning the annotated apparent *K*_d_. **i**, Measured concentration-dependent binding curves from microscale thermophoresis for an unlabeled P300 TAZ2 domain binding to four different labeled, IVTT-expressed ADs. *K*_d_s were fit using Monolith software.

**Extended Data Figure 2.**
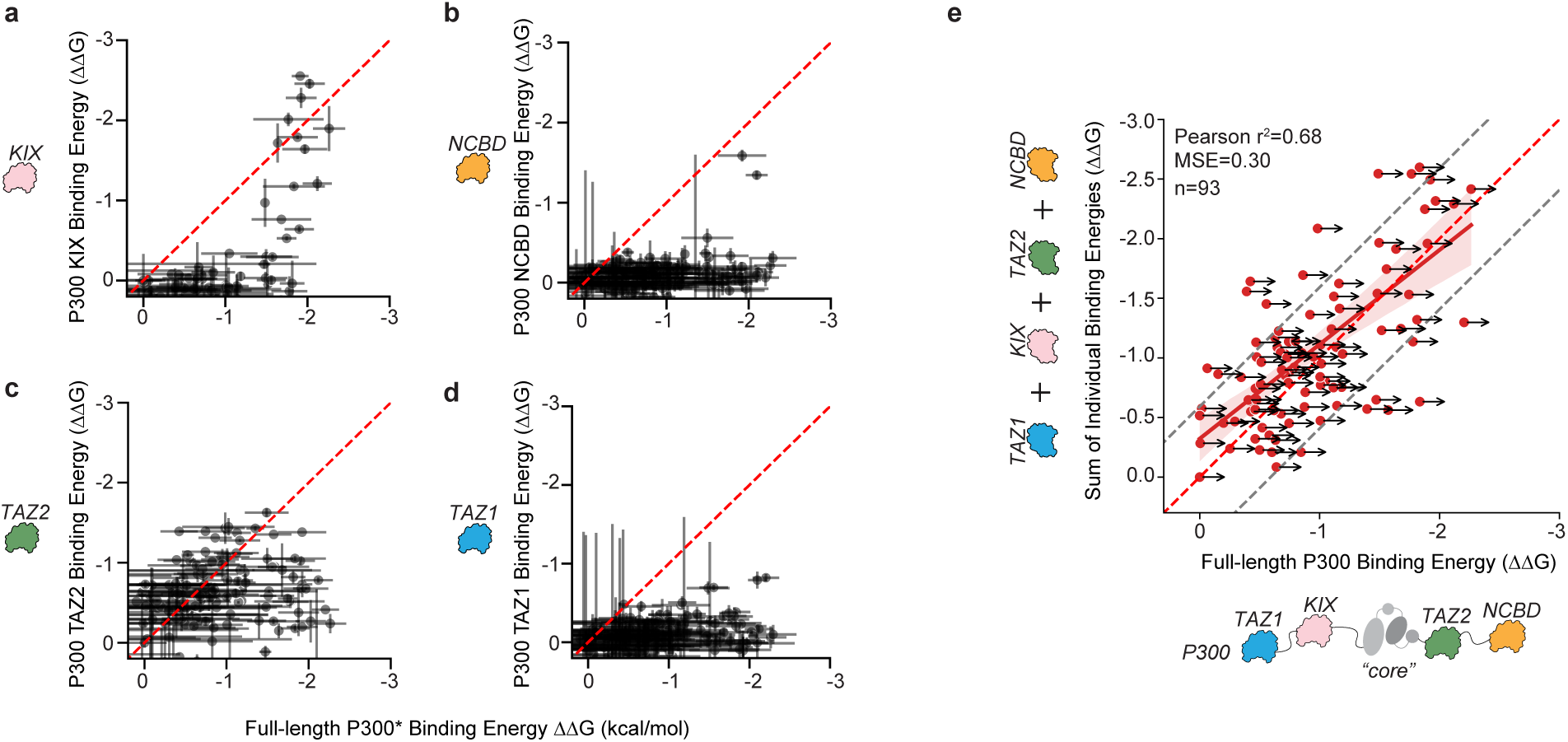
**Related to Figure 1 a-d**, Measured relative binding affinities (ΔΔGs) for individual P300 domains *vs.* full-length P300. Each marker represents an AD variant. Error bars indicate standard deviation. P300’s binding energies are marked with an asterisk to indicate the presence of punctae that may reduce the effective concentration of soluble protein such that *K*_d_s represent an upper limit (points could lie further to the right, Supplementary Note 1). **e**, Scatter plot comparing sum of binding affinities (ΔΔGs) across P300 subunits with measured ΔΔG for full-length P300. Dotted gray lines denote +/- 1 k_b_T from 1:1 line. ADs with confident binding energies (standard deviation < 0.5 kcal/mol) are plotted. Error bars represent standard deviation and propagated standard error for the sum of energies. Arrows indicate the presence and direction of measurement limits.

**Extended Data Figure 3.**
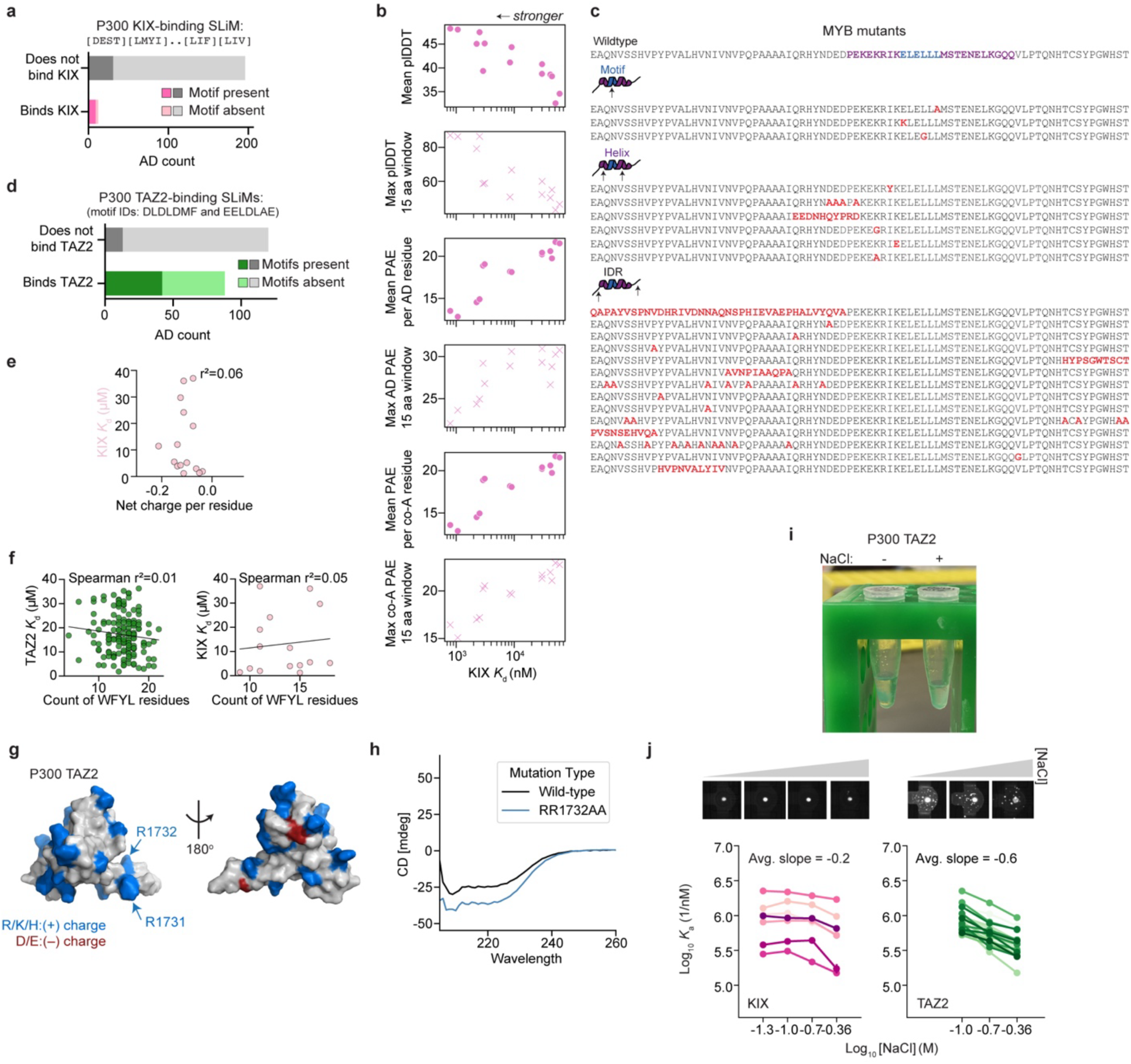
**Related to Figure 2 a**, Number of ADs that contain a KIX-binding SLiM (regular expression shown in the figure) and bind (bottom) or don’t bind (top) KIX in STAMMPPING measurements (**Supplementary Table 4**). ADs that do not contain the SLiM are shown in each category on top as lighter colors. **b**, Scatter plots showing various AlphaFold-Multimer complex prediction scores plotted against STAMMPPING-measured affinities to P300 KIX. **c**, MYB sequences containing mutations within the SLiM/motif, the helix in the AlphaFold-Multimer predicted complex (see Fig. 2c), or the intrinsically disordered regions. Within each category, sequences are ordered from largest to smallest effect on affinity. **d**, Number of ADs that contain either of two TAZ2-binding SLiMs and bind (bottom) or don’t bind (top) TAZ2 in our measurements. ADs that do not contain the SLiM are shown in each category on top as lighter colors. **e**, Scatter plot showing each AD sequence’s net charge per residue *vs.* measured affinities with *K*_d_<40 µM for the P300 domain KIX; r^2^ indicates Spearman correlation coefficient. **f**, Scatter plot showing the relationship between each AD sequence’s count of WFYL residues and P300 TAZ2 domain’s affinity (left) and P300 KIX domain’s affinity (right). Spearman r^2^ shown for each; plot includes all ADs with *K*_d_<40 µM. **g**, PDB structure 2MZD with the P53 AD removed highlighting the charged residues on P300 TAZ2. **h**, Circular dichroism spectra for purified WT TAZ2 and the RR1732AA mutant. **i**, Pictures showing purified TAZ2 reproducibly formed visible aggregates in salt-free buffer. Concentration is roughly the same in the two tubes. **j**, Pictures (top) of KIX (left) and TAZ2 (right) as salt is increased and scatter plots (bottom) showing salt dependence of measured association constants for seven KIX-binding ADs and 22 TAZ2-binding ADs. Average slope is shown across different ADs shown in different colors (*p*-value=1.53e-4, two-sample Kolmogorov-Smirnov test). Measured *K*_d_s represent upper limits because TAZ2 punctae likely reduce the effective concentration of soluble protein.

**Extended Data Figure 4.**
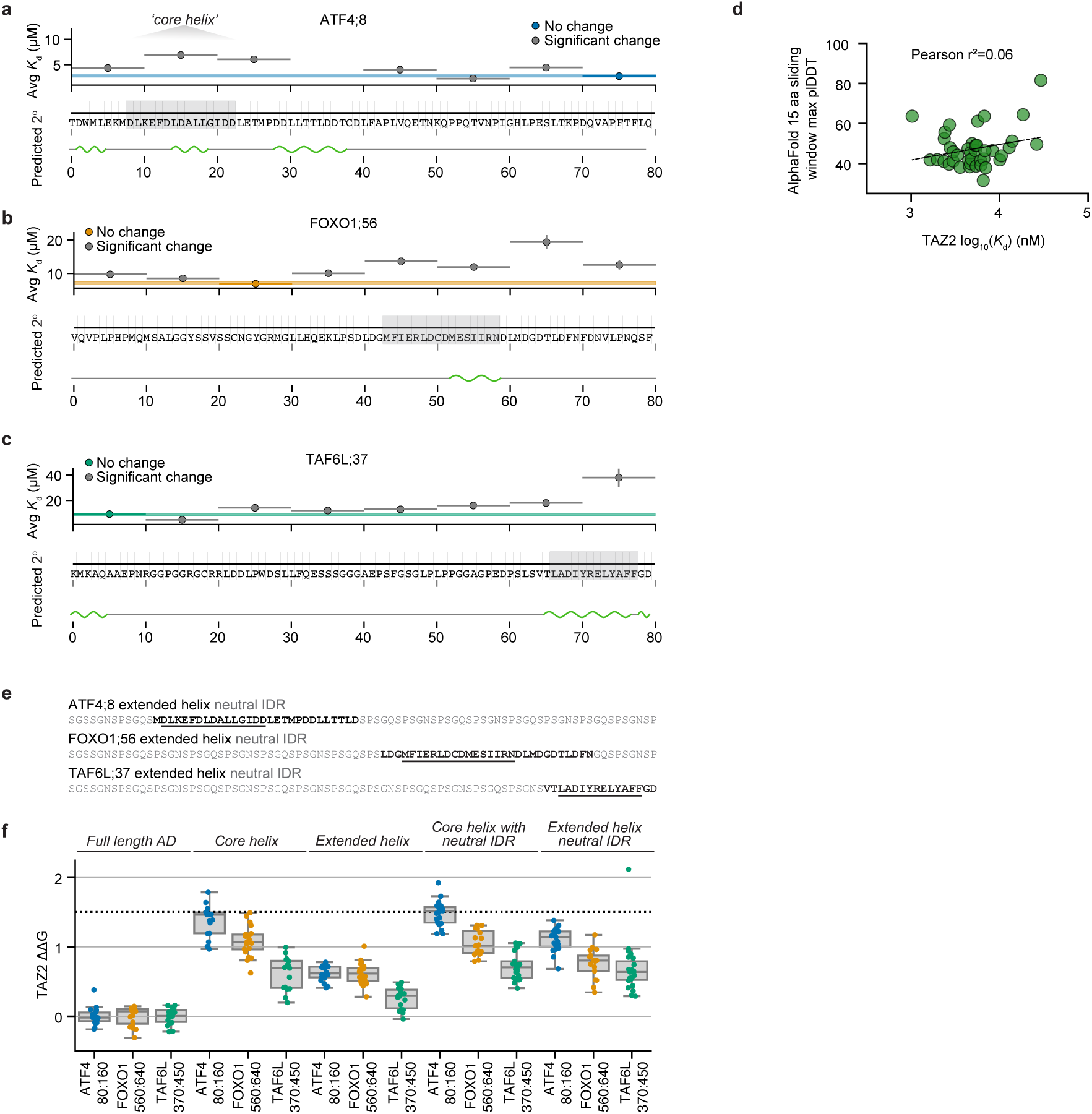
**Related to Figure 2 a**, STAMMPPING-measured affinities (*K*_d_s) (top), sequence (middle) and AlphaFold-predicted secondary structure (bottom, green regions denote alpha helices, prediction from whole protein sequence) for constructs containing 10 amino acid deletions (deletion locations denoted by horizontal lines) within the TAZ2-binding AD tile ATF4;8. Blue line indicates measured affinity for the wild-type AD tile; vertical height of the line denotes the average standard error of fits across replicates. Deletions are colored by statistical significance (one-sided z-test): deletions with *p*-values <0.05 (gray) are significantly different from WT, while the rest are not (colors). Grey box highlights the ‘core helix’. **b**, Results as shown in a for the FOXO1;56 AD tile. **c**, Results as shown in a for the TAF6L;37 AD tile. **d**, Scatter plot showing AlphaFold-Multimer-predicted confidence scores *vs.* STAMMPPING affinities for 54 TAZ2-binding ADs. **e**, Rationally designed neutral IDR sequences that were tested in panel f (last column). Underline indicates ‘core helix’ residues. **f**, Boxplots show median ΔΔG and interquartile range for TAZ2 binding to different rationally designed mutants of ATF4;8, FOXO1;56, and TAF6L;37. ΔΔGs were calculated in reference to each wild-type AD’s *K*_d_. Dashed line is the noise floor for this experiment, defined as the average relative affinity for negative control meGFP binding to TAZ2.

**Extended Data Figure 5.**
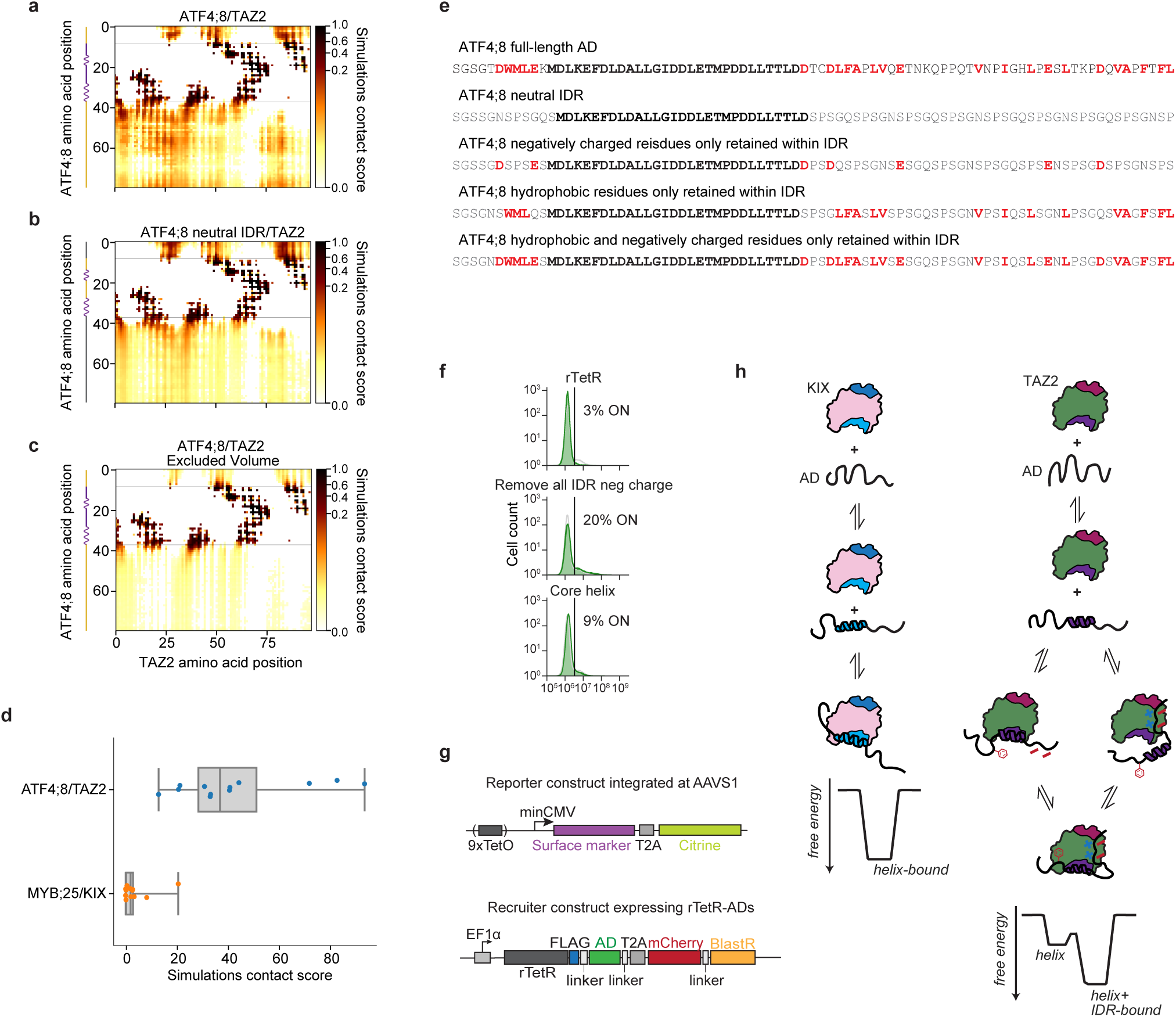
**Related to Figure 2 a**, Heatmap showing *in silico* normalized contact scores between IDRs from ATF4;8 (y-axis) and folded TAZ2 (x-axis) averaged over nine independent all-atom Monte Carlo simulations. Dashed lines indicate where the ‘extended helix’ was pinned. White pixels indicate unsampled regions. Darker pixels represent higher contact frequencies. **b**, Heatmap showing *in silico* normalized contact scores between ‘neutral’ IDRs (second sequence in panel **e** from ATF4;8, y-axis) and folded TAZ2 (x-axis) from nine independent all-atom Monte Carlo simulations. **c**, Heatmap showing *in silico* normalized contact scores between IDRs from ATF4;8 (y-axis) and folded TAZ2 (x-axis) from nine independent all-atom Monte Carlo ‘Excluded Volume’ negative control simulations where all physical interactions other than steric repulsions were turned off. **d**, Boxplot shows median *in silico* normalized contact scores and interquartile range between IDRs from MYB;25 and KIX and IDRs from ATF4;8 and TAZ2. **e**, Rationally designed sequences tested in Fig. 2j**-l**. Black residues are wild-type; gray indicate modified neutral residues; red indicate rescued wild-type negatively charged or certain hydrophobic residues. **f**, Flow cytometry distributions of ATF4;8 mutants and controls recruited to minCMV Citrine reporter gene integrated in K562 cells. Gray curves have no dox and serve as no recruitment negative controls; green curves have been treated with dox such that proteins are recruited to the reporter gene. **g**, Schematic of dual reporter gene consisting of a synthetic surface marker (Igκ-hIgG1-Fc-PDGFRβ) and the fluorescent protein citrine. Lentiviral recruitment construct expressing the dox-inducible DNA binding domain rTetR fused to ADs. **h**, Cartoon summary of results.

**Extended Data Figure 6.**
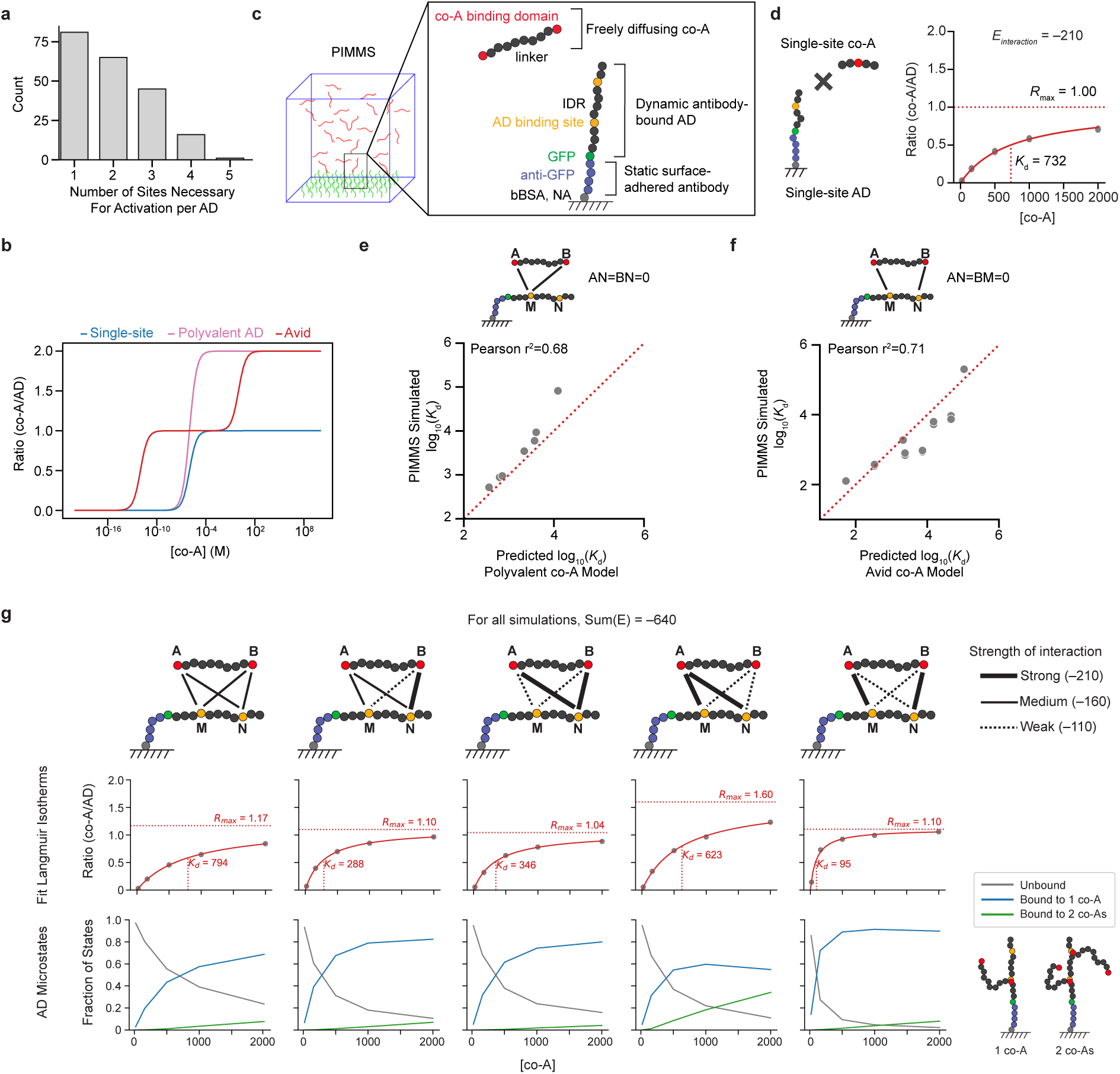
**Related to Figure 3 a**, Number of sites that reduce activation when deleted per 80-amino-acid AD tile; data analyzed from ref:^20^ **b**, Simulated binding curves of polyvalent and avid binding between a multi-site AD and single-site co-A or multi-site co-A, respectively (see **Methods**). **c**, Schematic of PIMMS simulation in which freely diffusing co-As with two binding sites contact surface-immobilized ADs containing two binding sites; the binding strength of each interaction varies between non-binding, weak, medium, and strong interactions (**Methods**). **d**, For each set of AD and co-A pairs, we enumerated AD/co-A contacts across five co-A concentrations (gray points), fitting *K_d_* and *R_max_* as for experimental data (**Methods**). **e**, Comparison between PIMMS-simulated affinity for two site co-A/one site AD cases and predictions from analytical models for the polyvalent co-A case. Red dashed line indicates the identity line. **f**, Comparison between PIMMS-simulated affinity for two site co-A/two site AD cases and predictions from analytical models for the avid case. Red dashed line indicates the identity line. **g**, Fitted Langmuir isotherms (top) and observed frequencies of AD microstates (bottom) across increasing co-A concentration for two site co-A/two site AD simulations in which the sum of interaction energies was held constant at –640 units; cartoon at right shows example observed microstates.

**Extended Data Figure 7.**
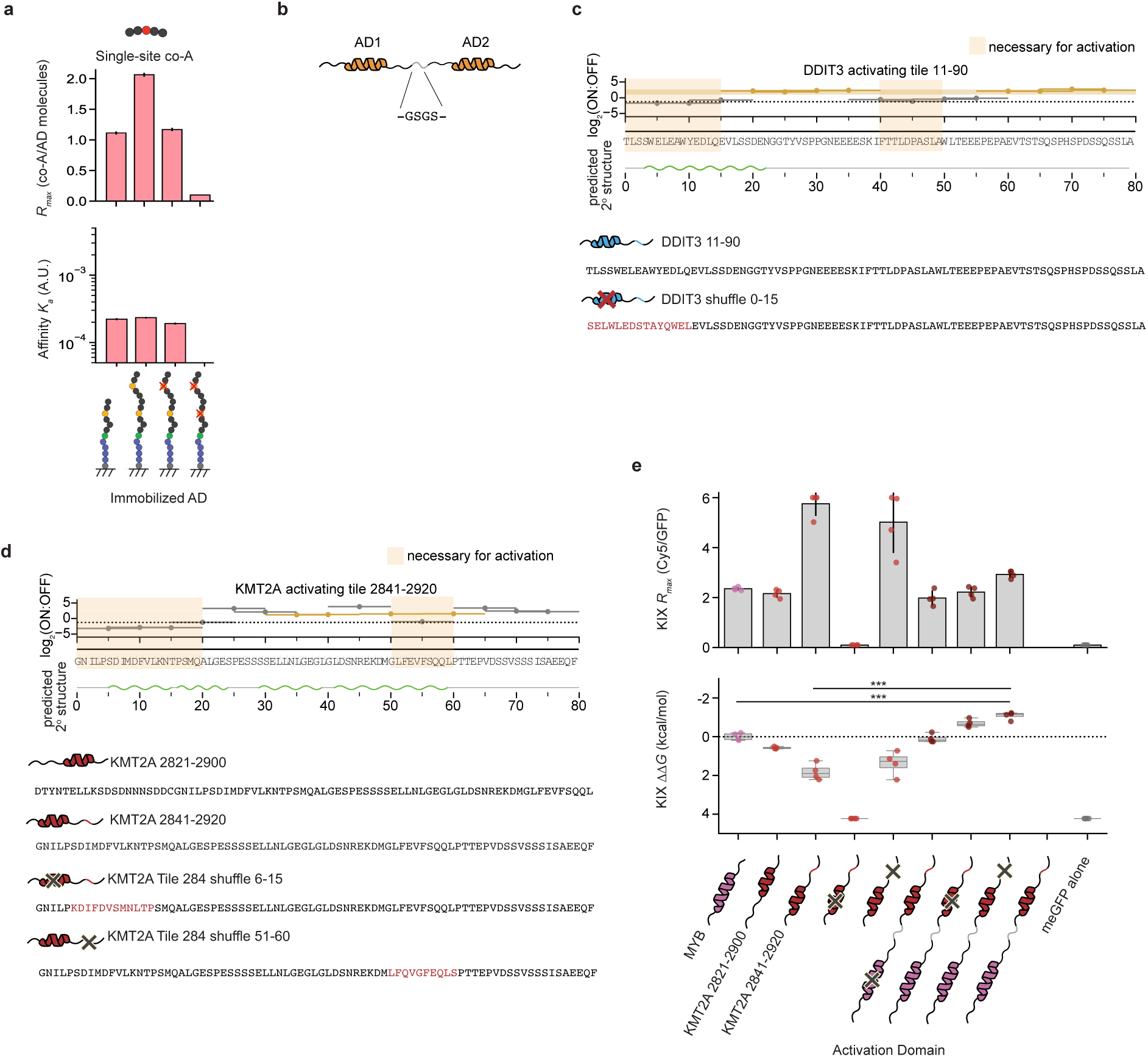
**Related to Figure 3 a**, Predicted affinity (*K_a_*) and occupancy (*R_max_*) from PIMMS simulations for a co-A with a single site interacting with surface-immobilized ADs containing one or two cognate (yellow) or mutated (red) binding sites. **b**, Tandem synthetic ADs measured within STAMMPPING assays (Fig. 3e) were linked by a (GS)_2_ linker, with lengths between neighboring activation-necessary regions estimated by ALBATROSS^75^ (**Supplementary Table 13, Supplementary Note 3**). **c**, Measured fraction of cells activated for DDIT3 constructs containing tiled 10 aa deletions to identify activation-necessary regions^20^ and full sequences of WT and shuffled mutants profiled via STAMMPPING. **d**, Measured fraction of cells activated for KMT2A constructs containing tiled 10 aa deletions to identify activation-necessary regions^20^ and full sequences of WT and shuffled mutants profiled via STAMMPPING. **e**, Measured relative affinity (*ΔΔG*) and occupancy (*R_max_*) for the P300 KIX domain binding synthetic multi-site ADs based on KMT2A and MYB. Bars indicate mean, vertical lines represent standard deviation; n.s.: p>0.05, *:p<0.05, **:p<0.001 (two-sided t-test). No data were collected for MYB-GS-KMT2A due to failed expression on-chip.

**Extended Data Figure 8.**
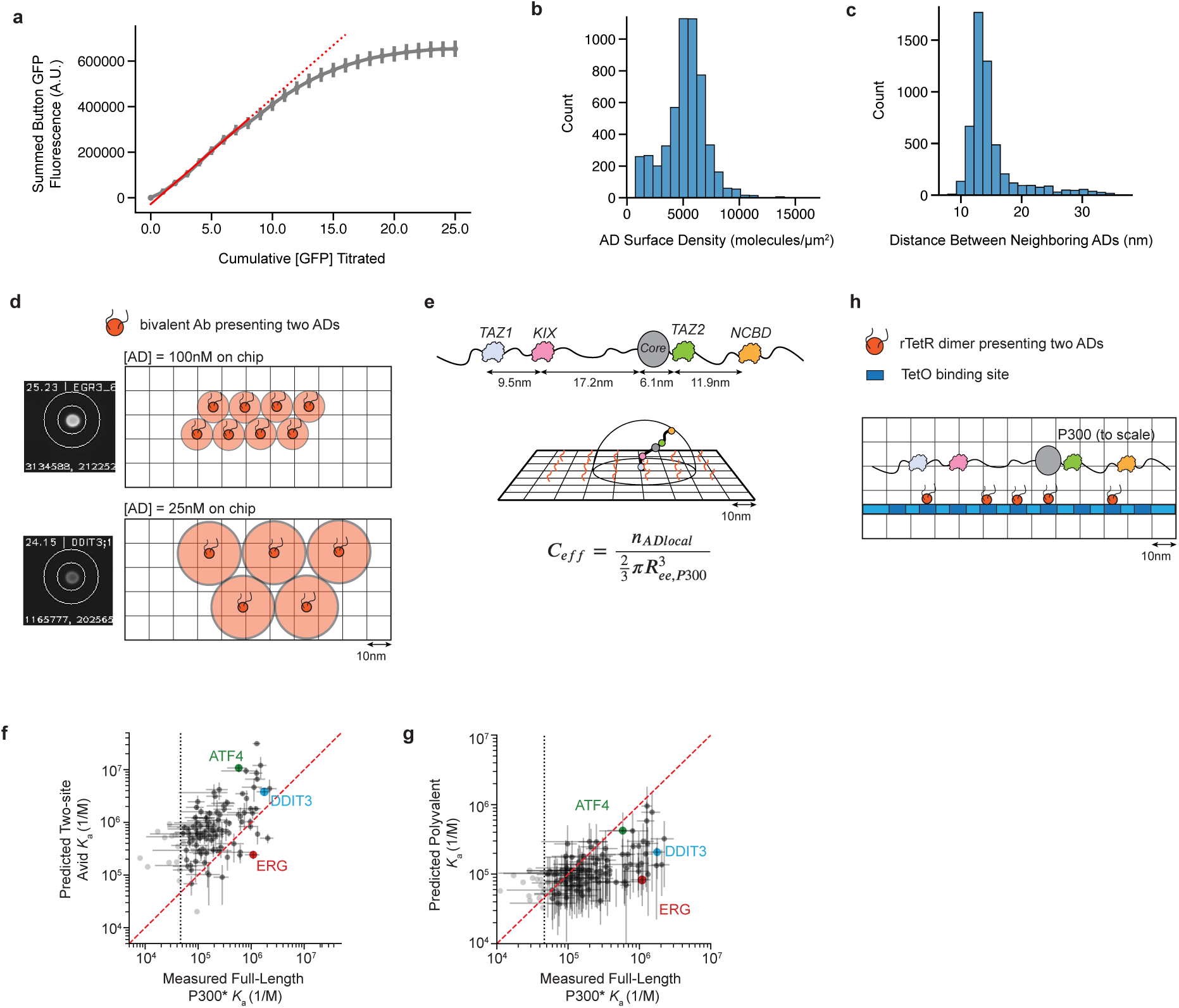
**Related to Figure 3 a**, Standard curve relating measured microfluidic well spot fluorescence to cumulative titrated GFP concentration; red dashed line indicates a linear fit to the first eight data points (y=mx) with a best fit slope of 43,000 a.u./nM GFP. **b-c**, Distributions of AD surface density and computed distance between neighboring ADs for chambers across a standard STAMMPPING chip (data shown for full-length P300 binding experiments) (**Methods**). **d**, Scale diagrams of AD density as seen from above on patterned surfaces *in vitro*. Light-orange circles represent average distance between bivalent antibodies each presenting two ADs (12 nm for 100 nM case, 24 nm for 25 nM case). **e**, Scale diagram of P300 and estimation of *C_eff_* for prediction of affinity in an avid binding scenario on-chip. P300 IDR lengths estimated by ALBATROSS^75^. **f**, Predicted affinity from “two-site avid” model *vs.* measured affinity for full-length P300/AD binding interactions; “Two-site avid” prediction calculated by *K_a,avid_2site_* = *K_a1_ K_a2_ C_eff_*, where *K_a1_* and *K_a2_* are affinities of the two tightest-binding P300 subunits for a given AD and *C_eff_* estimated as shown in panel **e**. P300’s *K*_a_s are marked with an asterisk to indicate the presence of punctae that may reduce the effective concentration of soluble protein such that the *K*_a_s represent a lower limit (**Supplementary Note 1**). **g**, Predicted affinity from polyvalent model *vs.* measured affinity for full-length P300/AD binding interactions. Dashed line indicates average meGFP negative control; red dashed line indicates the identity line. **h**, Scale diagram of P300 aside the synthetic nine TetO array used as a proximal promoter in this study. For illustration, five rTetR dimers are shown simultaneously bound, based on ref:^76^.

**Extended Data Figure 9.**
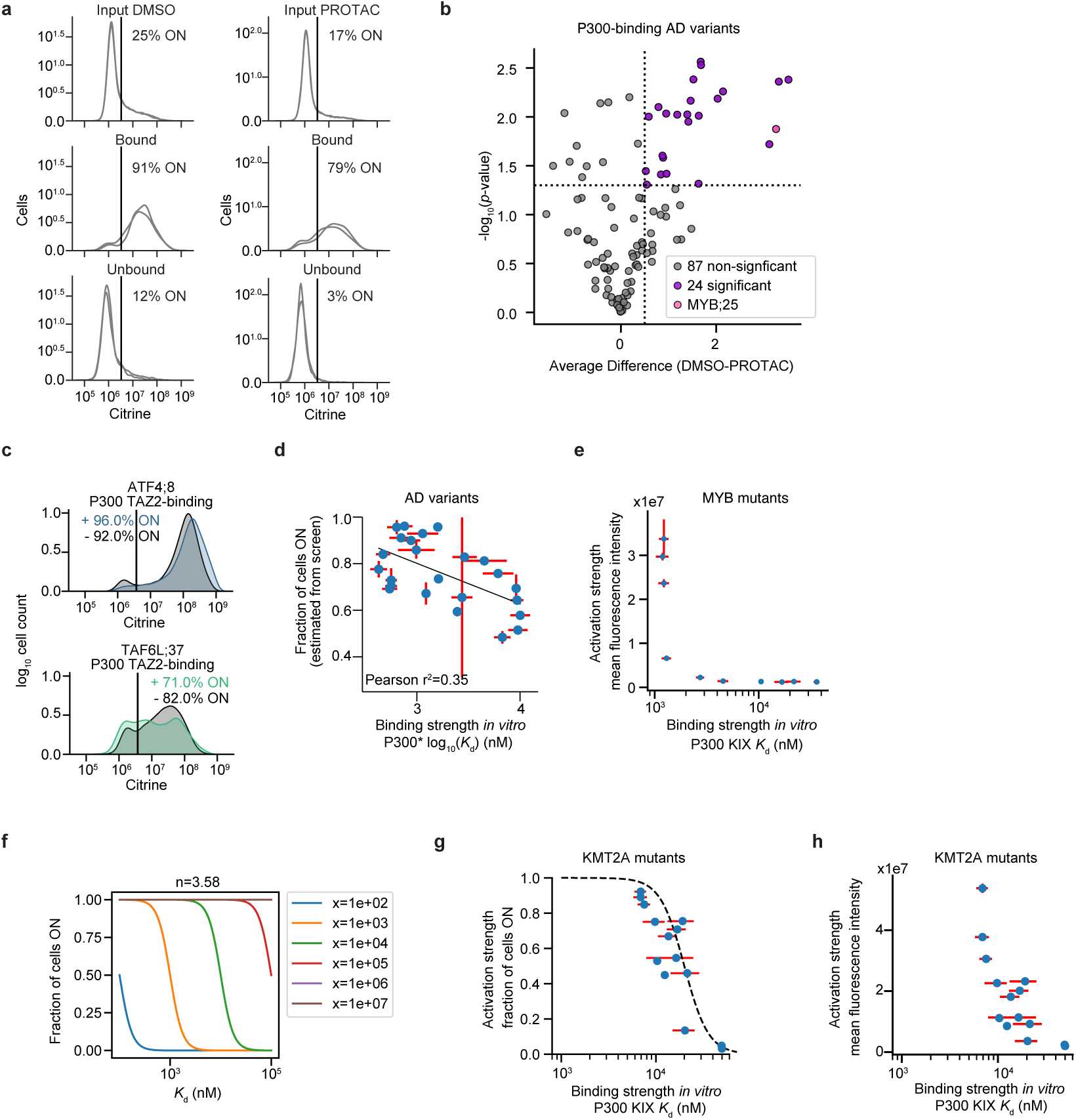
**Related to Figure 4 a**, Distributions of single-cell citrine reporter distributions measured via flow cytometry for the PROTAC HT-recruit screen on the day of magnetic separation (Input) and after separation (Bound, Unbound). Overlapping histograms are shown for two separately transduced biological replicates. The average percentage of cells ON is shown to the right of the vertical line indicating the citrine level gate. **b**, Volcano plot comparing log_10_-transformed *p*-values (calculated from a t-test comparing the distribution of enrichment scores between vehicle and drug conditions) *vs.* average difference in activation enrichment scores for all P300-binding ADs; activating sequences with *p*≤0.05 and average difference>0.5 (dashed lines) were considered significant (purple points). **c**, Recruitment to minCMV promoter in the absence (black) or presence (colors) of 500 nM of dCBP-1. **d**, Comparison between HT-recruit activation scores that were estimated from screen enrichment scores to fraction ON using a calibration curve based on flow cytometry individual measurements (**Methods**) from ref:^20^ and STAMMPPING-measured affinities for PROTAC-sensitive P300-binding ADs/P300. P300’s *K*_d_s are marked with an asterisk to indicate the presence of punctae that may reduce the effective concentration of soluble protein such that the *K*_d_s represent an upper limit (**Supplementary Note 1**). **e**, Comparison between mean citrine fluorescence intensities from cells containing individually recruited MYB mutants and corresponding STAMMPPING-measured affinities for P300’s KIX domain. **f**, Simulation showing that the cooperativity equation shown in Fig. 4j has the same general shape for different AD concentrations, here plotted as x shown in different colors. **g**, Comparison between fraction of cells ON from individually recruited KMT2A mutants and corresponding STAMMPPING-measured affinities for P300’s KIX domain. The same MYB mutant data fit-hill coefficient of 3.58 was used when plotting the cooperativity equation (dashed line). **h**, Comparison between mean fluorescence intensities from individually recruited KMT2A mutants and corresponding STAMMPPING-measured affinities for P300’s KIX domain.

## Acknowledgements

We thank members of our laboratories for helpful conversations and assistance; Renee Hastings for helpful suggestions to the writing; Micah Olivas for assistance in running AlphaFold predictions on a local server; Michael Hayes, Matt DeJong, Albert Lee, Eli Costa, and Michaela Hinks for conversations related to models of binding and activation. This work was supported by NSF CAREER Award 2142336 (PMF); National Cancer Institute NIH-NCI DP2 CA290639-01 (ASH); NIH MIRA R35GM128947 (LB); NSF Graduate Research Fellowship DGE-1656518 (ND, MUS); NSF Graduate Research Fellowship DGE-2139839 (DG, JML); ARCS Foundation Fellowship (ND); Stanford Bio-X Bowes Fellowship (PHS); Agilent Fellowship (PHS); MIT Career Advising & Professional Development’s Career Exploration Fellowship (SK); Graduate Fellowship Award from Knight-Hennessy Scholars at Stanford University (MUS); PMF is also a Chan Zuckerberg Biohub San Francisco Investigator.

## Authorship Contributions

ND and PHS performed and analyzed STAMMPPING experiments. PHS designed and tested microfluidic molds and devices. ND and PHS fabricated microfluidic molds and devices. ND performed microscale thermophoresis measurements. ND and SK performed *in cellulo* experiments. PHS derived multivalency models. DG, JL, and BN designed and implemented *in silico* experiments with supervision from ASH. MUS analyzed STAMMPPING AD expression data. ND, PHS, DG, JL, BN, ASH, LB, PMF conceived of the study. ND wrote the manuscript with significant contributions from PHS, PMF, and input from all authors. PMF and LB supervised the project.

## Supplementary Figures

**Supplementary Figure 1: Microfluidic device designs**

Detailed schematic of the microfluidic devices used for STAMMPPING measurements. “Control” (orange) and “flow” (blue) layers with common and block-specific input ports are labeled.

**Supplementary Figure 2: STAMMPPING concentration-dependent binding measurements for 204-member activation domain library binding 13 co-activators and co-activator domains**

**a,** Molecular weights of tagged co-activators tested in this experiment. **b**, GFP expression levels of AD variants (blue) and empty chambers (orange) across the device. **c**, Denaturing gel and quantification (right) comparing purified AlexaFluor647-labeled SNAP protein masses to IVTT-expressed AD masses. **d**, Binding curves with Langmuir isotherm fit overlaid and average *K*_d_ shown grouped by AD where negative controls are shown last. **e**, Pairwise comparisons between replicate *K*_a_ values where gray indicates non-binding ADs and blue indicates binding ADs (where both replicates are greater than or equal to one standard deviation above the median meGFP measurements). Values lower than one standard deviation below the median meGFP measurements were set to this value. Red marker indicates meGFP measurements; at least one meGFP measurement always passed quality control checks, but if only one meGFP measurement is present, no red marker will be plotted on joint scatter plots. **f**, CY5 and GFP images with quantification overlaid for a subset of binding and non-binding ADs.

**Supplementary Figure 3: MYB mutant/P300-KIX concentration-dependent binding curves**

Concentration-dependent binding data with Langmuir isotherm fit overlaid and average *K*_d_ shown; data corresponds to Supplementary Table 5.

**Supplementary Figure 4: TAZ2RR1732AA concentration-dependent binding curves**

Concentration-dependent binding data with Langmuir isotherm fit overlaid and average *K*_d_ shown; data corresponds to Supplementary Table 6.

**Supplementary Figure 5: P300 KIX and P300 TAZ2 concentration-dependent binding curves across varying salt concentrations**

Concentration-dependent binding data with Langmuir isotherm fit overlaid and average *K*_d_ shown; data corresponds to Supplementary Table 7.

**Supplementary Figure 6: P300 TAZ2 concentration-dependent binding curves for ATF4;8, FOXO1;56 and TAF6L;37 mutant activation domains**

Concentration-dependent binding data with Langmuir isotherm fit overlaid and average *K*_d_ shown; data corresponds to Supplementary Table 8.

**Supplementary Figure 7: P300 TAZ2 concentration-dependent binding curves for ATF4;8 intrinsically disordered region mutants**

Concentration-dependent binding data with Langmuir isotherm fit overlaid and average *K*_d_ shown; data corresponds to Supplementary Table 9.

**Supplementary Figure 8: P300 KIX concentration-dependent binding curves for synthetic multivalent activation domains**

Concentration-dependent binding data with Langmuir isotherm fit overlaid and average *K*_d_ shown; data corresponds to Supplementary Table 10.

**Supplementary Figure 9: MYB mutant individual reporter gene activities**

Flow cytometry intensity distributions of reporter gene expression levels after 48 hours of recruitment of MYB mutants (red, gray indicates no recruitment).

**Supplementary Figure 10: P300 KIX concentration-dependent binding curves for KMT2A mutant activation domains**

Concentration-dependent binding data with Langmuir isotherm fit overlaid and average *K*_d_ shown.

**Supplementary Figure 11: KMT2A mutant individual reporter gene activities**

Flow cytometry intensity distributions of reporter gene expression levels after 48 hours of recruitment of KMT2A mutants (red, gray indicates no recruitment).

## Methods

### Fabrication of microfluidic molds and devices

The STAMMPPING device was designed in AutoCAD 2020 (Autodesk). The following changes were made to the MITOMI device^44,113,114^: four parallel flow blocks with two inlets and one outlet per block were created to simultaneously measure four prey proteins; the chamber array was extended to a 32x56 array containing 1,792 total chambers (a 40% increase over previous generation chips); and neck valves were widened and their shape was altered to maximize valve surface area and minimize opportunity for valve crosstalk defects (**Supplementary Figure 1**). Masks for control layers were enlarged 1.5% relative to the flow layer to account for features shrinking after PDMS control layer bakes. Flow and control molds were fabricated on 4” silicon wafers (University Wafer) using transparency masks printed at 25,000 DPI (FineLine Imaging).

Flow molds were composed of the following layers: (1) a ∼5 µm uniform layer of SU-8 2005 (Microchem) covering the entire wafer surface to improve feature adhesion (soft bake at 65°C 2 min, 95°C 3 min, 65°C 2 min; UV flood exposure 20 sec; post-exposure bake at 65°C 2 min, 95°C 4 min, 65°C 2 min) (2) a ∼15 µm layer of AZ50 XT (Capitol Scientific) (let sit for 10 min; soft bake from 50-112°C, 450°C/hr, 18 min; rehydrate overnight in box with water; UV mask exposure 3x20 sec; developed in AZ400k) subjected to a slow final hard bake to generate rounded valve features (from 65-190°C, 10°C/hr, 14 hrs), and (3) a ∼15 µm layer of SU-8 2015 (Microchem) to generate square flow channels (let sit for 5 min; soft bake at 65°C 2 min, 95°C 4 min, 65°C 2 min; UV mask exposure 22 sec; post-exposure bake at 65°C 3 min, 95°C 4 min, 65°C 3 min, developed in SU-8, hard bake from 65-165°C, 120°C/hr, 2 hrs). Control molds were composed of the following layers: (1) a ∼5 µm uniform layer of SU-8 2005 (Microchem) covering the entire wafer surface to improve feature adhesion (same bake times as above), (2) a single ∼25 µm layer of SU-8 2025 (Microchem) (soft bake at 65°C 2 min, 95°C 7 min, 65°C 2 min; UV mask exposure 25 sec; post-exposure bake at 65°C 3 min, 95°C 8 min, 65°C 3 min, developed in SU-8, hard bake from 65-165°C, 120°C/hr, 2 hrs).

PDMS devices were fabricated from molding masters as described in^114^, with the following modification: PDMS was spin coated onto the flow mold with modified parameters to account for taller features on newer molds (10 s ramp at 500 rpm, 133x acceleration; 75 s spin coating at 1775 - 1825 rpm, 266x acceleration).

### Preparation of linear DNA template libraries for on-chip expression

#### Library synthesis and initial amplification

Putative activation domain sequences (**Supplementary Table 1**) were ordered as two 300 nucleotide sub-pools (Twist): activation domains from chromatin regulators (CR sub-library) and activation domains from transcription factors (TF sub-library). The pools were resuspended in EB to a final concentration of 10 ng/µL. 6x 50 µL PCR reactions per library were set up as follows (1x): 0.5 µL of template, 10 µL 5x Herc buffer, 1 µL dNTPs (10 mM), 1 µL each of forward and reverse primers (10 µM), 1 µL DMSO, 1 µL Herc polymerase, 34.5 µL water. Reactions underwent denaturation for 3 minutes at 98°C, then cycled through (1) 98°C 20 sec, (2) 61°C 20 sec, (3) 72°C 30 sec for 21 cycles (TF sub-pool) or 23 cycles (CR sub-pool), and a final elongation at 72°C for 3 min. The PCR reactions were combined and gel-purified (Qiagen). Golden gate reactions of pre-cut meGFP backbone (NVD006, to be deposited on Addgene) and each subpool were set up as follows: 75 ng of backbone, 5 ng of insert, 2 µL of 10x T4 DNA ligation buffer, 1 µL of Golden Gate Assembly Kit BsmBI-v2 (NEB), and water to 20 µL. Reactions were thermocycled between 42°C 5 min and 16°C 5 min for 65 cycles with a final incubations of 42°C for 5 min and 70°C for 20 min. Reactions were pooled and concentrated (MinElute, Qiagen).

#### Bacterial transformation and colony picking

Each PCR-purified pool was transformed into bacteria by combining 2 µL of the pool with 25 µL of electrocompetent cells (Lucigen). The DNA/cell mixture was added to a chilled cuvette and electroporated using a Gene Pulser Xcell (BioRad) with the following settings: 1.8 kV, 600 Ohm, 10 µF. Immediately afterwards, 2 mL of recovery medium was added to the cuvettes and transferred to a 14 mL round bottom tube. Tubes were shaken at 250 RPM in a 37°C incubator for 60 min. A 1:500 dilution of the culture was plated and spread evenly on 10“x10” LB-agar plates with 100 µg/mL Carbenicillin. Plates were wrapped with plastic bags to prevent them from drying out and incubated in a warm room for 15 hours. Single colonies were picked from source LB agar plates into 384 well (120 μL well volume) destination microwell plates containing 60 μL LB with ampicillin. We manually picked a total of 3,840 colonies for the 1,155 variant library. Microwell plates were sealed with gas-permeable AeraSeal film (MilliporeSigma) and grown to saturation (15 hours) with shaking (900 RPM) at 37°C. Master bacterial plate cultures were diluted 1:10 dilution into water (2 µL into 18 µL) using a 96 pinhead Rainin Liquidator pipette (Mettler Toledo).

#### Colony PCR and well-specific barcoding

Colony PCR reactions were then performed as follows: 2 µL of KAPA HiFi HotStart ReadyMix (Roche) was assembled with 0.8 µL of 1:10 bacteria dilution, 0.12 µL of forward and reverse primers (100 µM), and water up to 4 µL. Reactions underwent denaturation for 5 minutes at 95°C, cycled through (1) 98°C 20 sec, (2) 60°C 15 sec, (3) 72°C 2 min for 25 cycles, and underwent a final elongation at 72°C for 2 min. Completed PCR reactions were serially diluted into water 1:10 then 1:100. Libraries were then barcoded and amplified using KAPA HiFi polymerase and a collection of Nextera XT 12mer dual unique index sequencing primers (purchased from IDT and supplied by CZ-Biohub). First, 1.2 µL of a master mix containing 0.08 µL of KAPA HiFi, 0.8 µL of 5x buffer (Roche), 0.12 µL dNTPs (10 mM) and water was added to 1 µL of the 1:100 diluted colony PCR reactions using the Mantis liquid handler. Next, 0.8 μL of unique i5/i7 primer mix (2.5 μM each) was transferred from source plates to each sample using the Mosquito instrument. Reaction plates were sealed, briefly vortexed, collected by centrifugation, and amplified with the following thermal cycler conditions: 95°C, 30 sec; 18 × [98°C, 10 s; 55°C, 15 s; 72°C, 1 min]; 72°C, 5 min. Resulting libraries were pooled (with the Mosquito) and purified using AMPure XP magnetic beads at a ratio of 0.8:1 beads:sample volume (Beckman Coulter, Brea, CA, USA). Library yield and quality were determined by TapeStation electrophoresis with an HSD1000 ScreenTape system (Agilent Technologies).

#### Next-generation sequencing (NGS) and analysis

Sequencing was performed by SeqMatic (Fremont, CA, USA). Libraries were sequenced using MiSeq Micro 2x150bp with the addition of 20% PhiX control DNA; samples were submitted with i5/i7 barcodes corresponding to each variant and demultiplexed by the instrument. NGS reads were aligned to the oligonucleotide index using Bowtie^115^ allowing for 0 mismatches, read length of 100 and trim length of 30. Wells below 10 counts were filtered out. Only wells that had the same variant identified via read 1 and 2 and had single variants identified per well were kept. This resulted in 805 unique variants, which were then cherry-picked from the 1:10 dilution of the original master bacterial plate into new plates of only single variant bacteria wells using the Biomek instrument (Beckman Coulter). 743 of these 805 variants activated in a validation screen^20^. Only variants that activated in this validation screen were chosen for co-A binding experiments.

#### Generation of final templates for in vitro transcription/translation

To add the components necessary for in vitro transcription and translation to DNA templates (T7 promoter, ribosome binding site, T7 terminator), PCR reactions were performed as follows: 4 µL of KAPA HiFi HiFi HotStart ReadyMix (Roche) was assembled with 1.6 µL of 1:10 bacteria dilution, 0.24 µL of forward and reverse primers (100 µM), and water up to 8 µL. Reactions underwent denaturation for 5 minutes at 95°C, then cycled through (1) 98°C 20 sec, (2) 60°C 15 sec, (3) 72°C 2 min for 25 cycles, and underwent a final elongation at 72°C for 2 min. 1 µL of the completed PCR reactions was then combined with 4 µL of water and 5 µL of print buffer (2x: 20 mg/mL UltraPure BSA (Thermo Fisher), 24 mg/mL trehalose dihydrate, 100 mM NaCl) prior to printing on glass slides.

### Preparation of plasmid libraries for on-chip expression

Smaller libraries (<100 variants) were prepared as plasmids. Activation domain sequences were ordered as eBlock Gene Fragments (IDT). These fragments were then cloned into a PurExpress meGFP backbone (NVD006, to be deposited on Addgene) by mixing 75 ng of backbone, 5 ng of insert, 1 µL of 10x T4 DNA ligation buffer, 1 µL of Golden Gate Assembly Kit BsmBI-v2 (NEB), and water to 10 µL. Reactions thermocycled between 37°C 5 min and 17°C 5 min for 15 cycles and then underwent a final incubation at 42°C for 5 min and 70°C for 20 min. Reactions were transformed in Stbl3 Chemically-Competent E. coli, plated in LB plates containing 100 µg/ul Carbenicillin, inoculated into liquid culture, miniprepped (Qiagen), and eluted in UltraPure distilled water. Sequences were validated by Sanger sequencing (Quintara Bio).

### Preparation of epoxysilane-coated slides

Plain 25x75mm glass slides (Corning) were functionalized with epoxysilane (based on ref:^116^). First, slides were cleaned in an 80°C heated bath of hydrogen peroxide and ammonium hydroxide for 30 min, rinsed, and blow-dried with nitrogen. Next, epoxysilane was deposited by submerging slides in toluene doped with 3-glycidyloxypropyl-trimethoxymethylsilane (3-GPS, Millipore-Sigma) for 20 min. Slides were then rinsed in toluene and dried by baking at 120°C for 30 minutes.

### Printing DNA libraries onto functionalized slides

For all on-chip experiments, DNA templates (plasmids or linear templates) encoding AD-meGFP constructs were spot-printed on epoxysilane-functionalized slides in an array with pitch dimensions matching the distance between chambers of our PDMS devices. Custom Python scripts were used to uniquely assign mutants in 384-well plates to be deposited in specific locations within spotted arrays. Typical print maps included 4-8 replicates per template per block and empty chambers every 8 rows to sensitively detect any cross-talk between chambers. Prior to printing, DNA templates were resuspended in a “print buffer” of 10 mg/ml UltraPure BSA (Thermo Fisher), 12 mg/ml trehalose dihydrate, and 50 mM NaCl in UltraPure distilled water. Each template was then printed as 250-350 pL droplets across 15-20 replicate epoxysilane-coated slides using a sciFLEXARRAYER S3 fitted with PDC60 or PDC70 nozzles with Type 1 coating (Scienion).

### Preparation of IVTT-expressed proteins

First, co-As were cloned into a PurExpress backbone that either contains an N-terminal 3x FLAG tag and C-terminal SNAP tag (NVD103, to be deposited on Addgene) or only a C-terminal SNAP tag (NVD005, to be deposited on Addgene). PurExpress parts A and B were incubated for either 45 minutes or overnight at 4°C before combining with at least 100 ng of template, 1 µL of RNase Inhibitor (NEB), 1 µL of 25x cOmplete protease inhibitor solution (Roche, Millipore Sigma), and water to 25 µL. Reactions were incubated at either 30 or 37°C for at least 2 hours or up to overnight. Proteins were then labeled with Alexa647 by combining 2 µL of 250 µM SNAP-AF647 label with 23 µL of PBS and incubated at 37°C for 30 minutes. Several reactions were then pooled and spun-concentrated with Amicon ultra centrifugal filters 10 kDa MWCO (Sigma-Aldrich). We used 50 µL of Thermo Scientific Pierce Anti-DYKDDDDK Magnetic Agarose and a binding buffer composed of 20 mM KH_2_PO_4_, 200 mM KCl, 0.05% (v/v) Tween-20 to purify FLAG-containing proteins. We incubated the proteins and beads for 4 hours at 4°C with end-over-end mixing. Bead-bound FLAG protein samples were washed twice with binding buffer and eluted twice with 75 µL of 3 mg/mL 3xFLAG peptide (Sigma-Aldrich) before the eluate was spun-concentrated once more. For each co-A, we quantified concentrations via comparison of gel electrophoresis band intensities for Alexa647-conjugated SNAP-tagged proteins and an in-gel concentration standard (**Supplementary Figure 2**).

For commercially purchased FLAG-tagged co-As (full-length P300 (Active Motif, 81158) and BRD4’s BRD domain (Active Motif, 81155)), we measured binding via co-introduction of fluorescent anti-FLAG and performed additional negative control experiments quantifying binding between the 208-member library and the anti-FLAG antibody incubated with equimolar amounts of 3xFLAG peptide (Sigma-Aldrich) as a negative control (**Supplementary Figure 2**).

### Preparation of purified proteins from bacterial cultures

First, co-As were cloned into a Pet24a backbone that contained an N-terminal 6x-His tag along with a C-terminal SNAP tag. Plasmids were transformed into BL21(DE3) competent *E. Coli* cells (NEB). Single colonies were inoculated first into small 5 mL cultures of 2x YT medium with 50 µg/mL Kanamycin. After shaking for 5 hours at 37°C, the cultures were transferred to large 1 L cultures, and incubated again at 37°C until the optical density reached 0.6. Once the cultures were in the log growth phase, the flasks were chilled on ice for 3 minutes before 0.5 mM IPTG was added to induce expression. The cultures were then incubated at 16°C overnight. The cultures were then spun down at 4,200 x g for 30 minutes, resuspended in 50 mM Tris buffer, and left in the -80. The next day, the frozen pellet was thawed. Protease inhibitor was added (600 µL of 50 mM PMSF) and left at room temperature for 10 minutes before 4.2 mL of 4M NaCl was added. Cells were lysed using a sonicator for a total of 5 minutes (pulse 30 seconds on 59 seconds off, amplitude 20%). Lysed cells were then centrifuged at 12,000 x g for 30 minutes. Imidazole was then added to the supernatant to reach a final concentration of 40 mM. A nickel chloride gravity column was first equilibrated with wash buffer (50 mM HEPES, 500 mM NaCl, 10% w/v glycerol, 5 mM DTT, 40 mM Imidazole). Next, the lysate was introduced into the column and washed once with wash buffer. 1 mL of elution buffer (50 mM HEPES, 500 mM NaCl, 10% w/v glycerol, 1 mM DTT, 500 mM Imidazole) was added to the column and each fraction was collected and labeled. Expression was confirmed with SDS-PAGE electrophoresis before combining fractions and spin-concentrating down to 500 µL-1 mL with Amicon ultra centrifugal filters 10 kDa MWCO (Sigma-Aldrich). The protein was then SNAP labeled in 2x molar excess of SNAP-Surface AlexaFluor 647 (NEB) and incubated at 37°C for 1 hour. Next, the labeled protein was filtered with a 0.22 µm syringe filter and purified using a Superdex 75 10/300 (GE) size exclusion column loaded on an AKTA liquid chromatography system. Purification was confirmed with SDS-PAGE electrophoresis and pure fractions were combined and concentrated. Protein concentration was estimated from predicted extinction coefficients and triplicate 280 nm absorbance measurements.

### Microscopy instrumentation

Chip imaging was performed using a Nikon Ti-2 Microscope equipped with motorized XY-stage (Applied Scientific Instruments, MS-2000 XY stage), LED light source (Lumencor SOLA SE Light Engine), 4x Objective (CFI Plan Apo 4X NA 0.20, Nikon), and CMOS camera (Andor Zyla 4.2). The Nikon Ti-2 core itself was equipped with an automated Z-axis, turret aperture, and filter turret, containing pre-arranged filter cubes for EGFP (Semrock, GFP-4050B), Cy5 (Semrock, Cy5-5070A). All imaging was performed using 2x2 pixel binning with exposure times as follows: EGFP: 100 ms and 500 ms, Cy5: 50 ms, 100 ms, 500 ms, 1000 ms, and 3000 ms.

### STAMMPPING device operation and experimental pipeline

Microfluidic valves were operated via a custom-built open-source WAGO-controlled pneumatic manifold^117^. Automated control of pneumatic valves and microscopy was run through custom Python scripts (RunPack, AcqPack) as described previously^114^; here, we used slightly updated scripts to manage novel “block” valves. Briefly, standard STAMMPPING experiments consist of the following steps: (1) pattern button-valve surfaces for AD immobilization, (2) express AD-meGFP constructs from DNA templates on chip, and (3) iteratively introduce labeled co-A protein, allow interactions to come to equilibrium, trap bound material using ‘button’ valves, wash away unbound material, and image in both the eGFP and Cy5 channels to measure binding. Steps 1 and 2 were carried out largely as described previously^44,114^, with the following modifications. For surface patterning, we used a biotinylated monoclonal anti-GFP antibody (Thermo, GFP Monoclonal 9F9.F9). We also optimized on-chip expression for disordered AD-GFP constructs such that expression conditions varied from 30-37°C for 2-12 hrs. Following expression and binding of AD-GFP to anti-GFP-patterned buttons, device chambers were washed in TrypLE and regions surrounding the protected buttons were re-patterned with biotinylated BSA. For step 3, dye-labeled proteins made off-chip were prepared as serial dilutions in an Assay Buffer of 20 mM KH_2_PO_4_, 200 mM KCl, 5 mg/ml UltraPure BSA (Thermo Fisher), 0.05% v/v Tween-20 (Thermo Fisher) in UltraPure distilled water. On-chip iterative measurements of binding over multiple concentrations were carried out via scripts based on those used previously^44^, with the modification that proteins were incubated for 1 hour to reach equilibrium and at each step in a concentration series and proteins were added to the block inlets (A1, B1, etc., **Supplementary Figure 1**) to simultaneously test 4 or 8 co-As per chip without any chance of crosstalk or contamination.

### STAMMPPING image processing

Raw 7x7 image rasters of fluorescence images covering the full chip area were flat-field corrected using manual flat-field images or an implementation of BaSiCpy^118^ and stitched into single full-chip images using ImageStitcher, an in-house publicly-available software package (https://github.com/FordyceLab/ImageStitcher). These stitched images were processed using ProcessingPack, an in-house publicly-available software package (https://github.com/FordyceLab/ProcessingPack). Briefly, circle-finding algorithms identified fluorescent spots under chamber “button” valves and per-spot metrics were mapped to Scienion print array position. Downstream analysis primarily utilized background-subtracted summed intensities of GFP and Cy5 per-spot fluorescence.

### STAMMPPING quality control

Prior to fitting curves to determine binding affinities, we performed several quality control steps. First, we filtered out any chambers with GFP expression levels above the threshold set by considering blank chambers (mean + one or more standard deviations) (**Supplementary Figure 2**). Second, we manually culled chambers that had visible dust or flow issues. Third, we computed a linear regression between measured Cy5 intensities of soluble co-A in each chamber and the estimated input concentration and required r^2^ >0.9 (ensuring that soluble co-A was available for binding as expected). Fourth, we removed any chambers that had Cy5/GFP ratios below 0 across all concentrations.

### Determination of equilibrium binding constants

All *K_d_*s were estimated from globally fitting a Langmuir isotherm (**Eqn. 1**) to concentration-dependent binding data.

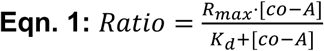

For initial AD/co-A activator measurements, we used global fits assuming that all AD/co-A interactions bind with the same stoichiometry to accurately determine *K*_d_s even for interactions that did not saturate at the highest concentrations tested. First, we estimated the saturation point for a given co-A experiment by fitting the top binders (binders with a Cy5/GFP Ratio above a set threshold set to identify most-saturated binders) to **Eqn. 1** with *K_d_* and *R_max_* as free parameters. The mean *R_max,top_binders_* was then used as a global *R_max_* to re-fit all remaining AD-co-A interactions with *K_d_* as a free parameter. From these fits, we calculated the median *K_d_* value across replicates for each AD/co-A pair.

The assumption that a given co-A will bind all ADs with a given stoichiometry likely holds for most ADs binding individual co-A subunits. Fits interrogating synthetic multivalent ADs with the potential to vary significantly in *R_max_* between samples (**Fig. 3e**, **Extended Data Fig. 7e**) were conducted without the global *R_max_* estimate described above such that fits were conducted with free *K_d_* and *R_max_*.

All data fitting was done in custom Python scripts using open-source lmfit and scipy packages^119,120^.

### Determination of noise floor and classifying binding

The experimental noise threshold for each co-A was set to each co-A’s median meGFP *K*_d_ measurement, with the exception of TFIID’s TAF12, where the meGFP measurement was unreliable (>100 µM *K*_d_); for this co-A, we used Random control protein 778’s binding affinity for TAF12. Next, we set all returned fitted *K*_d_ values weaker than this threshold to the meGFP *K*_d_ and denoted the value as a lower limit. ΔΔG values are calculated by subtracting each meGFP/co-A’s ΔG (ΔG=-RTln(*K*_d_)) from each AD/co-A interaction’s ΔG (for TFIID’s TAF12 subunit, we again used Random control protein 778).

Affinities were binarized as binding or non-binding (**Fig. 1j**) by comparing the distribution of affinities between AD/co-A and negative control/co-A interactions (meGFP, Random control 810, Random control 778). For each co-A, we removed negative control outliers (>1.5 times the interquartile range (IQR)). AD/co-A pairs were labeled as binding if their affinities were above the noise floor and significantly different (one-sided t-test) from the negative controls.

Top binders (**Fig. 1i**) were classified from each co-A’s *K*_d_ distribution’s IQR. Specific co-As have more narrow IQRs, whereas promiscuous co-As have wider IQRs; therefore, we used two different metrics for these two groups. For promiscuous co-As (P300 TAZ2 and P300), we classified ADs with *K*_d_s equal to or below the 10th percentile as top binding. For specific co-As (P300 KIX, P300 NCBD, P300 TAZ1, BRD7, TFIID TAF12 and TAF6L), we fit a gaussian to each of these distributions and classified ADs with *K*_d_s that were 3 sigma (P300 KIX, P300 NCBD, P300 TAZ1), 4 sigma (BRD7, TAF12), or 6 sigma (TAF6L) below the mean as top binders.

### Determination of on-chip AD density via GFP titration

GFP titration experiments were performed in order to create a calibration curve between GFP fluorescence intensity and protein concentration on-chip^114^. First, we patterned STAMMPPING devices with antiGFP pedestals as described above. Next, we prepared a solution of 1 nM purified eGFP (OriGene Technologies) containing 2% UltraPure BSA (Thermo) (to minimize non-specific adsorption). We then introduced this 1 nM GFP onto the chip in an iterative cycle with the following steps: (1) flow on GFP with buttons closed to fill chambers, (2) close sandwich valves to trap one chamber volume of GFP and open buttons to allow for binding, (3) incubate for 30 minutes, and (4) image GFP intensity of buttons and chamber background. Iterations were repeated until buttons were saturated with GFP, approximately 25 cycles. We generated a standard curve by fitting the initial slope relating raw fluorescence intensity (A.U.) to effective protein concentration within the chamber volume (nM). We estimated AD surface density by converting the effective concentration within the chamber volume to a 2D button surface density (*ρ_AD_*) using the formula *ρ_AD_ =* [AD] * *V_chamber_ / A_button_*, where *V_chamber_* is the volume of a single microfluidic chamber and *A_button_* is the 2D area of the button spot on which ADs are immobilized. *V_chamber_* was estimated by multiplying the flow channel height of 15 µm (determined by profilometry of flow wafers, Alpha-Step D-500 from KLA-Tencor with 13.2 mg stylus force) with the area of the enclosed region of the chamber (calculated from AutoCAD design files). *A_spot_* was estimated as 35 µm from calibrated images. Average distance between neighboring ADs was calculated as the square root of the inverse of AD surface density, assuming uniform density on the button surface. Physical distances between TetO sites in the minCMV promoter reporter plasmid (ref: ^84^, Addgene #161928) were converted from base pairs to nanometers using a conversion of 0.3 nm per base pair of DNA.

### Circular dichroism

Circular dichroism spectra were collected using a Jasco 815 spectropolarimeter. Purified proteins were diluted to 0.5 mg/mL in 250 µL of buffer (20 mM KH_2_PO_4_, 200 mM KCl, pH 7). Measurements were taken in a 1.0 mm path length glass cuvette. Wavelength scans were collected and averaged over 3 accumulations.

### Microscale thermophoresis

Microscale thermophoresis equilibrium affinity measurements were collected using a Monolith NanoTemper using Monolith LabelFree Premium capillaries (MO-Z025). ADs were expressed in IVTT as 1-5x 25 µL reactions where parts A and B were incubated together for 1 hour on ice before other components were added. Proteins were expressed for 12 hours at 30°C then incubated at 16°C for 16 hours. Concentrations were estimated from a GFP standard curve that was measured on a DeNovix spectrophotometer. ADs were then diluted to 125 nM in buffer (20 mM KH_2_PO_4_, 200 mM KCl, 0.05% v/v Tween-20) and checked to ensure no AD formed aggregates. Affinity measurements were collected from a capillary scan across a concentration series of purified, unlabeled TAZ2-SNAP proteins that were mixed with 125 nM of AD. Affinities were estimated using the Monolith Affinity Analysis software.

### AlphaFold-Multimer plDDT and PAE for KIX and TAZ2 binders

AlphaFold-Multimer^61^ models were generated using the ColabFold v1.1.5 web server using alphafold_multimer_v2 with MSA from mmseqs2_uniref_env, num_recycles=12, pairing_strategy=’greedy’, relax_max_iterations=200. Top 5 ranked models were analyzed for plDDT and PAE, with reported scores representing the mean across top 5 models. plDDT was calculated for a 15-aa sliding window across each AD, with the maximum 15-aa tile plDDT reported for each AD. Mean PAE values per AD residue and per co-A residue corresponding to inter-protein contacts were calculated using the lower of each off-diagonal value within the PAE matrix, as described by Oefnner et al., 2022^121^. Again, mean, minimum, and maximum values across 15-aa sliding window were calculated and averaged across top 5 ranked models. ColabFold accessed at https://colab.research.google.com/github/sokrypton/ColabFold/blob/main/AlphaFold2.ipynb.

### TAZ2-binding SLiM annotation

SLiMs within TAZ2-binding ADs were discovered using the programs FIMO^66^ and XSTREME^65^ using shuffled sequences as negative controls.

### Cell culture

All experiments presented here were carried out in K562 cells (ATCC, CCL-243, female). Cells were cultured in a controlled humidified incubator at 37C and 5% CO_2_ in RPMI 1640 (Gibco, 11-875-119) media supplemented with 10% FBS (Takara, 632180), and 1% Penicillin Streptomycin (Gibco, 15-140-122). HEK293T-LentiX (Takara Bio, 632180, female) cells (used to produce lentivirus, as described below) were grown in DMEM (Gibco, 10569069) media supplemented with 10% FBS (Takara, 632180) and 1% Penicillin Streptomycin Glutamine (Gibco, 10378016). minCMV reporter (pDY32, Addgene: 161928) cell line generation is described in ref^84^. These cell lines were not authenticated. All cell lines routinely tested negative for mycoplasma.

### HT-recruit with dCBP-1 PROTAC

Plugs of cells containing the Activating Hits Validation Library from ref:^20^ were thawed and expanded to go from ∼4,000x to ∼30,000x coverage. Two replicates were treated with 500 nM P300/CBP PROTAC (dCBP-1, MedchemExpress) and two replicates were treated with vehicle (0.1% DMSO). After 6 hours of treatment, 1,000 ng/mL of doxycycline was added to induce reporter localization. Fresh 500 nM of PROTAC was also added at this time. After 18 hours of doxycycline treatment, the cells were spun down at 300 x *g* for 5 minutes and media was aspirated. Note, this is an earlier time point than measured previously for this library (18 hours here, 48 hours before). This earlier time-point was chosen to minimize cell toxicity due to the PROTAC but still allow for activation. Cells were then resuspended in the same volume of PBS (GIBCO), and the spin down and aspiration was repeated to wash the cells and remove any IgG from serum. Dynabeads M-280 Protein G (ThermoFisher, 10003D) were resuspended by vortexing for 30 s. 50 mL of blocking buffer was prepared per 2 x 10^8^ cells by adding 1 g of biotin-free BSA (Sigma Aldrich) and 200 mL of 0.5 M pH 8.0 EDTA into DPBS (GIBCO), vacuum filtering with a 0.22-mm filter (Millipore), and stored on ice until needed. 60 uL of beads/10 million cells were used and magnetic separation was performed as previously described in ref:^20,84^. Library preparation, sequencing, and enrichment computations were performed as previously described in ref:^20^.

### Individual recruitment assays

Protein fragments were cloned as a fusion with rTetR upstream of a T2A-mCherry-BSD marker (**Extended Data Fig. 5g**) using GoldenGate cloning in the backbone pJT126 (Addgene #161926). K562 citrine reporter cells (pDY32 Addgene: 161928, stably integrated at the AAVS1 locus) were then transduced with each lentiviral vector and, 3 days later, selected with blasticidin (10 mg/mL) until > 80% of the cells were mCherry positive (6-9 days). Cells were split into separate wells of a 24-well plate and either treated with doxycycline (Fisher Scientific) or left untreated. Time points were measured by flow cytometry analysis of >10,000 cells (Biorad ZE5). Doxycycline was assumed to be degraded each day and fresh doxycycline media was added each day of the timecourse.

In order to estimate the fraction of cells ON from HT-recruit enrichment scores, we used a calibration curve between screen enrichment ratios and the fraction of cells ON from individual recruitment assays as measured by flow cytometry, fit to a logistic function (see Extended Data Fig. 3e in^20^).

### Analytical solutions to multivalent binding models

Kinetic models of polyvalent and avid binding models were defined based on prior work^122^. These models were defined by systems of ordinary differential equations representing conversion of species between kinetic steps, which were then solved at equilibrium algebraically (setting the rate of change for all species to 0). To simulate effects on varying parameters on *K_d_* and *R_max_*, we solved the systems as a function of co-A concentrations, simulating binding curves as measured in experiments. For details on parameterization, see **Supplementary Note 2**.

### Bound AD/co-A simulations

Initial complex structures of 80 residue ADs bound to folded co-As were generated with AlphaFold-Multimer version 3^61^ using the default ColabFold parameters with AMBER relaxation. For each system, we designated the specific AD “motif” residues in the bound structure using a combination of manual inspection, AF2 PAE scores, and tiling deletion in-cell activity data^20^. The three highest-confidence structures for each system underwent a subsequent 3-step minimization procedure in order to be used as replicate starting structures for the all-atom simulations. First, the output AF2 structures were minimized in explicit solvent using GROMACS. Second, the AD structures were trimmed to the motif, capped with ACE/NME, and minimized using CAMPARI/ABSINTH^68,123^. Lastly, the motif-co-A complexes underwent a 2 nanosecond low temperature MD simulation, also using CAMPARI/ABSINTH. The purpose of this multistep minimization process was to ensure that the structures of the bound motif were energetically compatible with the ABSINTH implicit solvent model. This minimized bound structure was then used for subsequent Monte Carlo (MC) simulation in CAMPARI. In our simulations, we imposed harmonic restraints between AD motif alpha carbons and nearby folded domain alpha carbons to ensure that the IDR did not dissociate over the course of the simulation. During the simulation, we restricted the MC moveset such that the IDR motif and folded domain residues were only allowed to sample sidechain dihedral rotations, while the flanking IDR residues were allowed to sample from sidechain and backbone dihedral moves. Additional information on minimization and simulation parameters and keyfiles can be found in the supporting information GitHub repository: https://github.com/holehouse-lab/supportingdata/tree/master/2024/delrosso_suzuki_et_al_2024.

We performed three replicate simulations from each of the starting structures (9 replicates total per system). In addition to simulations using the full energy Hamiltonian, we also ran excluded volume (EV) simulations in parallel in which the only energies in the system were steric repulsions. These EV simulations enabled us to normalize across the geometric constraints in our different simulation starting structures. For each simulation, both full and EV, we computed an inter-residue contact frequency map, by calculating the fraction of simulation frames where a pair of IDR and folded domain alpha carbons were within 10 angstroms. We subtracted the EV contact frequency maps from the full simulation maps, then summed this map over all flanking IDR residues to compute the flanking contacts score.

### Coarse-grained STAMMPPING simulations

Simulations for modeling multivalent binding AD/co-A binding were carried out using PIMMS, a coarse-grained, lattice-based, Monte Carlo simulation engine^124^ (https://github.com/holehouse-lab/pimms). We manually defined bead types and chain architectures *in silico* in order to approximate our experimental STAMMPPING system. Specifically, we ran simulations using a 300x300x300 lattice. On one face of this lattice, we initialized a 10x10 evenly-spaced grid of inert “nanobody” chains, which were kept fixed in place during the simulation. The terminal bead of the nanobody chain was bound (using an arbitrarily strong interaction) to one end of the “AD” chain, which consisted of two “binding motif” beads within a chain of inert “IDR” beads. In addition to tethered ADs, these simulations contained freely diffusing “co-A” chains, which were composed of two terminal “co-A domain” beads connected by an inert linker. We titrated the total number of co-A chains within our simulations and varied the pairwise interaction energies between the two co-A domain beads and AD binding motif beads. We defined binding events in our simulations as occurrences where co-A domains were adjacent to AD binding motifs. Using this definition, we computed binding affinities by quantifying the ratio of bound co-A to total ADs, and extracting *K*_d_ and R_max_ by fitting to a Langmuir model. Example PIMMS keyfiles and parameter files for running these simulations, as well as a more detailed description of these files, can be found at https://github.com/holehouse-lab/supportingdata/tree/master/2024/delrosso_suzuki_et_al_2024.

**Supplementary Note 1.**
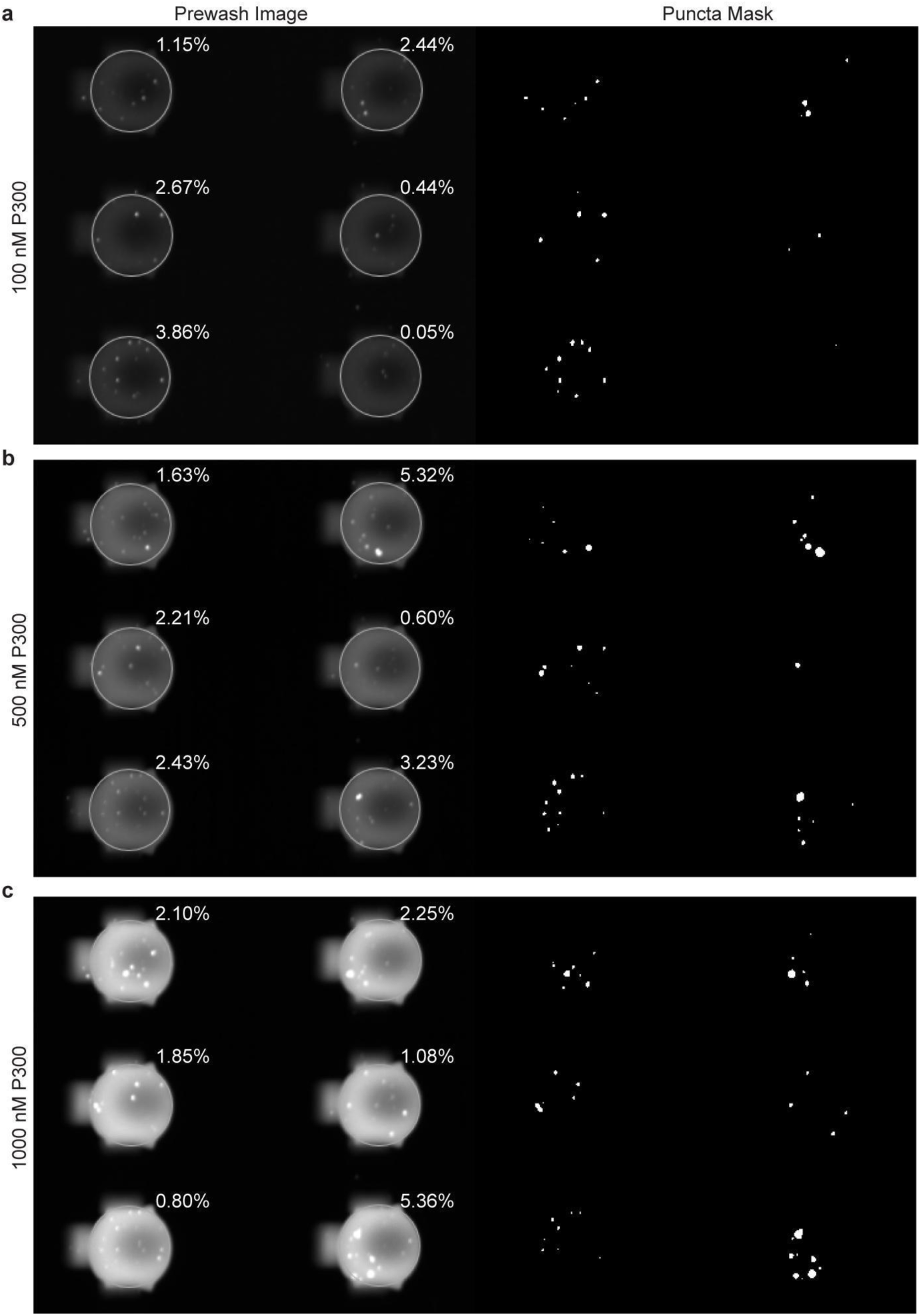
In order to estimate the fraction of P300 that is non-soluble and trapped in punctae, we used PunctaTools^125^ to enumerate and quantify the cumulative intensities within fluorescent punctae visible in the images taken before washing away any unbound fraction (Prewash Image). First, we generated binarized “Puncta Masks” denoting contiguous clusters of pixels associated with visible punctae. Next, we calculated the percentage of protein sequestered within punctae in these Prewash Images by: (1) multiplying (using ImageJ’s Calculator Plus plugin) each Puncta Mask by each Prewash Image to extract the pixel intensities of only the non-soluble protein, (2) drawing a region of interest (ROI) subtending each chamber (white overlay), and then (3) dividing the puncta fraction summed pixel intensities (ImageJ’s Integrated Density’s “RawIntDen”) by the total summed pixel intensities per chamber to generate a normalized estimate of the fraction of the total protein intensities contained in punctae. If we assume P300 can only bind ADs in its soluble form and that any P300 within non-soluble fluorescent punctae cannot bind ADs, partitioning of P300 into punctae at a given concentration should lower the effective concentration of available binding-competent protein such that fitted macroscopic affinities (*K*_d_s) represent an upper limit.

### Supplementary Note 2 | Estimating relative concentrations for use in avid binding models

Polyvalent models assume independent binding by each individual site and therefore predict multivalent affinities based only on the intrinsic affinities of individual potential interactions:

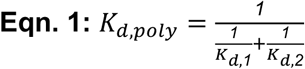

Avid models, by contrast, require estimating the effective concentration (*C_eff_*) of ADs accessible at any given time by a single freely diffusing co-A molecule:

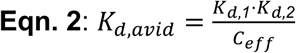

To estimate these effective concentrations, we: (1) calculate the volume subtended by a single freely-diffusing co-A (*R_ee_*, computed using linker lengths predicted using ALBATROSS^75^), (2) determine the number of surface-immobilized ADs within this volume (*n_ADlocal_*, based on the calculated eGFP surface density), and then (3) insert these values into **Eqn. 3**:

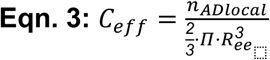

#### Case 1: P300 KIX/synthetic AD library measurements

For P300 KIX binding to synthetic AD combinations (**Fig. 3e**), the polyvalent model predicts *K_d,poly_* = 1/(1/*K_d,MYB_* + 1/*K_d,DDIT3_*) = 1.05 µM. For the avid model, we estimate that only 1 immobilized AD appears within the spherical volume subtended by the disordered linker between MYB and DDIT3 in the MYB-GS-DDIT3 construct, yielding a *C_eff_* of 30 µM and a predicted affinity of *K_d,avid_* = *K_d,MYB_*K_d,DDIT3_* / *C_eff_* = 0.29 µM. The STAMMPPING measurements here yield an observed affinity of *K_d,MYB-GS-DDIT3_* = 0.11 µM, slightly stronger than the avid prediction. This ∼3-fold discrepancy could either reflect experimental noise or the fact that the avid case pre-pays the entropic penalty of localizing the second binding site, as proposed by Jencks: ΔG_interaction_ term^126^.

#### Case 2: Full-length P300 measurements

Estimates of the total distance subtended by full-length P300 (with disordered linker lengths calculated using ALBATROSS^75^) suggest that a single P300 molecule can simultaneously “bridge” multiple surface-immobilized ADs (**Extended Data Fig. 8**). Most P300-binding ADs (87/115) bound at least one of P300’s annotated binding domains. 34/115 P300-binding ADs were capable of binding multiple domains, as pictured below (**Supp. Note Fig. 1**).

**Supplementary Note Figure 1.**
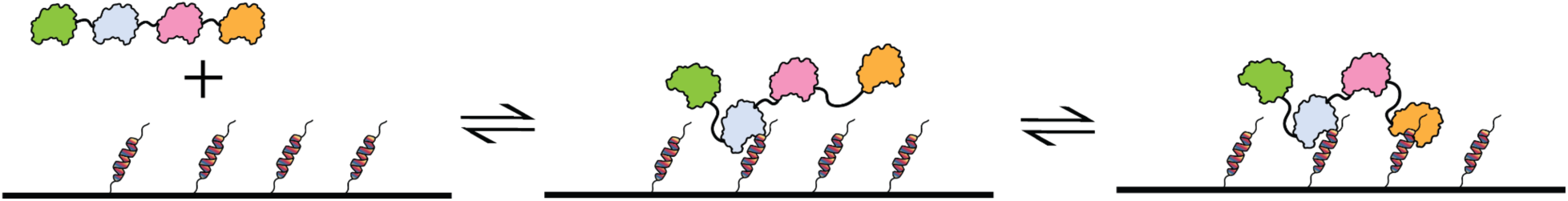
Avid binding between a multi-domain co-A and multi-site AD.

For simplicity, here we estimate a two-site avid binding model in which at most two P300 domains are bound to the AD surface simultaneously. For each AD, we calculated the expected end-to-end distance *R_ee_* between the two P300 domains bound (*e.g.* using an *R_effective_* of 23.3 nm for an AD observed to bind KIX and TAZ2 and an *R*_effective_ of 26.41 nm for an AD observed to bind TAZ1 and TAZ2). Next, we estimated *n_ADlocal_* as the maximum number of AD molecules expected within a sphere of radius *R_ee_* when drawn on a lattice as below (**Supp. Note Fig. 2**, *e.g.* using *n_ADlocal_* = 19 for an AD observed to bind TAZ1 and NCBD, and *n_ADlocal_* = 7 for an AD observed to bind KIX and TAZ2). Two-site avid predicted affinities for each P300-bound AD were calculated using **Eqn. 2** plugging in the two tightest-binding P300 subunit affinities and *C_eff_* from **Eqn. 3** (ranging from 540-880 µM depending on identity of the two P300 domains bound) (**Fig. 3g**).

**Supplementary Note Figure 2.**
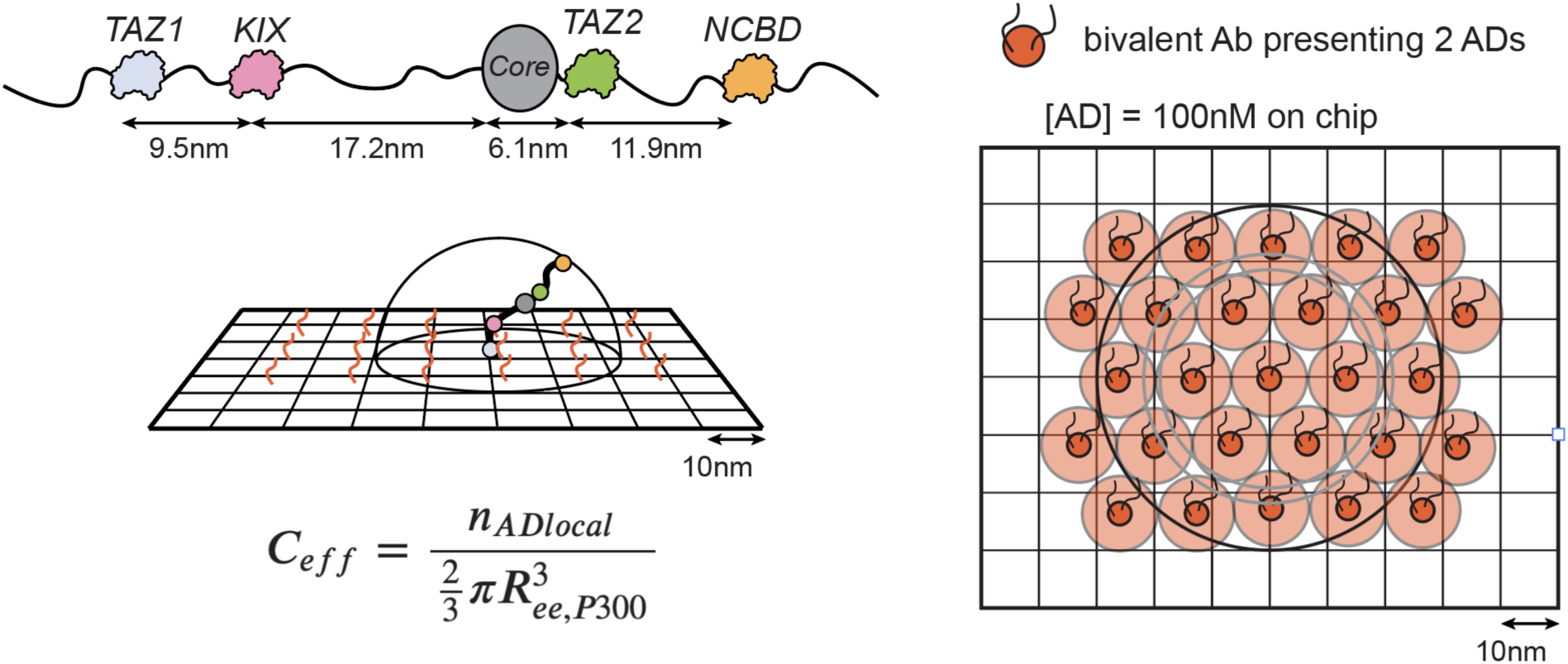
Scale diagram of P300 and estimation of *C_eff_* for prediction of affinity in an avid binding scenario on-chip. P300 IDR lengths estimated by ALBATROSS^75^.

### Supplementary Note 3 | Differentiating between affinity-enhancing models of multivalency

Drawing out toy models can be useful for visualizing expected binding behavior under different assumptions. Here, we consider three families of quantitative models to describe changes in apparent binding: polyvalent binding, avid binding, and allovalent binding. Polyvalent and avid binding are both described in the main text (see **Figure 3**). Allovalent binding is described by a three state model in which partner molecules are either free in solution (**Supp. Note Fig. 3**, left), proximal to each other (**Supp. Note Fig. 3**, middle), or bound (**Supp. Note Fig. 3**, right); through this architecture, allovalent models account for kinetic benefits of multi-site binders by allowing for competition between rebinding to nearby binding sites and diffusion away from the proximal state^69,127^. This addition of a third proximal state can co-occur with either avid or polyvalent binding modes.

**Supplementary Note Figure 3.**
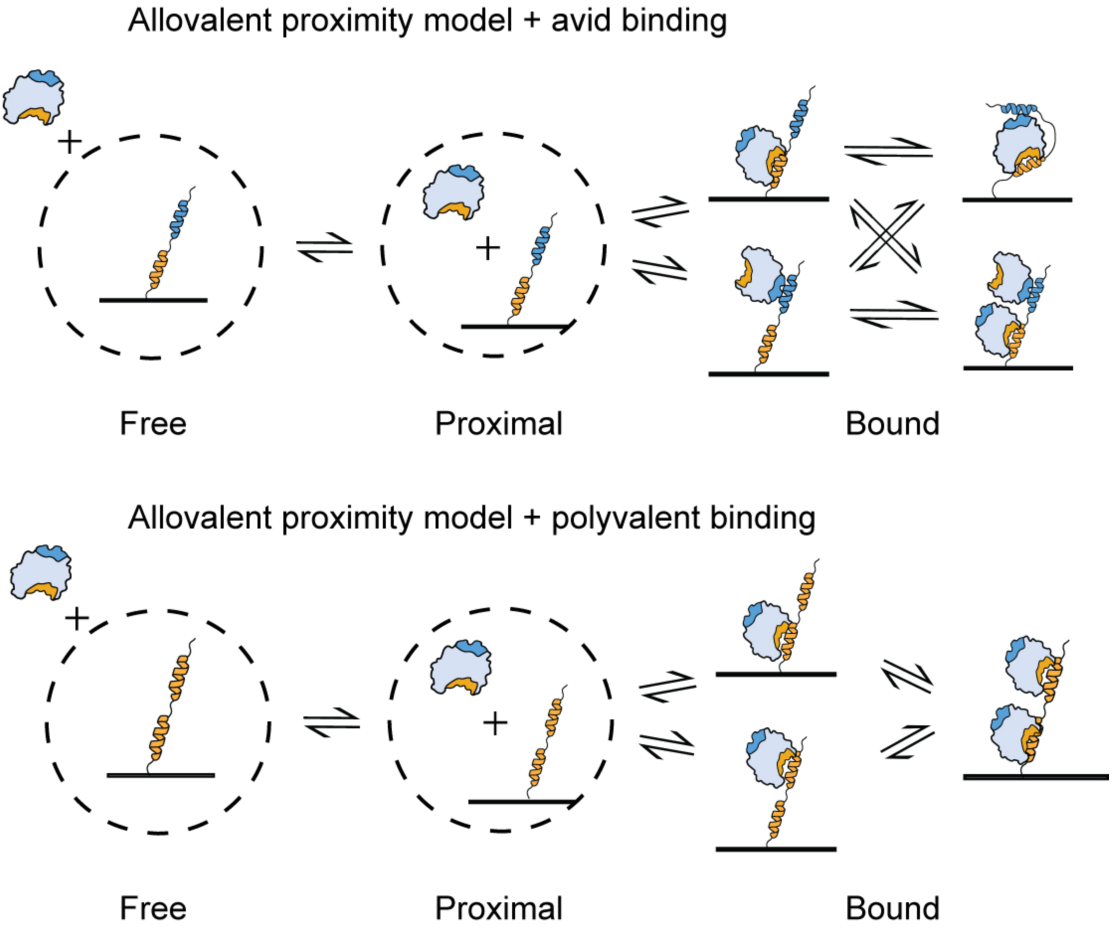
Allovalent binding model representing three states: free, proximal, and bound for both avid (top) and polyvalent binding (bottom).

**Supplementary Note Table 1:**
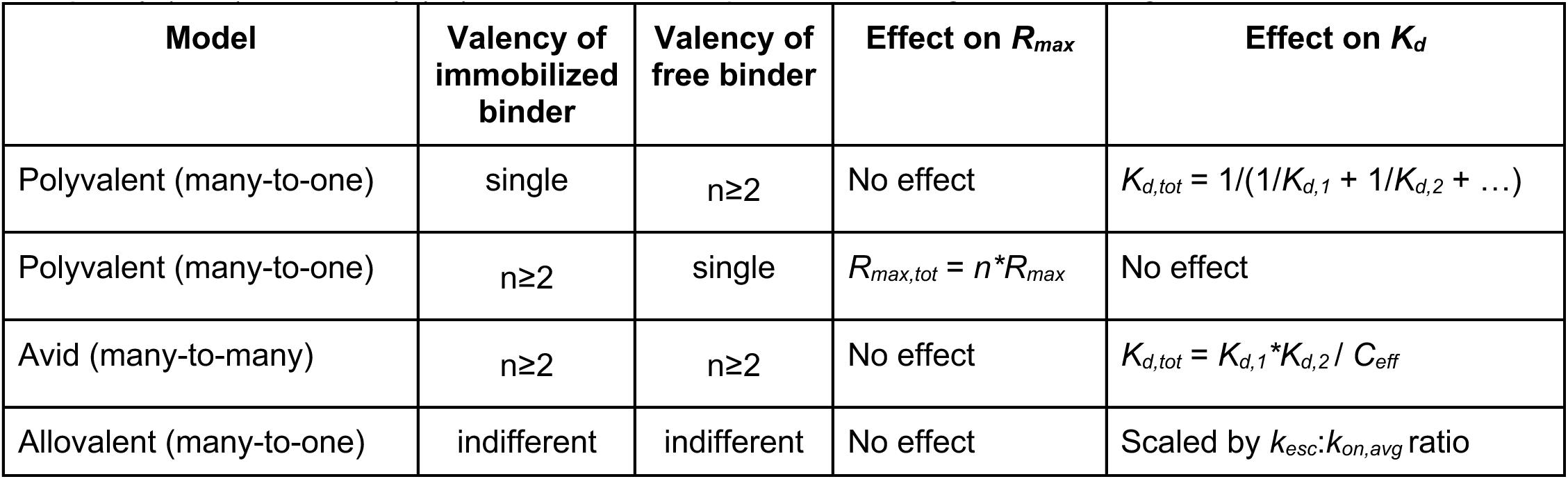
Quantitative models describing multivalent binding and expected impacts on occupancy (*R*_max_) and affinity (*K*_d_) calculated in comparison with single-site binding.

| Model | Valency of immobilized binder | Valency of free binder | Effect on $R_{max}$ | Effect on $K_d$ |
| --- | --- | --- | --- | --- |
| Polyvalent (many-to-one) | single | $n \geq 2$ | No effect | $K_{d,tot} = 1/(1/K_{d,1} + 1/K_{d,2} + \dots)$ |
| Polyvalent (many-to-one) | $n \geq 2$ | single | $R_{max,tot} = n * R_{max}$ | No effect |
| Avid (many-to-many) | $n \geq 2$ | $n \geq 2$ | No effect | $K_{d,tot} = K_{d,1} * K_{d,2} / C_{eff}$ |
| Allovalent (many-to-one) | indifferent | indifferent | No effect | Scaled by $k_{esc} : k_{on,avg}$ ratio |

As shown in **Supplementary Note Table 1** and described in the main text, polyvalent models impact *R_max_* or *K_d_* depending on whether the polyvalent molecule is immobilized (multi-domain AD) or free (multi-domain co-A). Meanwhile, allovalent and avid binding models change affinity (*K_d_*), but not occupancy (*R_max_*), the exception being avid bridging^72^ at co-A concentrations well above both physiological conditions and the concentrations investigated here.

If we observe minimal change in *R_max_*, but an increase in affinity (Fig. 3e), how do we know whether to attribute the increase to avidity effects or allovalency? Here, we have measured only the thermodynamics (and not kinetics) of KIX/synthetic AD binding interactions at equilibrium. Under these conditions, the ‘unbound far’ (Supp. Note Fig. 3, left) and ‘unbound close’ (Supp. Note Fig. 3, middle) states are macroscopically identical such that we cannot distinguish them, and addition of an upstream diffusion step uniformly shifts all affinities for a given co-A (since our synthetic ADs have similar radii of gyration and diffusion constants). Given this expected uniform shift for any double-AD system, the observation of unchanged affinity for polyvalent homotypic ADs compared with their WT counterparts suggests a minimal allovalent effect within our system. Future measurements of association and dissociation for AD constructs with varying diffusion constants (*e.g.* constructs with disordered linkers of different lengths and/or different radii of gyration) could further quantify any contributions of allovalency, but this additional step is not required to explain any increases in KIX/synthetic AD binding affinities observed here.

