## Supplementary figures and images for "High-throughput affinity measurements of direct interactions between activation domains and co-activators"

### Supplementary Figure 1

#### “PS4” 4-block MITOMI device - 2021.06.11

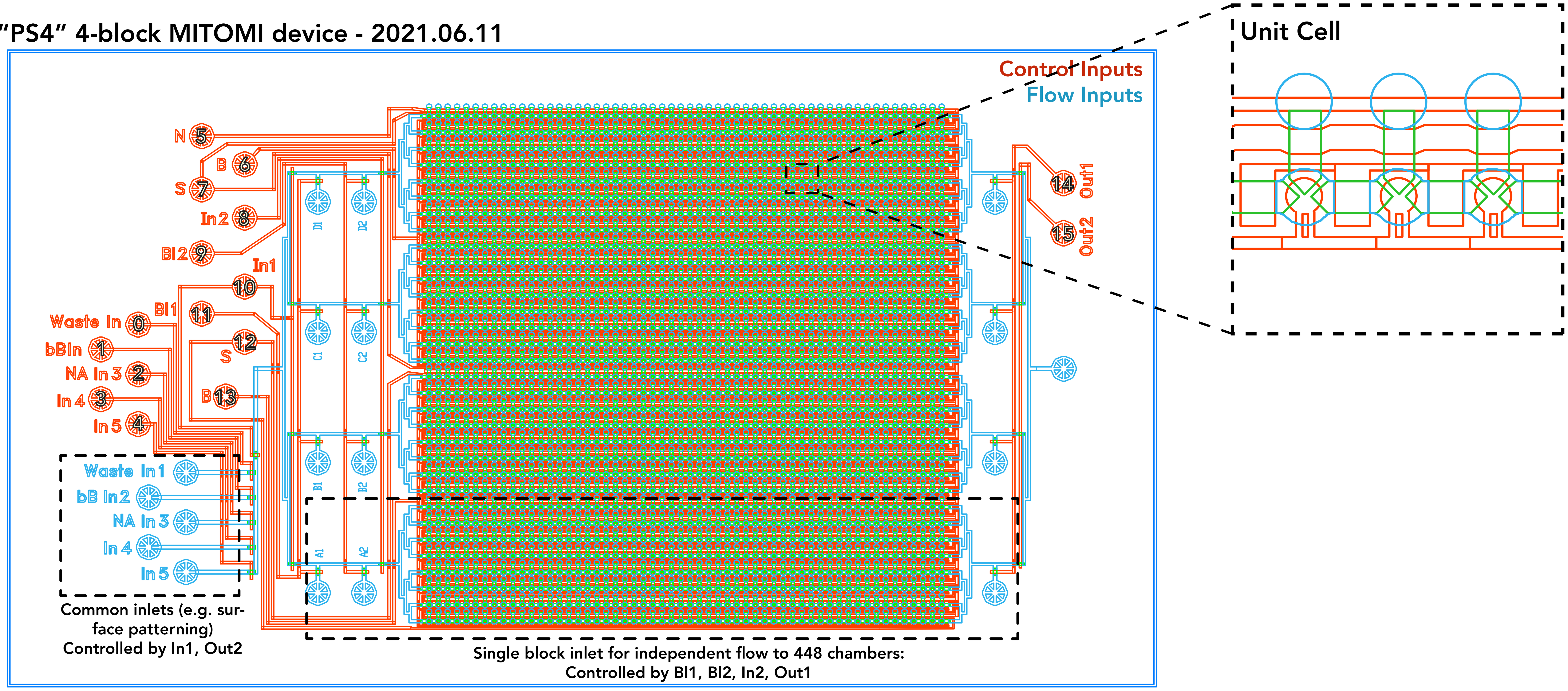


#### “PS8” 8-block MITOMI device - 2021.06.11

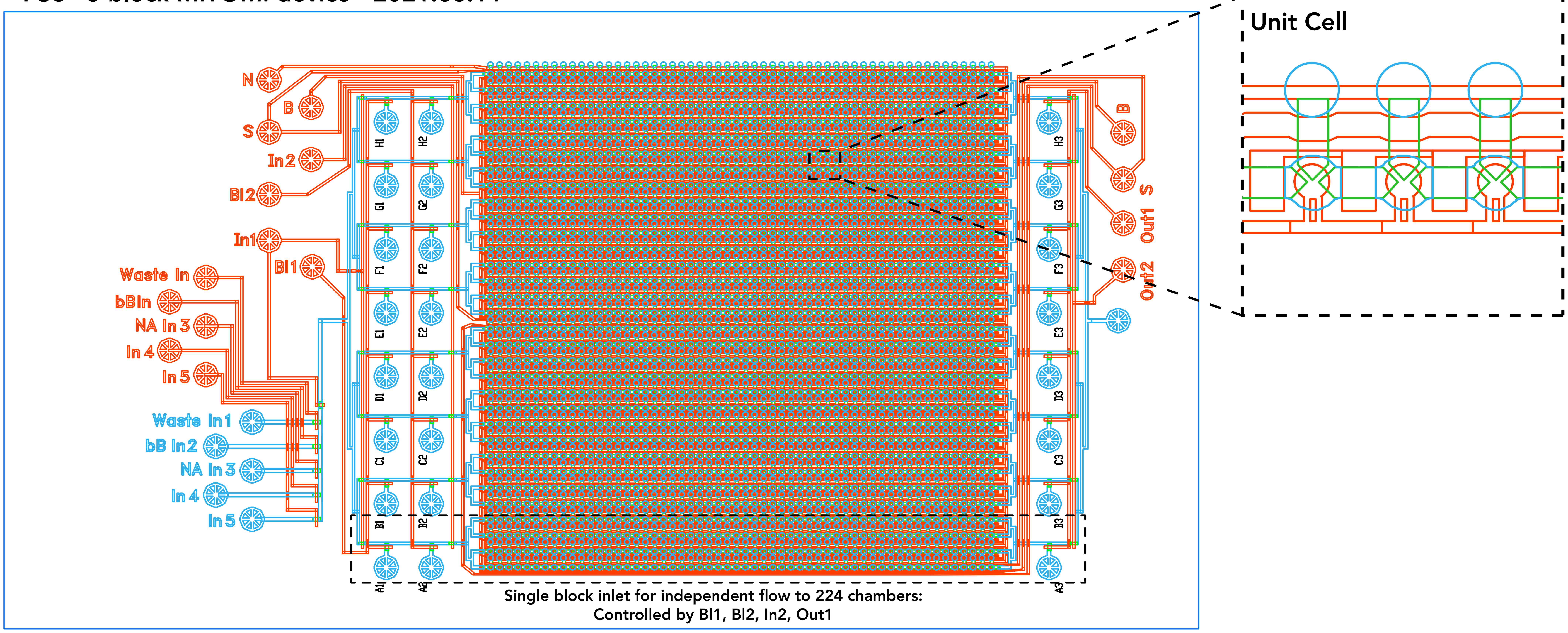

“PS1800” Standard MITOMI device - 2021.06.11

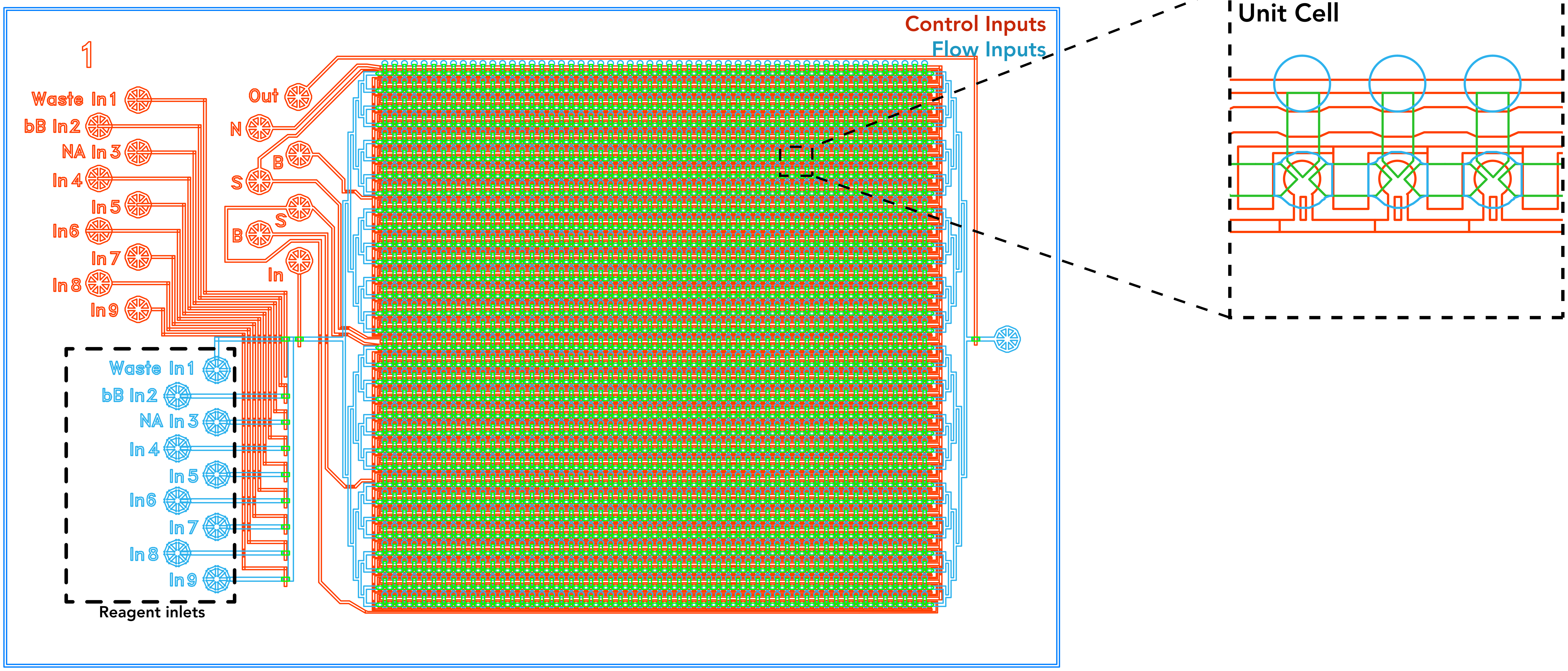

### Supplementary Figure 3

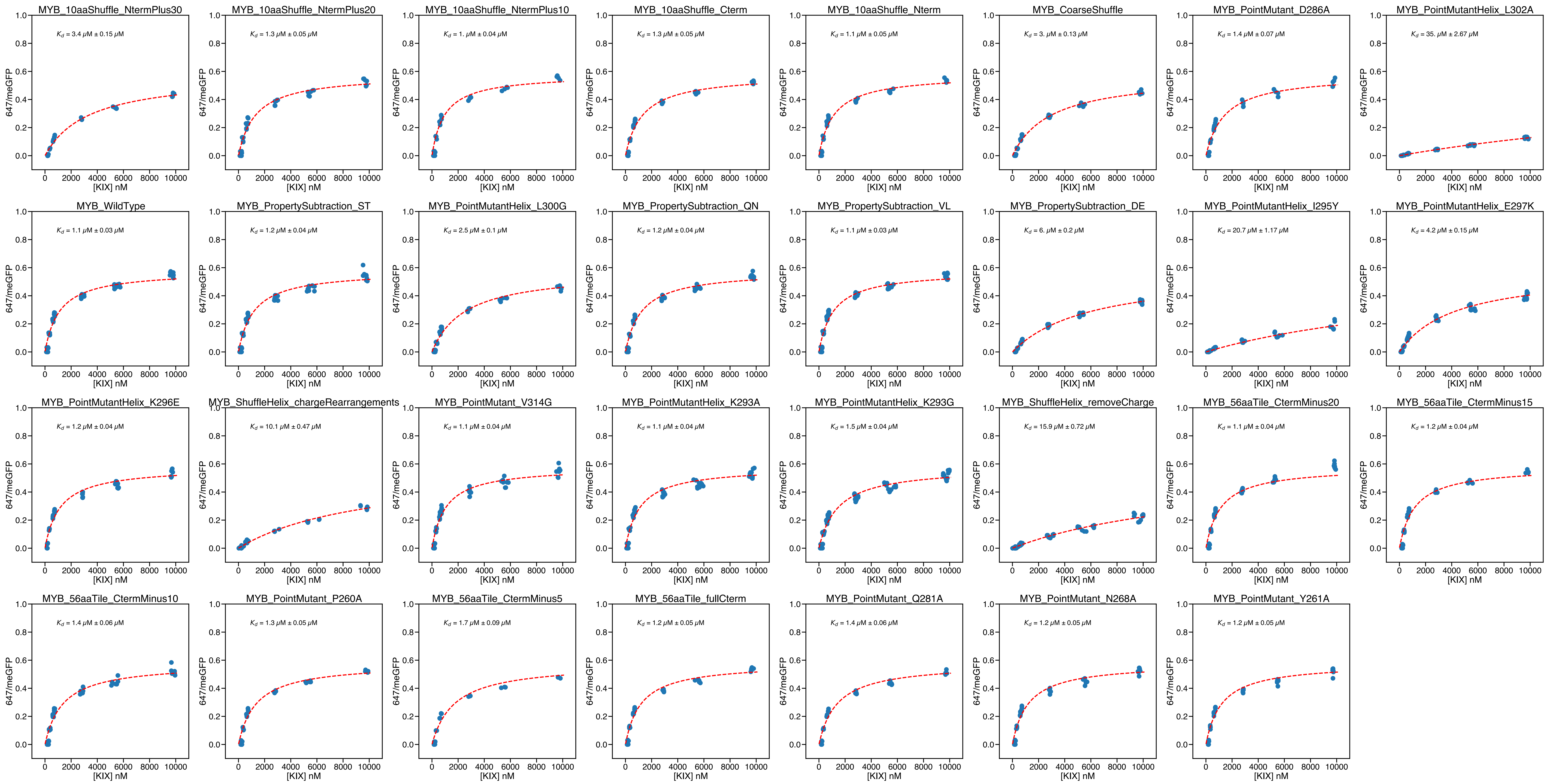

### Supplementary Figure 4

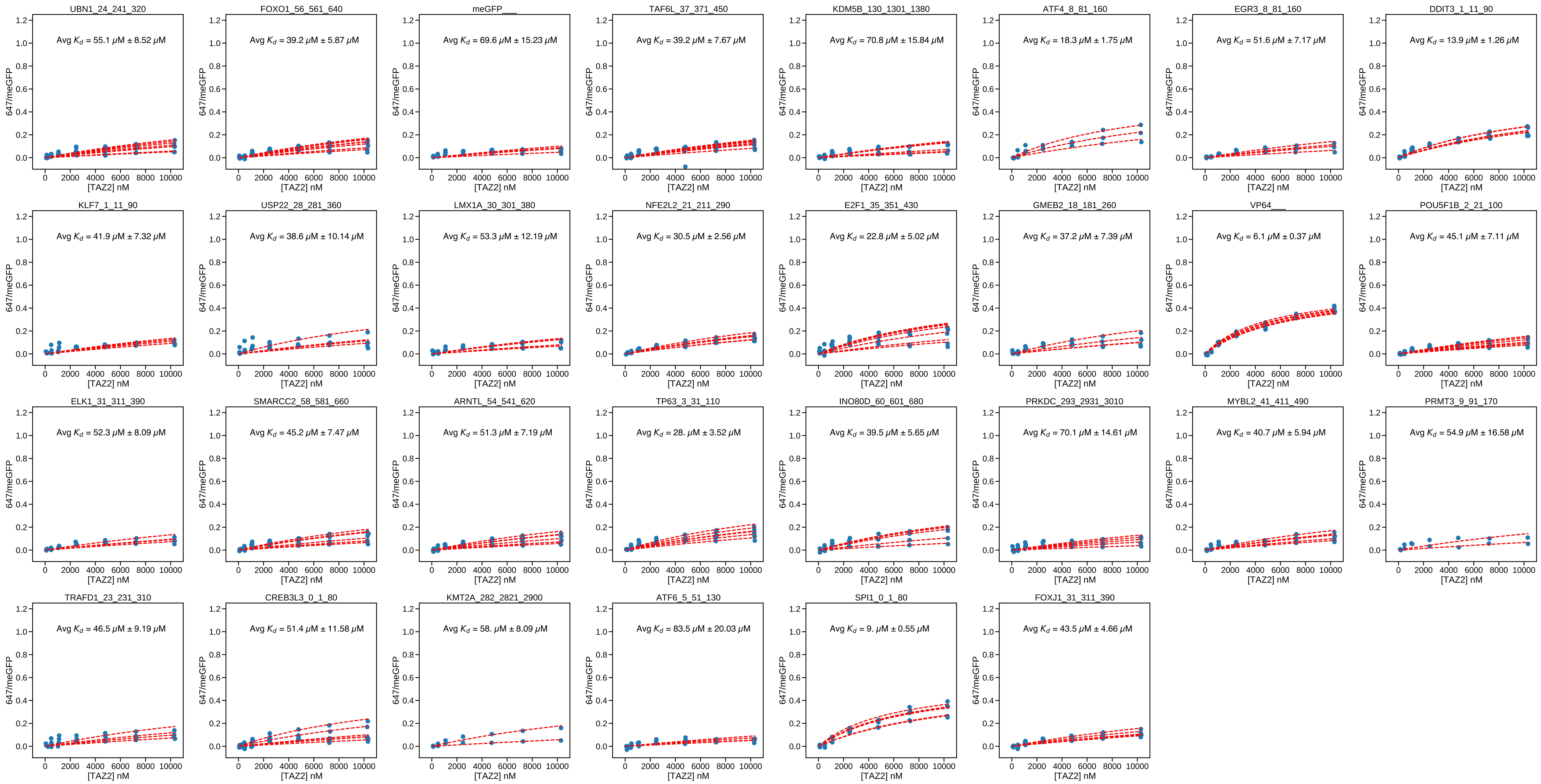

### Supplementary Figure 5

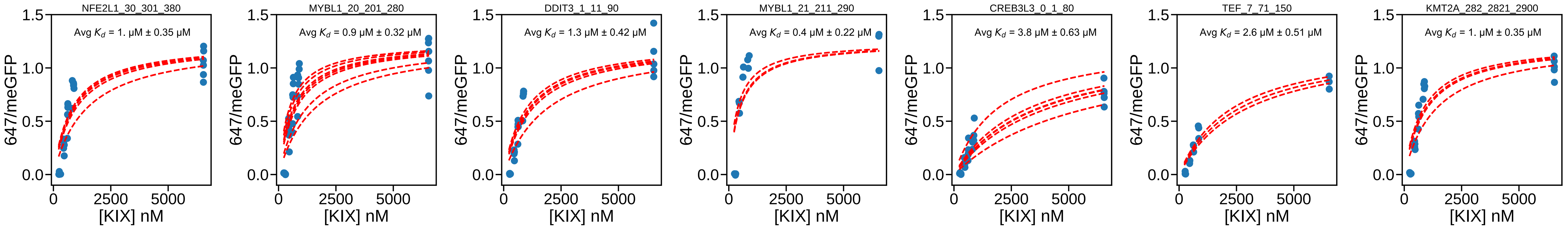


P300 KIX
NaCl Step 1

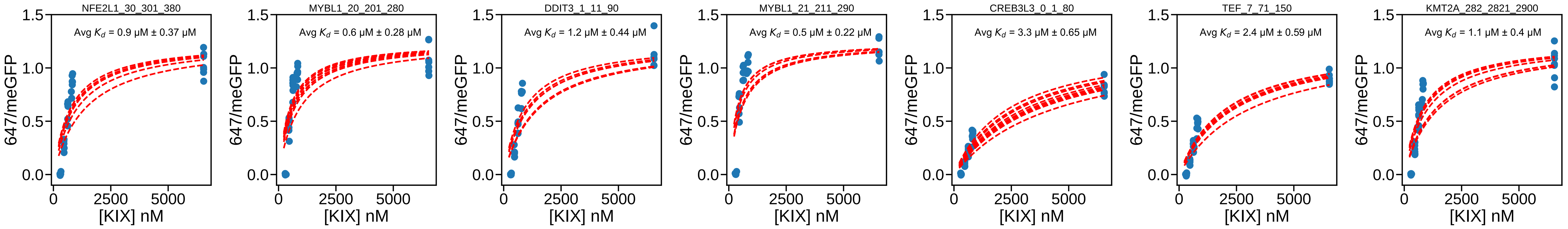


P300 KIX
NaCl Step 2

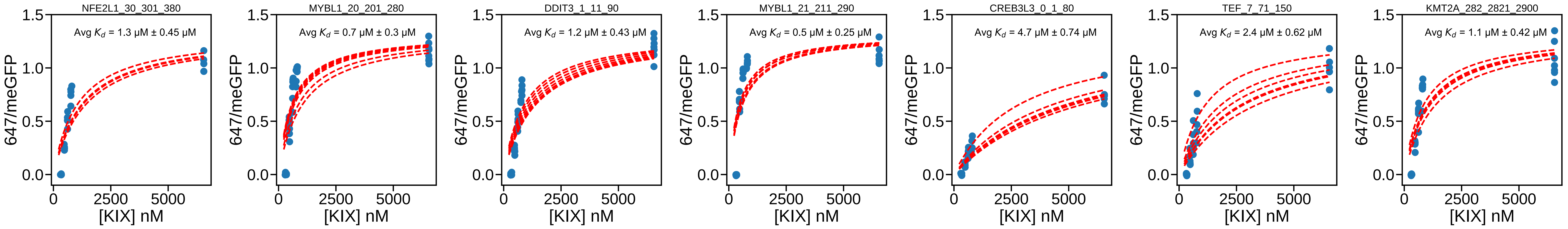


P300 KIX
NaCl Step 3

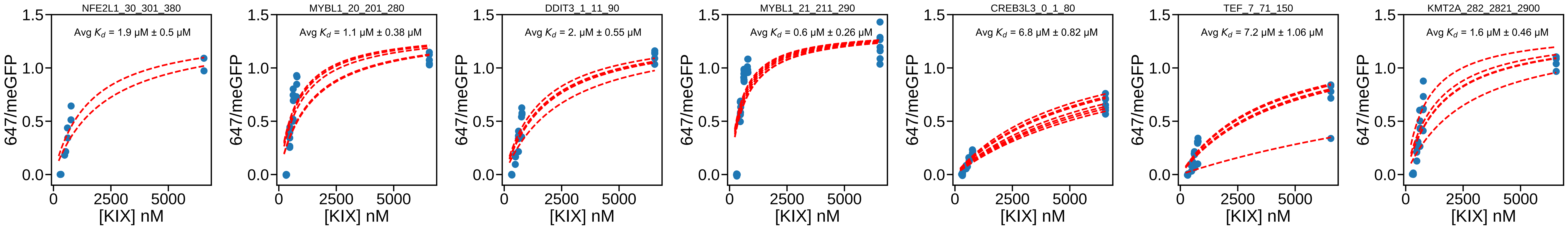


P300 KIX
NaCl Step 4

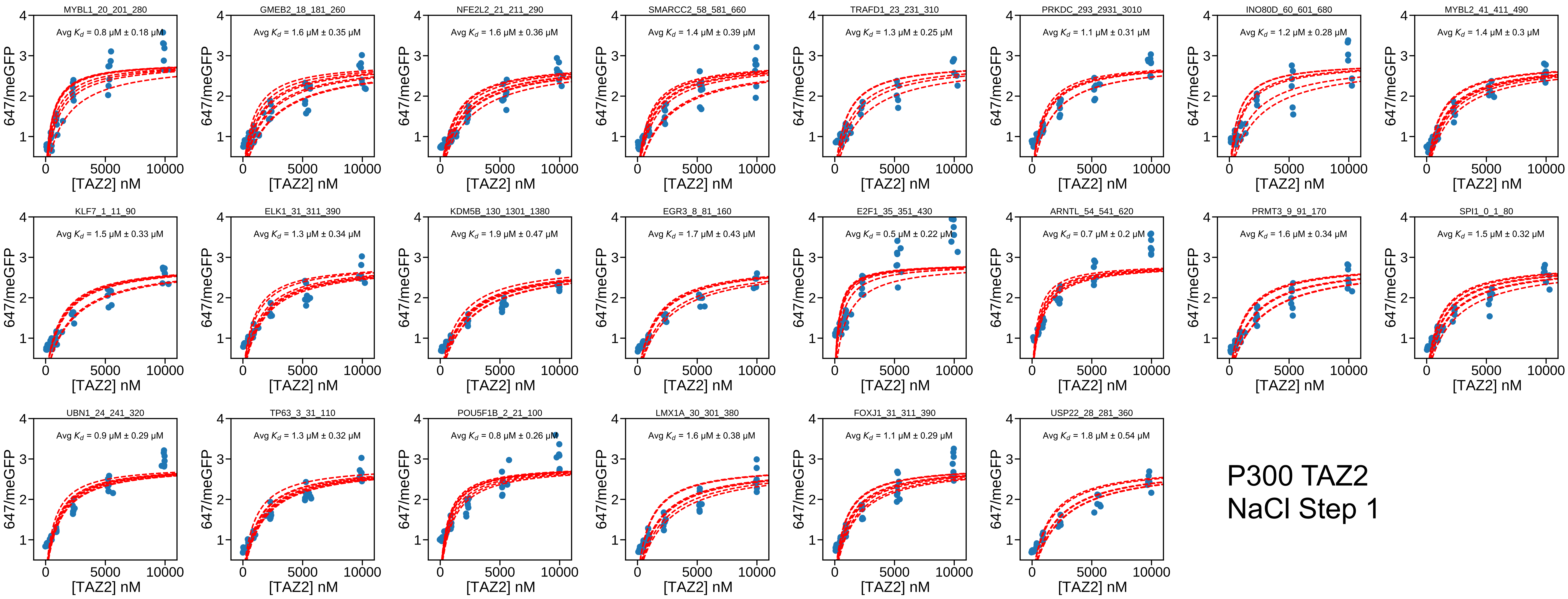

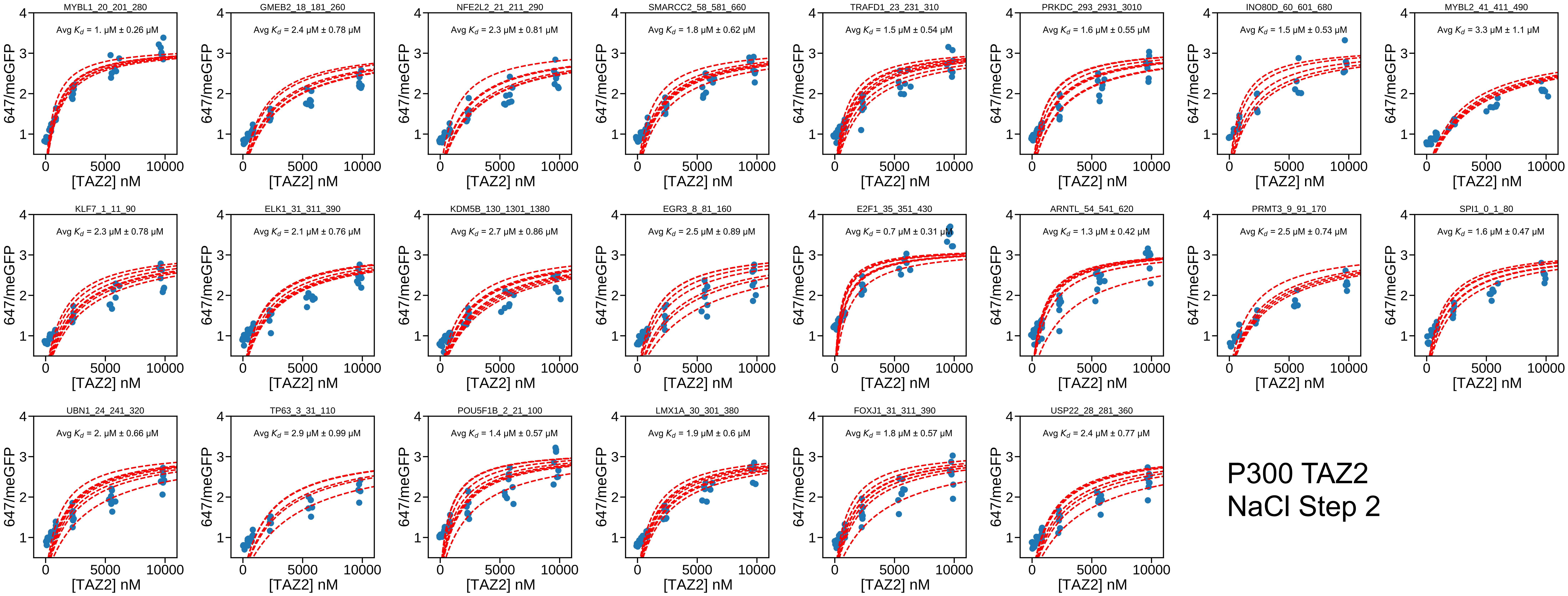

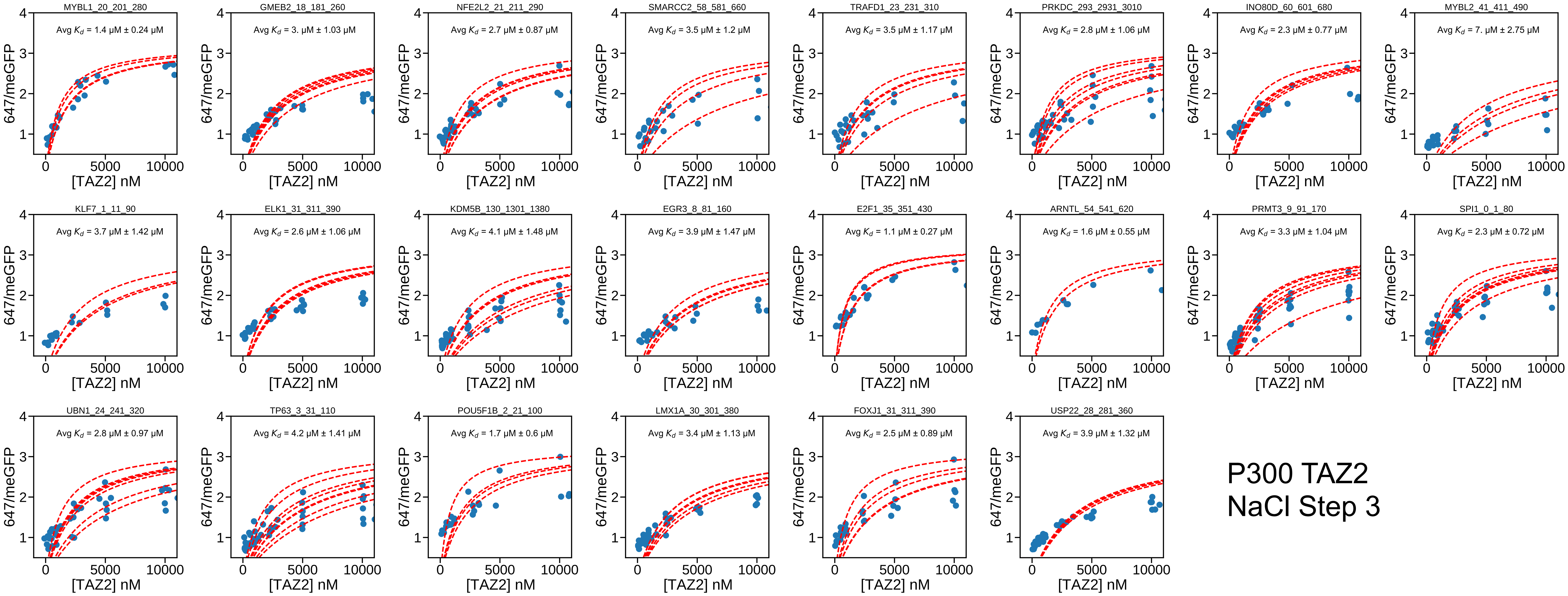

### Supplementary Figure 6

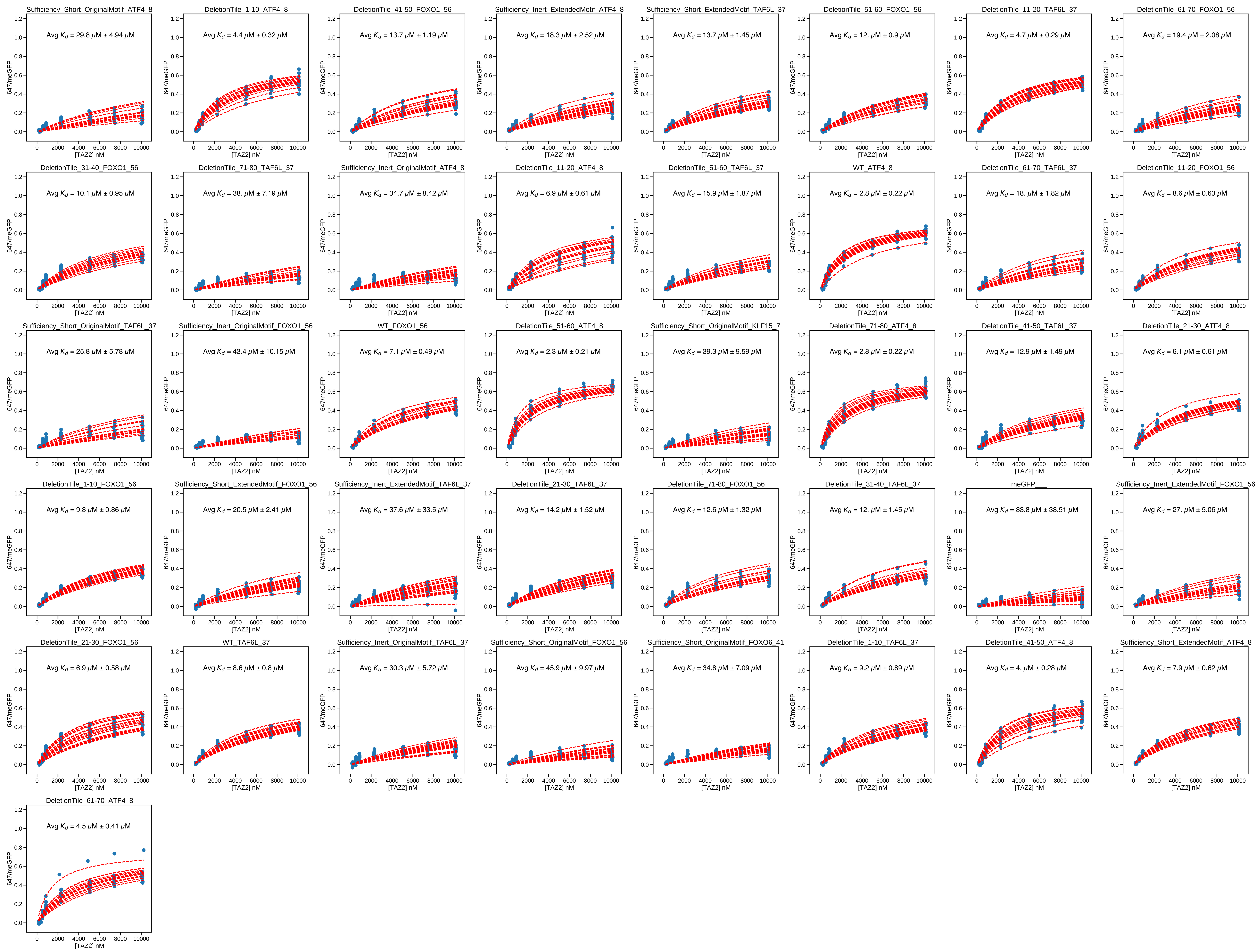

### Supplementary Figure 7

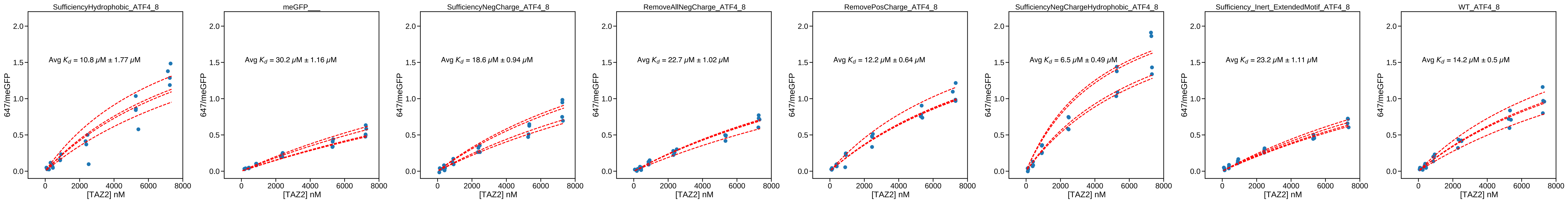

### Supplementary Figure 8

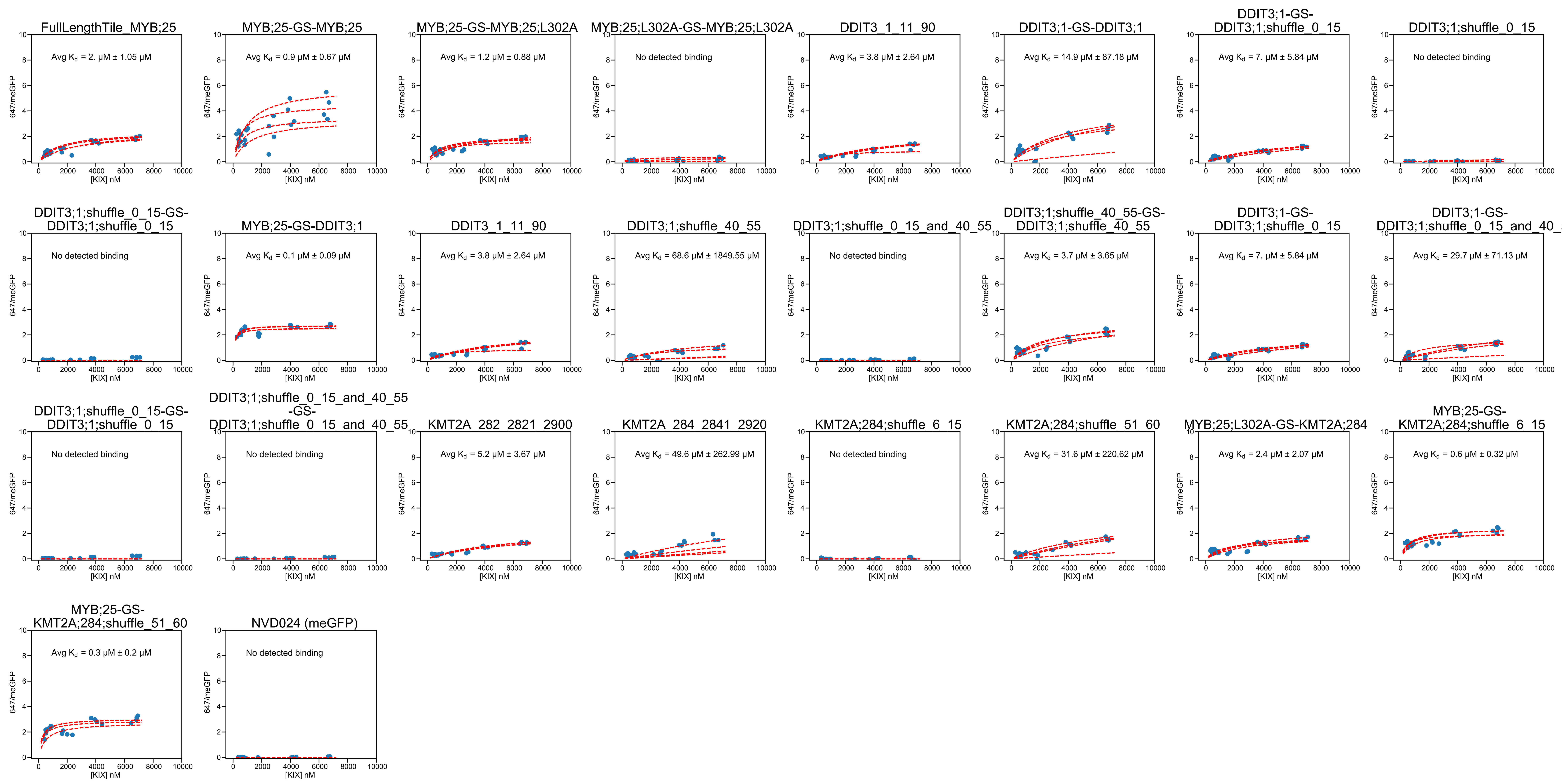

### Supplementary Figure 9

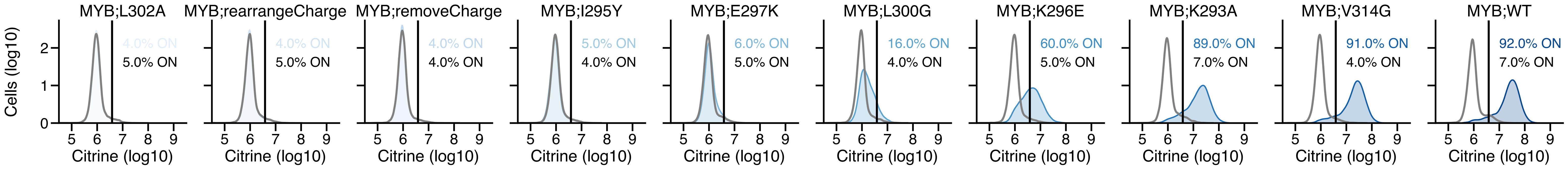

### Supplementary Figure 10

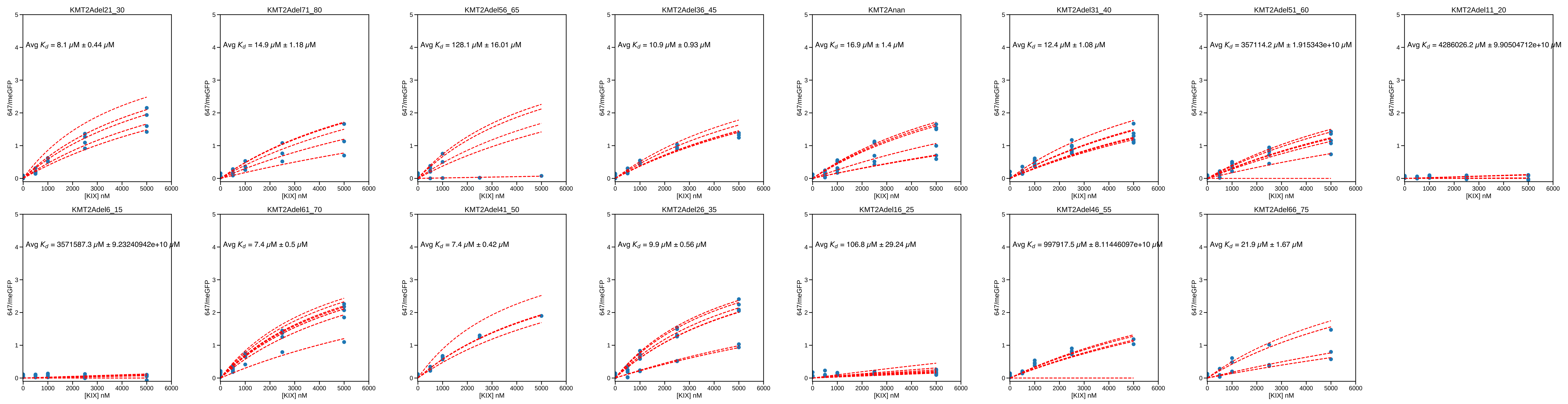

### Supplementary Figure 11

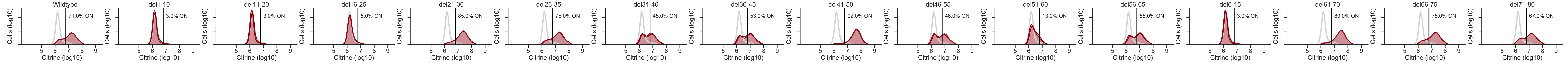
