## Supplementary Figure 2 for "High-throughput affinity measurements of direct interactions between activation domains and co-activators"

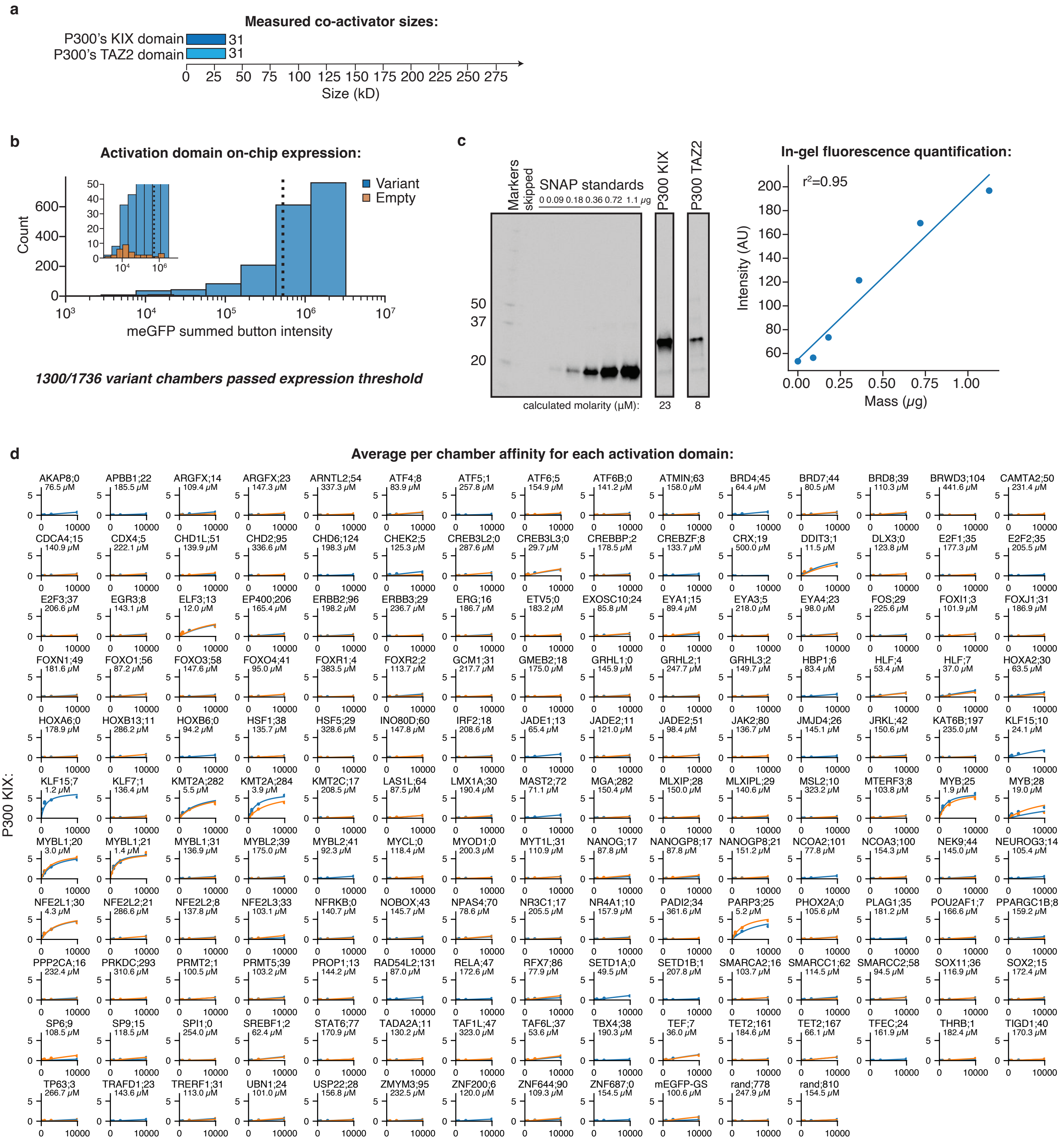
a
Measured co-activator sizes:
P300's KIX domain 31
P300's TAZ2 domain 31
0 25 50 75 100 125 150 175 200 225 250 275
Size (kD)
b
Activation domain on-chip expression:
Variant
Empty
Count
meGFP summed button intensity
1300/1736 variant chambers passed expression threshold
c
Markers
skipped
SNAP standards
0 0.09 0.18 0.36 0.72 1.1 μg
P300 KIX
P300 TAZ2
50
37
20
calculated molarity (μM): 23 8
In-gel fluorescence quantification:
r²=0.95
Intensity (AU)
Mass (μg)
d
Average per chamber affinity for each activation domain:
P300 KIX:
AKAP8;0 76.5 μM
APBB1;22 185.5 μM
ARGFX;14 109.4 μM
ARGFX;23 147.3 μM
ARNTL2;54 337.3 μM
ATF4;8 83.9 μM
ATF5;1 257.8 μM
ATF6;5 154.9 μM
ATF6B;0 141.2 μM
ATMIN;63 158.0 μM
BRD4;45 64.4 μM
BRD7;44 80.5 μM
BRD8;39 110.3 μM
BRWD3;104 441.6 μM
CAMTA2;50 231.4 μM
CDCA4;15 140.9 μM
CDX4;5 222.1 μM
CHD1L;51 139.9 μM
CHD2;95 336.6 μM
CHD6;124 198.3 μM
CHEK2;5 125.3 μM
CREB3L2;0 287.6 μM
CREB3L3;0 29.7 μM
CREBBP;2 178.5 μM
CREBZF;8 133.7 μM
CRX;19 500.0 μM
DDIT3;1 11.5 μM
DLX3;0 123.8 μM
E2F1;35 177.3 μM
E2F2;35 205.5 μM
E2F3;37 206.6 μM
EGR3;8 143.1 μM
ELF3;13 12.0 μM
EP400;206 165.4 μM
ERBB2;96 198.2 μM
ERBB3;29 236.7 μM
ERG;16 186.7 μM
ETV5;0 183.2 μM
EXOSC10;24 85.8 μM
EYA1;15 89.4 μM
EYA3;5 218.0 μM
EYA4;23 98.0 μM
FOS;29 225.6 μM
FOXI1;3 101.9 μM
FOXJ1;31 186.9 μM
FOXN1;49 181.6 μM
FOXO1;56 87.2 μM
FOXO3;58 147.6 μM
FOXO4;41 95.0 μM
FOXR1;4 383.5 μM
FOXR2;2 113.7 μM
GCM1;31 217.7 μM
GMEB2;18 175.0 μM
GRHL1;0 145.9 μM
GRHL2;1 247.7 μM
GRHL3;2 149.7 μM
HBP1;6 83.4 μM
HLF;4 53.4 μM
HLF;7 37.0 μM
HOXA2;30 63.5 μM
HOXA6;0 178.9 μM
HOXB13;11 286.2 μM
HOXB6;0 94.2 μM
HSF1;38 135.7 μM
HSF5;29 328.6 μM
INO80D;60 147.8 μM
IRF2;18 208.6 μM
JADE1;13 65.4 μM
JADE2;11 121.0 μM
JADE2;51 98.4 μM
JAK2;80 136.7 μM
JMJD4;26 145.1 μM
JRKL;42 150.6 μM
KAT6B;197 235.0 μM
KLF15;10 24.1 μM
KLF15;7 1.2 μM
KLF7;1 136.4 μM
KMT2A;282 5.5 μM
KMT2A;284 3.9 μM
KMT2C;17 208.5 μM
LAS1L;64 87.5 μM
LMX1A;30 190.4 μM
MAST2;72 71.1 μM
MGA;282 150.4 μM
MLXIP;28 150.0 μM
MLXIPL;29 140.6 μM
MSL2;10 323.2 μM
MTERF3;8 103.8 μM
MYB;25 1.9 μM
MYB;28 19.0 μM
MYBL1;20 3.0 μM
MYBL1;21 1.4 μM
MYBL1;31 136.9 μM
MYBL2;39 175.0 μM
MYBL2;41 92.3 μM
MYCL;0 118.4 μM
MYOD1;0 200.3 μM
MYT1L;31 110.9 μM
NANOG;17 87.8 μM
NANOGP8;17 87.8 μM
NANOGP8;21 151.2 μM
NCOA2;101 77.8 μM
NCOA3;100 154.3 μM
NEK9;44 145.0 μM
NEUROG3;14 105.4 μM
NFE2L1;30 4.3 μM
NFE2L2;21 286.6 μM
NFE2L2;8 137.8 μM
NFE2L3;33 103.1 μM
NFRKB;0 140.7 μM
NOBOX;43 145.7 μM
NPAS4;70 78.6 μM
NR3C1;17 205.5 μM
NR4A1;10 157.9 μM
PADI2;34 361.6 μM
PARP3;25 5.2 μM
PHOX2A;0 105.6 μM
PLAG1;35 181.2 μM
POU2AF1;7 166.6 μM
PPARGC1B;8 159.2 μM
PPP2CA;16 232.4 μM
PRKDC;293 310.6 μM
PRMT2;1 100.5 μM
PRMT5;39 103.2 μM
PROP1;13 144.2 μM
RAD54L2;131 87.0 μM
RELA;47 172.6 μM
RFX7;86 77.9 μM
SETD1A;0 49.5 μM
SETD1B;1 207.8 μM
SMARCA2;16 103.7 μM
SMARCC1;62 114.5 μM
SMARCC2;58 94.5 μM
SOX11;36 116.9 μM
SOX2;15 172.4 μM
SP6;9 108.5 μM
SP9;15 118.5 μM
SPI1;0 254.0 μM
SREBF1;2 62.4 μM
STAT6;77 170.9 μM
TADA2A;11 130.2 μM
TAF1L;47 323.0 μM
TAF6L;37 53.6 μM
TBX4;38 190.3 μM
TEF;7 36.0 μM
TET2;161 184.6 μM
TET2;167 66.1 μM
TFEC;24 161.9 μM
THRB;1 182.4 μM
TIGD1;40 170.3 μM
TP63;3 266.7 μM
TRAFD1;23 143.6 μM
TRERF1;31 113.0 μM
UBN1;24 101.0 μM
USP22;28 156.8 μM
ZMYM3;95 232.5 μM
ZNF200;6 120.0 μM
ZNF644;90 109.3 μM
ZNF687;0 154.5 μM
mEGFP-GS 100.6 μM
rand;778 247.9 μM
rand;810 154.5 μM

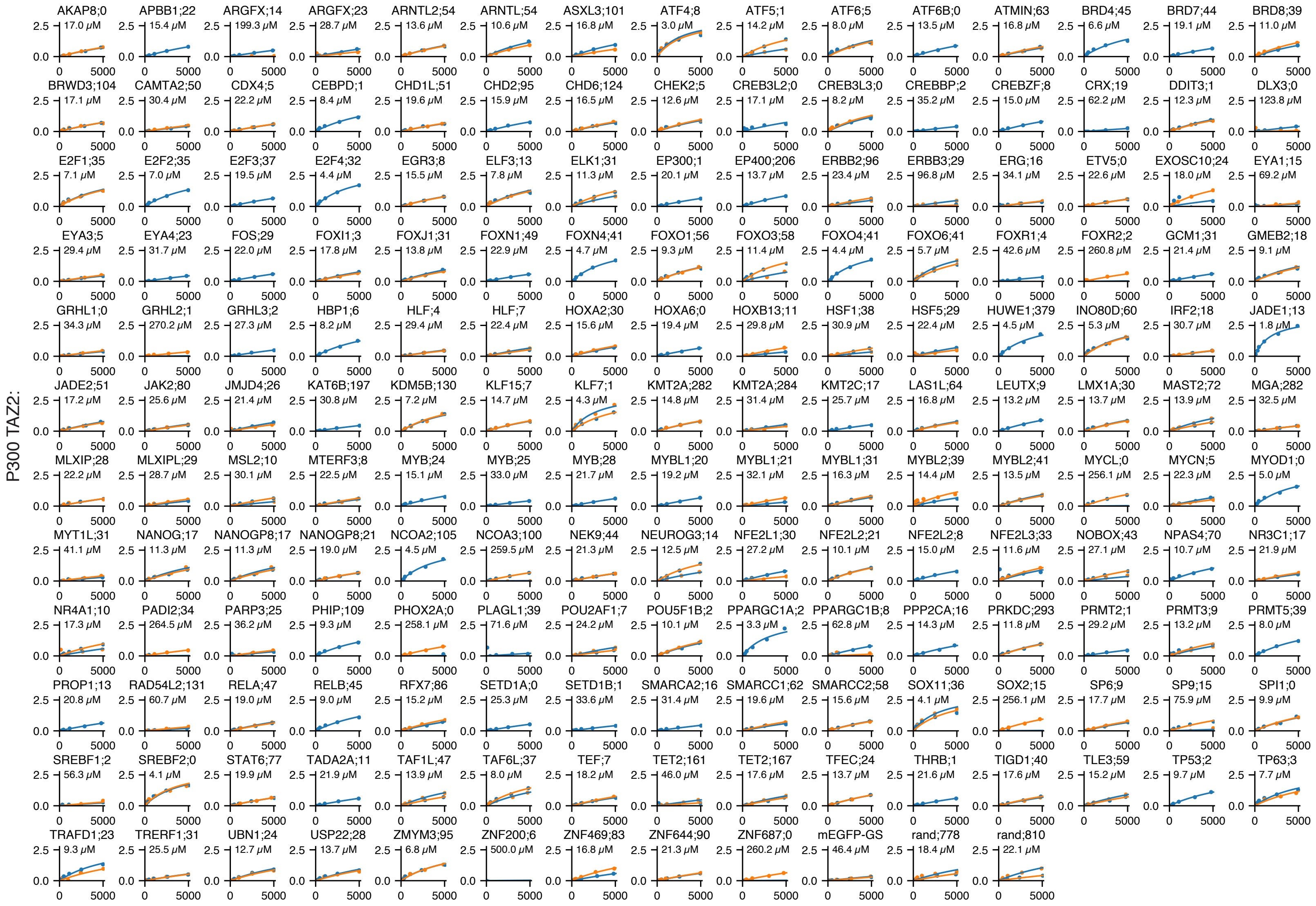


**e**

**Affinities replicability (red=meGFP):**

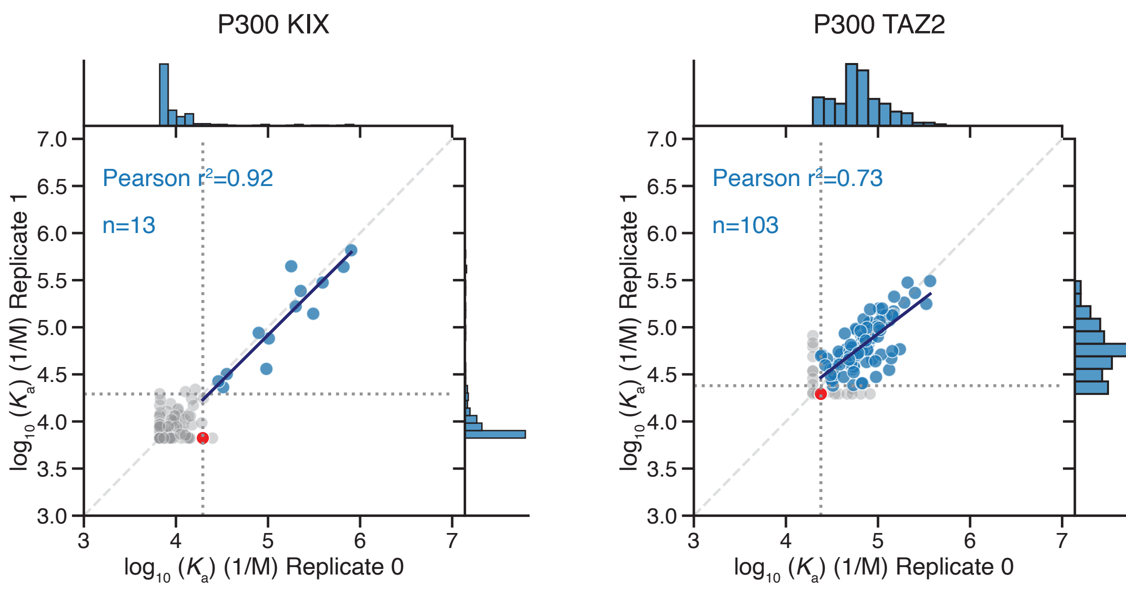


**f**

**Raw images at highest tested concentration co-A (top) AD (bottom)**

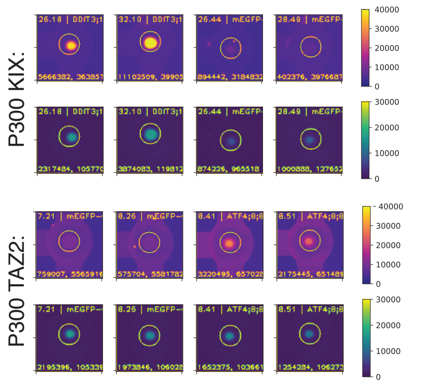

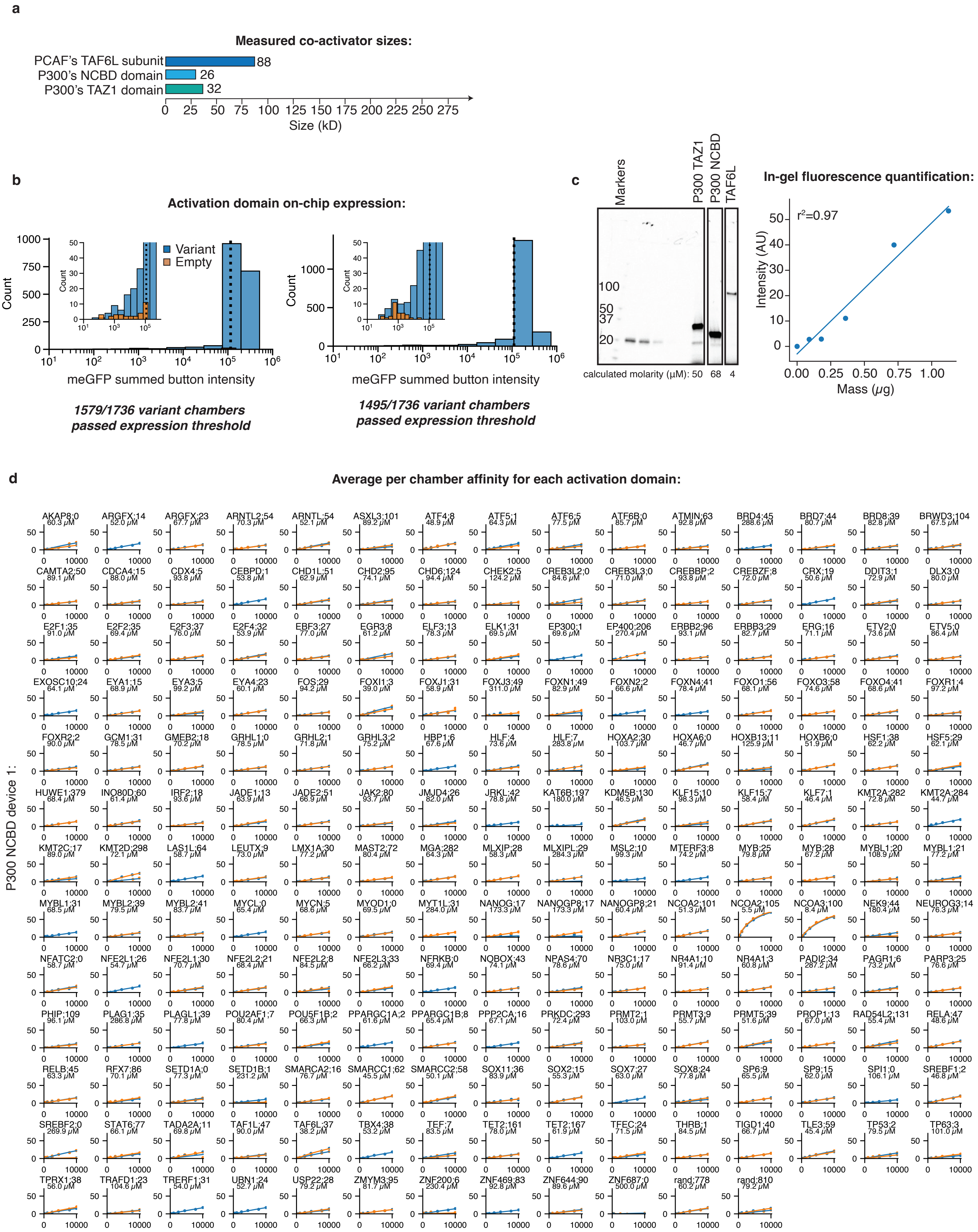
a
Measured co-activator sizes:
PCAF's TAF6L subunit 88
P300's NCBD domain 26
P300's TAZ1 domain 32
0 25 50 75 100 125 150 175 200 225 250 275
Size (kD)
b
Activation domain on-chip expression:
Variant
Empty
Count
meGFP summed button intensity
1579/1736 variant chambers passed expression threshold
1495/1736 variant chambers passed expression threshold
c
Markers
P300 TAZ1
P300 NCBD
TAF6L
100
50
37
20
calculated molarity (μM): 50 68 4
In-gel fluorescence quantification:
r²=0.97
Intensity (AU)
Mass (μg)
d
Average per chamber affinity for each activation domain:
P300 NCBD device 1:
AKAP8:0 60.3 μM
ARGFX:14 52.0 μM
ARGFX:23 67.7 μM
ARNTL2:54 70.3 μM
ARNTL:54 52.1 μM
ASXL3:101 89.2 μM
ATF4:8 48.9 μM
ATF5:1 64.3 μM
ATF6:5 77.5 μM
ATF6B:0 85.7 μM
ATMIN:63 92.8 μM
BRD4:45 288.6 μM
BRD7:44 80.7 μM
BRD8:39 82.8 μM
BRWD3:104 67.5 μM
CAMTA2:50 89.1 μM
CDCA4:15 88.0 μM
CDX4:5 93.8 μM
CEBPD:1 53.8 μM
CHD1L:51 62.9 μM
CHD2:95 74.1 μM
CHD6:124 94.4 μM
CHEK2:5 124.2 μM
CREB3L2:0 84.6 μM
CREB3L3:0 71.0 μM
CREBBP:2 93.8 μM
CREBZF:8 72.0 μM
CRX:19 50.6 μM
DDIT3:1 72.9 μM
DLX3:0 80.0 μM
E2F1:35 91.0 μM
E2F2:35 69.4 μM
E2F3:37 76.0 μM
E2F4:32 53.9 μM
EBF3:27 77.0 μM
EGR3:8 61.2 μM
ELF3:13 78.3 μM
ELK1:31 69.5 μM
EP300:1 69.6 μM
EP400:206 270.4 μM
ERBB2:96 93.1 μM
ERBB3:29 82.7 μM
ERG:16 71.1 μM
ETV2:0 73.6 μM
ETV5:0 86.4 μM
EXOSC10:24 64.1 μM
EYA1:15 68.9 μM
EYA3:5 99.2 μM
EYA4:23 60.1 μM
FOS:29 94.2 μM
FOXI1:3 39.0 μM
FOXJ1:31 58.9 μM
FOXJ3:49 311.0 μM
FOXN1:49 82.9 μM
FOXN2:2 66.6 μM
FOXN4:41 78.4 μM
FOXO1:56 68.1 μM
FOXO3:58 74.6 μM
FOXO4:41 68.6 μM
FOXR1:4 97.2 μM
FOXR2:2 90.0 μM
GCM1:31 78.5 μM
GMEB2:18 70.2 μM
GRHL1:0 78.5 μM
GRHL2:1 71.8 μM
GRHL3:2 75.2 μM
HBP1:6 67.6 μM
HLF:4 73.6 μM
HLF:7 283.8 μM
HOXA2:30 103.7 μM
HOXA6:0 46.7 μM
HOXB13:11 125.9 μM
HOXB6:0 51.9 μM
HSF1:38 62.2 μM
HSF5:29 62.1 μM
HUWE1:379 68.4 μM
INO80D:60 61.4 μM
IRF2:18 93.6 μM
JADE1:13 63.9 μM
JADE2:51 66.9 μM
JAK2:80 93.7 μM
JMJD4:26 82.0 μM
JRKL:42 78.8 μM
KAT6B:197 180.0 μM
KDM5B:130 46.5 μM
KLF15:10 98.3 μM
KLF15:7 58.4 μM
KLF7:1 46.4 μM
KMT2A:282 72.8 μM
KMT2A:284 44.7 μM
KMT2C:17 89.0 μM
KMT2D:298 72.1 μM
LAS1L:64 58.7 μM
LEUTX:9 73.0 μM
LMX1A:30 77.2 μM
MAST2:72 80.4 μM
MGA:282 64.3 μM
MLXIP:28 58.3 μM
MLXIPL:29 284.3 μM
MSL2:10 99.3 μM
MTERF3:8 74.2 μM
MYB:25 79.8 μM
MYB:28 67.2 μM
MYBL1:20 108.9 μM
MYBL1:21 77.2 μM
MYBL1:31 68.5 μM
MYBL2:39 79.5 μM
MYBL2:41 83.7 μM
MYCL:0 65.4 μM
MYCN:5 68.6 μM
MYOD1:0 69.5 μM
MYT1L:31 284.0 μM
NANOG:17 173.3 μM
NANOGP8:17 173.3 μM
NANOGP8:21 60.4 μM
NCOA2:101 51.3 μM
NCOA2:105 5.5 μM
NCOA3:100 8.4 μM
NEK9:44 180.4 μM
NEUROG3:14 76.3 μM
NFATC2:0 58.7 μM
NFE2L1:26 54.7 μM
NFE2L1:30 70.7 μM
NFE2L2:21 68.4 μM
NFE2L2:8 84.5 μM
NFE2L3:33 66.2 μM
NFRKB:0 69.4 μM
NOBOX:43 74.1 μM
NPAS4:70 78.6 μM
NR3C1:17 75.0 μM
NR4A1:10 91.4 μM
NR4A1:3 60.8 μM
PADI2:34 287.2 μM
PAGR1:6 73.2 μM
PARP3:25 76.6 μM
PHIP:109 96.1 μM
PLAG1:35 286.8 μM
PLAGL1:39 77.8 μM
POU2AF1:7 80.4 μM
POU5F1B:2 66.3 μM
PPARGC1A:2 61.6 μM
PPARGC1B:8 65.4 μM
PPP2CA:16 67.1 μM
PRKDC:293 72.4 μM
PRMT2:1 103.0 μM
PRMT3:9 55.7 μM
PRMT5:39 51.6 μM
PROP1:13 67.0 μM
RAD54L2:131 55.4 μM
RELA:47 48.6 μM
RELB:45 63.3 μM
RFX7:86 70.1 μM
SETD1A:0 77.3 μM
SETD1B:1 231.2 μM
SMARCA2:16 76.7 μM
SMARCC1:62 45.5 μM
SMARCC2:58 50.1 μM
SOX11:36 83.9 μM
SOX2:15 55.3 μM
SOX7:27 63.0 μM
SOX8:24 77.8 μM
SP6:9 65.5 μM
SP9:15 62.0 μM
SPI1:0 106.1 μM
SREBF1:2 46.8 μM
SREBF2:0 269.9 μM
STAT6:77 66.1 μM
TADA2A:11 69.8 μM
TAF1L:47 90.0 μM
TAF6L:37 38.2 μM
TBX4:38 53.2 μM
TEF:7 83.5 μM
TET2:161 78.0 μM
TET2:167 61.9 μM
TFEC:24 71.5 μM
THRB:1 84.5 μM
TIGD1:40 66.7 μM
TLE3:59 45.4 μM
TP53:2 79.5 μM
TP63:3 101.0 μM
TPRX1:38 56.0 μM
TRAFD1:23 104.6 μM
TRERF1:31 54.0 μM
UBN1:24 52.7 μM
USP22:28 79.2 μM
ZMYM3:95 81.7 μM
ZNF200:6 230.4 μM
ZNF469:83 92.8 μM
ZNF644:90 89.6 μM
ZNF687:0 500.0 μM
rand:778 60.2 μM
rand:810 79.2 μM

P300 NCBD device 2:

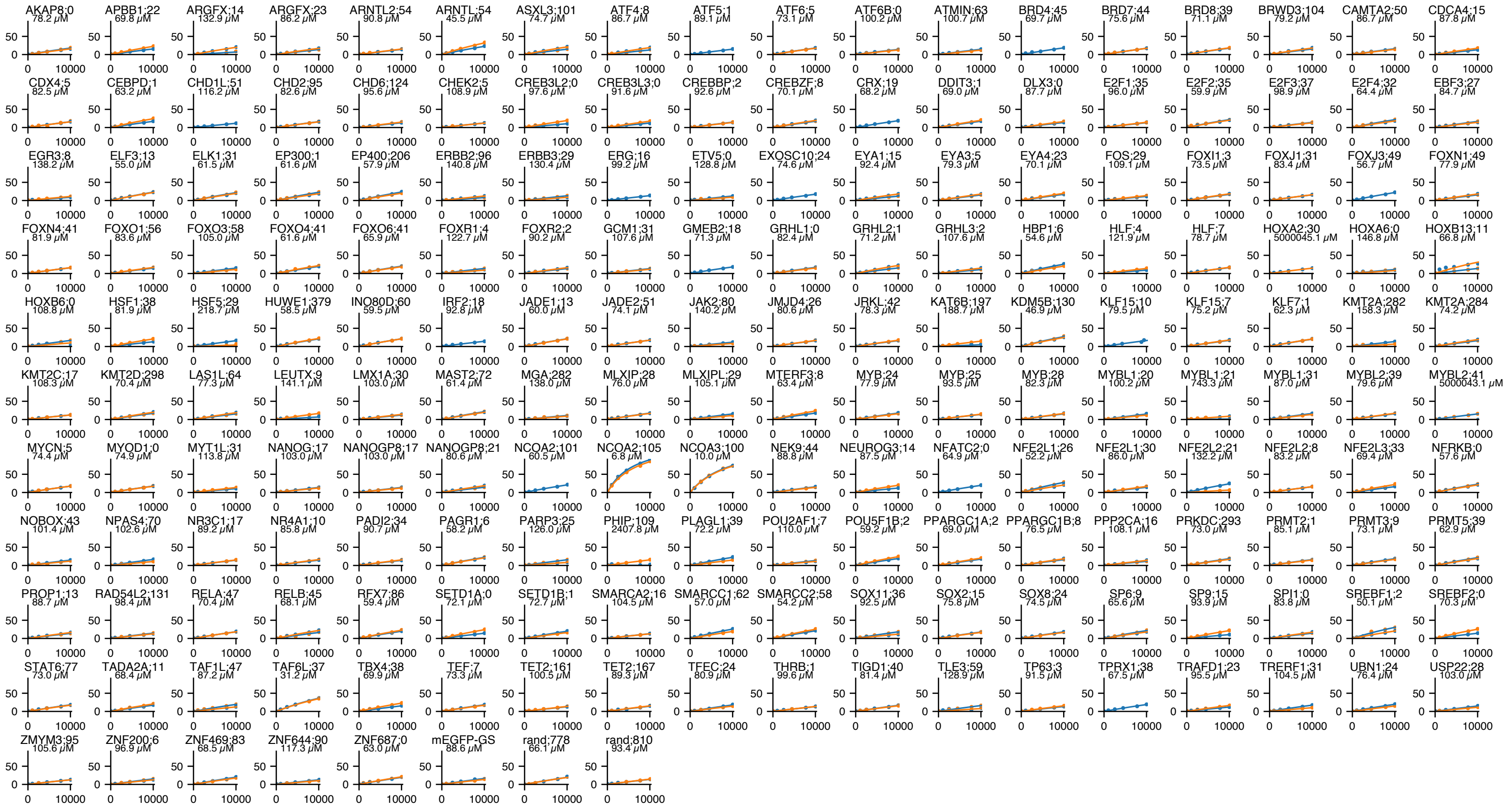


TAF6L device 1:

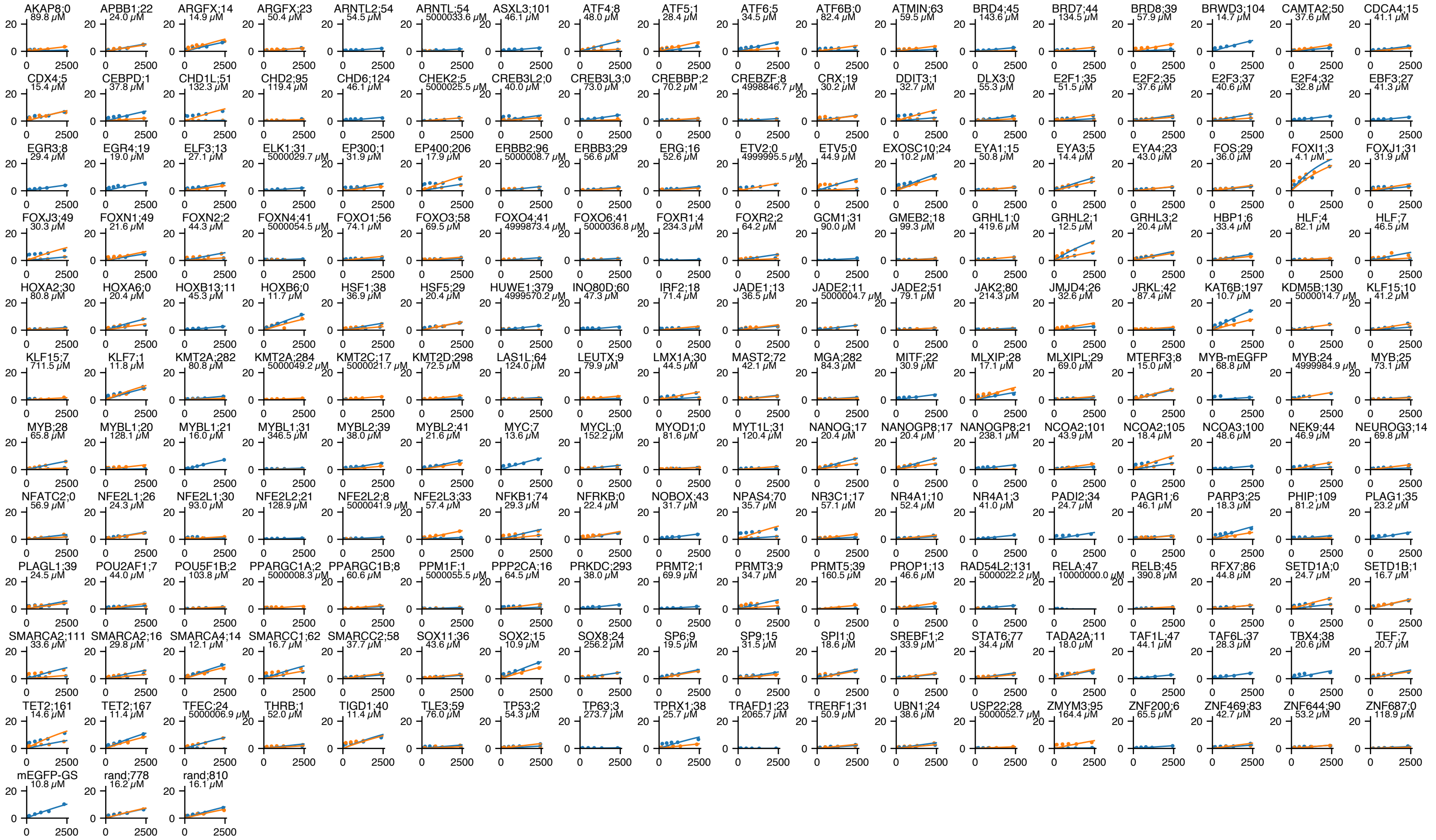

P300 TAZ1 device 1:

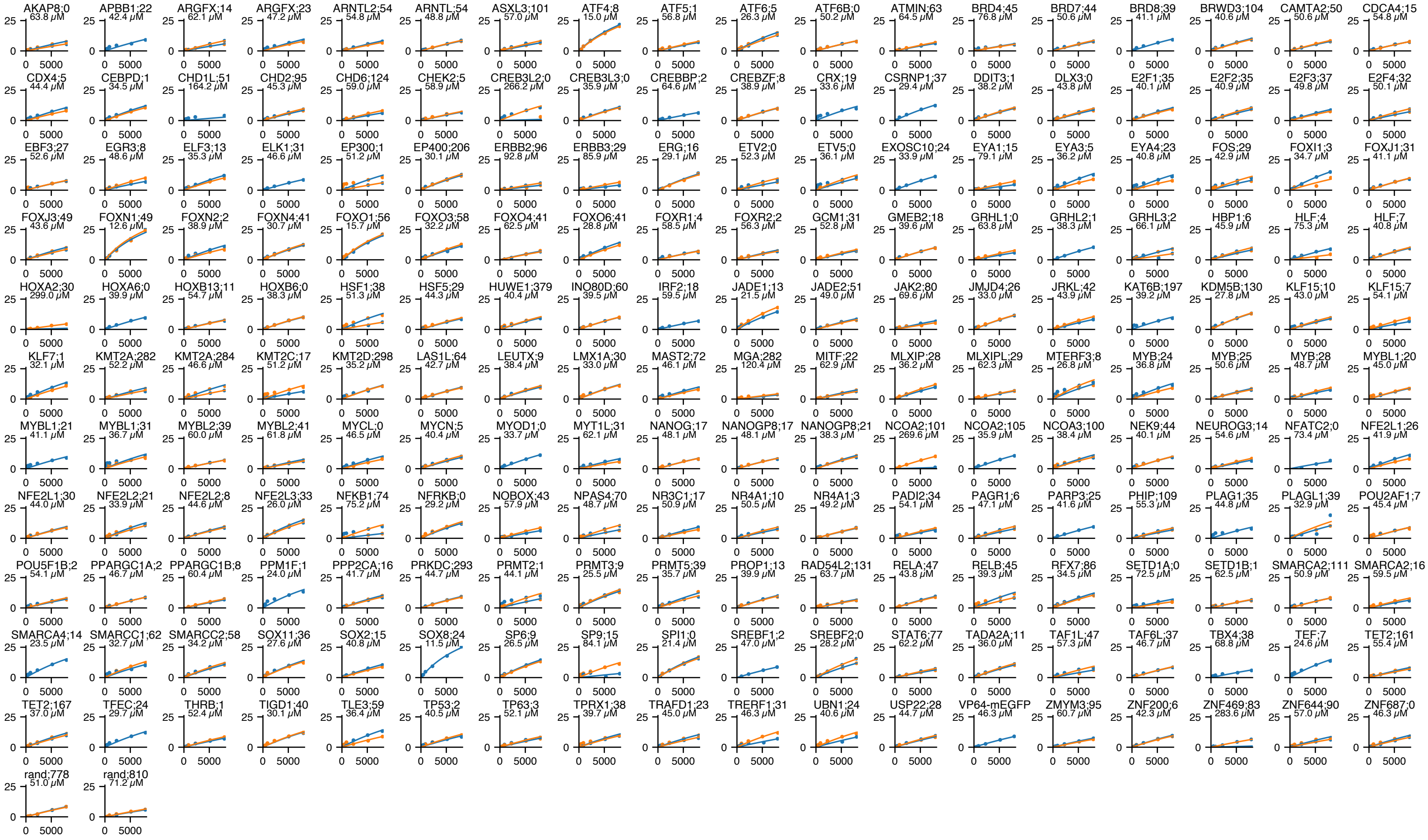


P300 TAZ1 device 2:

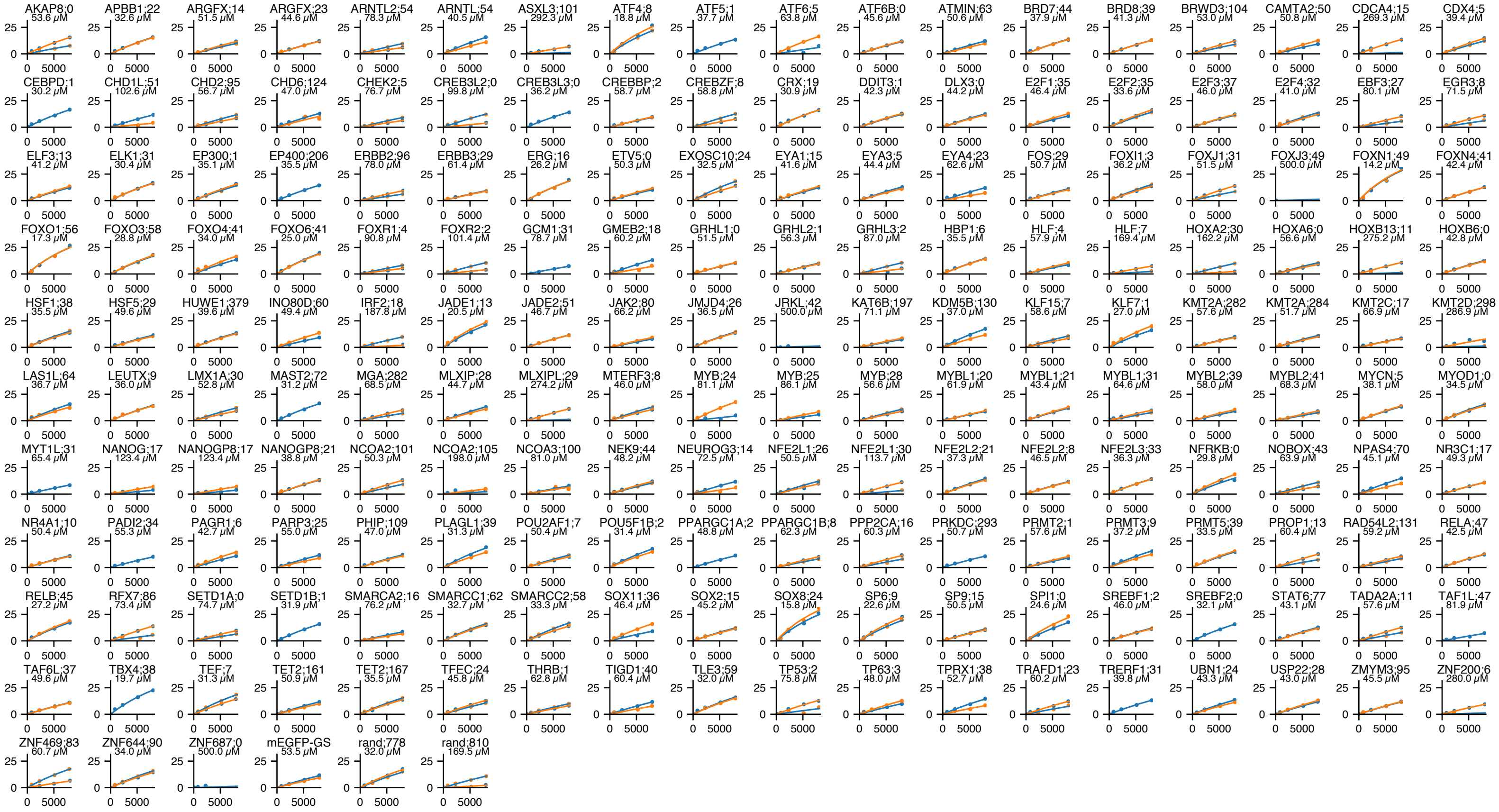

**e**

**Affinities replicability (red=meGFP):**

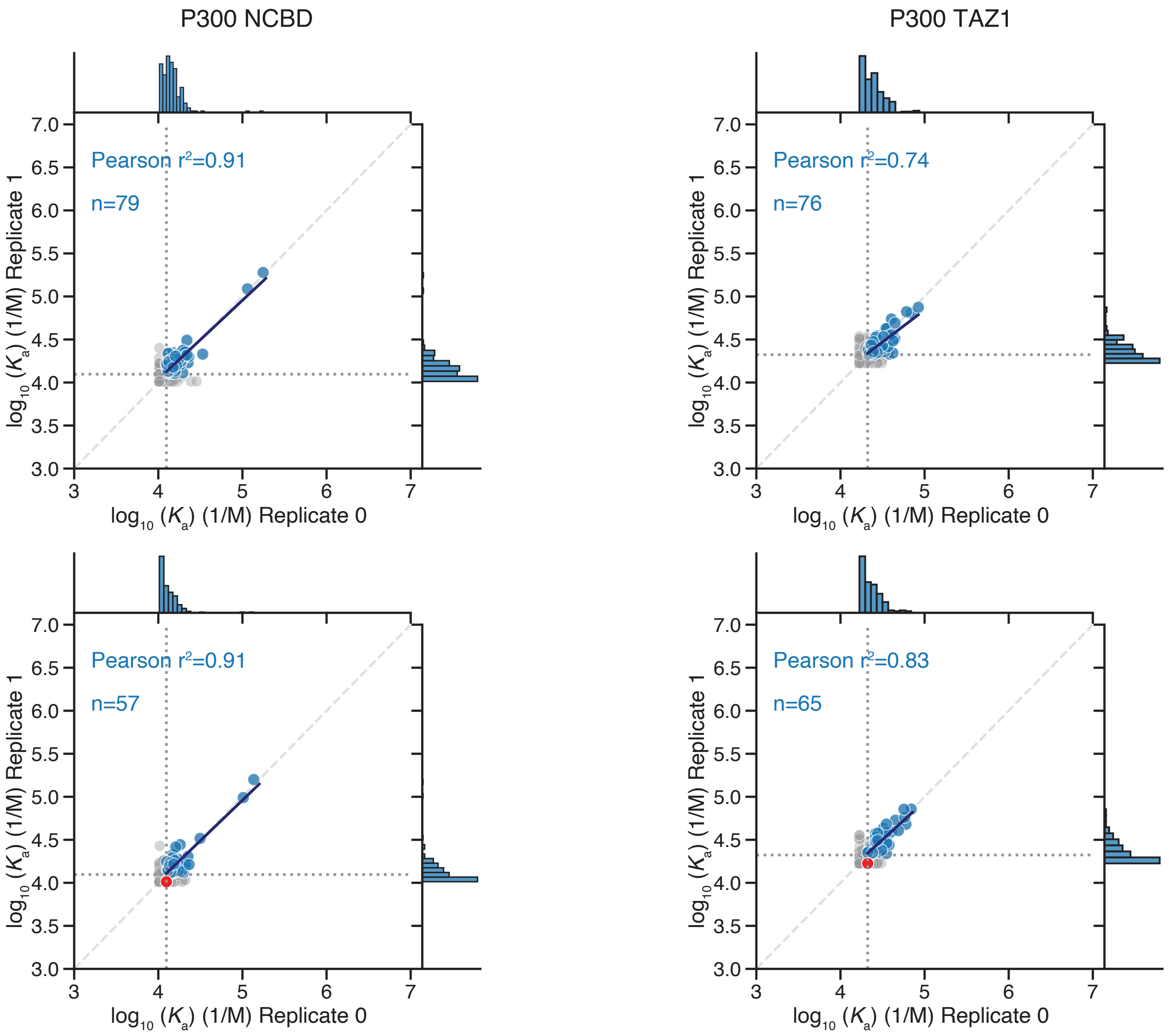


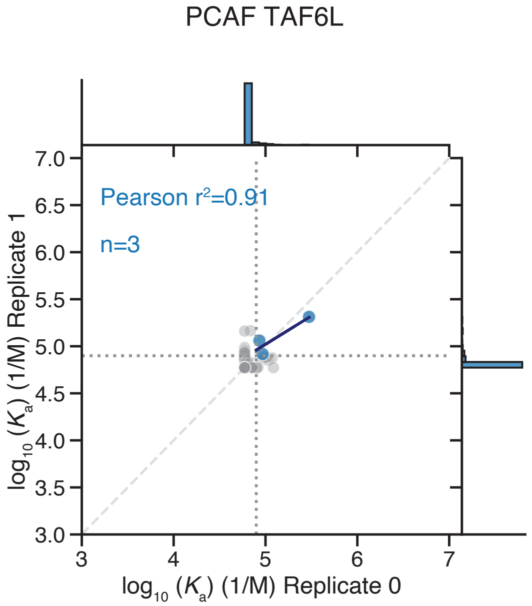


**f**

**Raw images at highest tested concentration co-A (top) AD (bottom)**

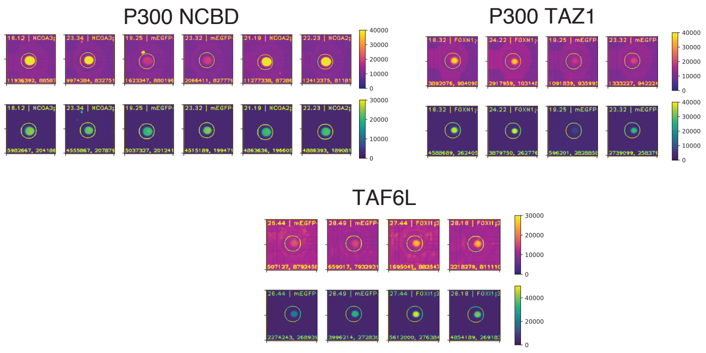

a

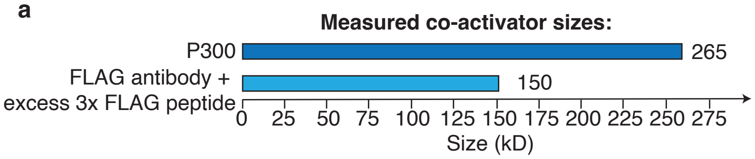


b

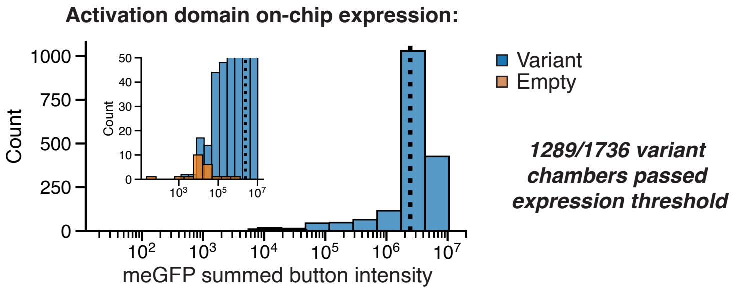


c

*All co-As were purchased for this experiment*

d

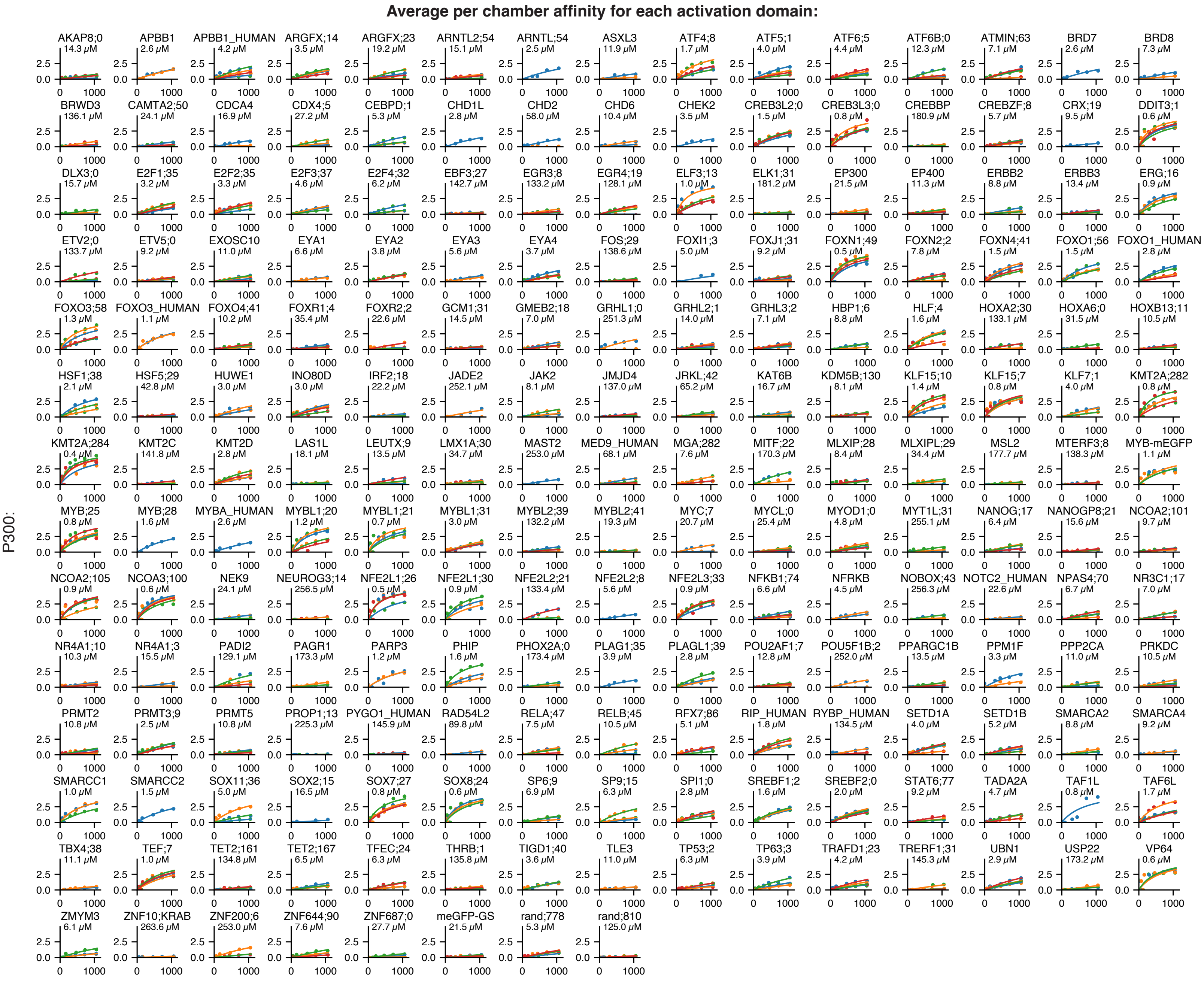

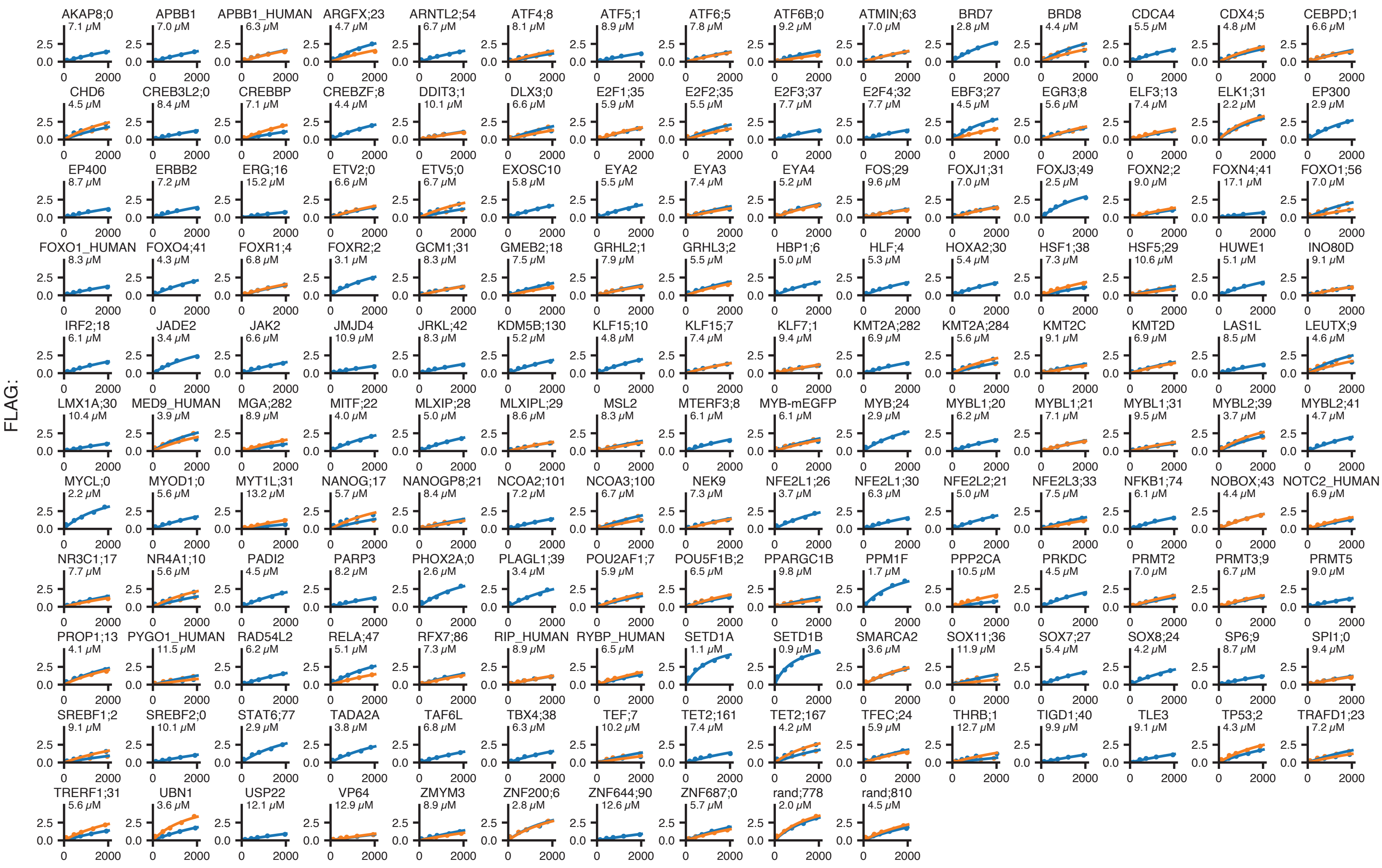


**e**

**Affinities replicability (red=meGFP):**

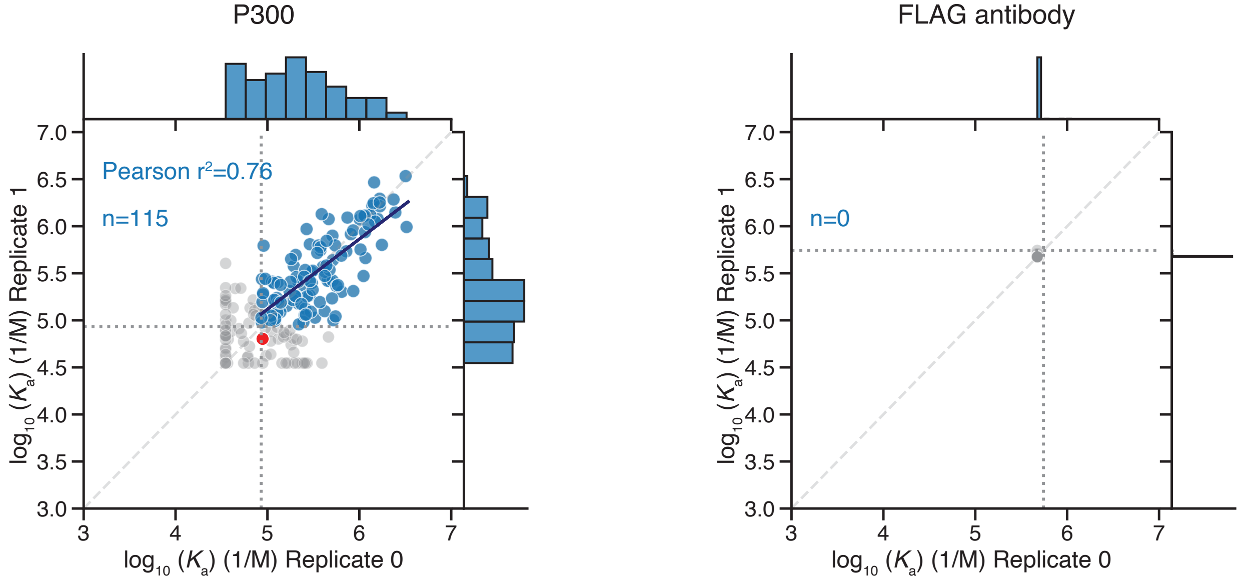


**f**

**Raw images at highest tested concentration, co-A (top) AD (bottom)**

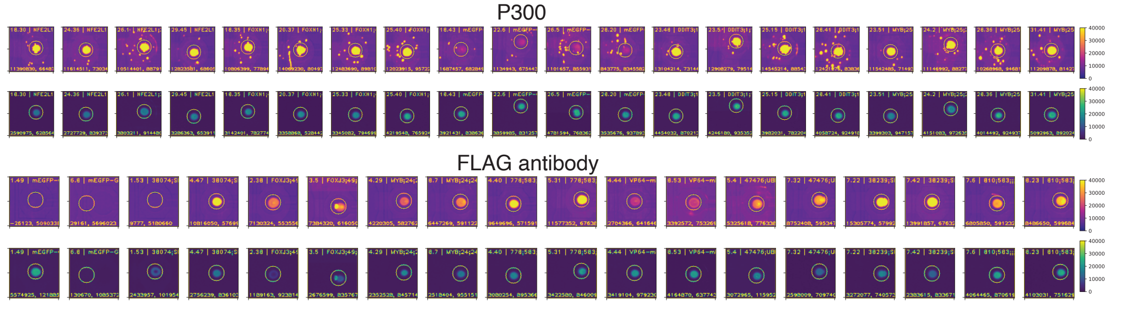

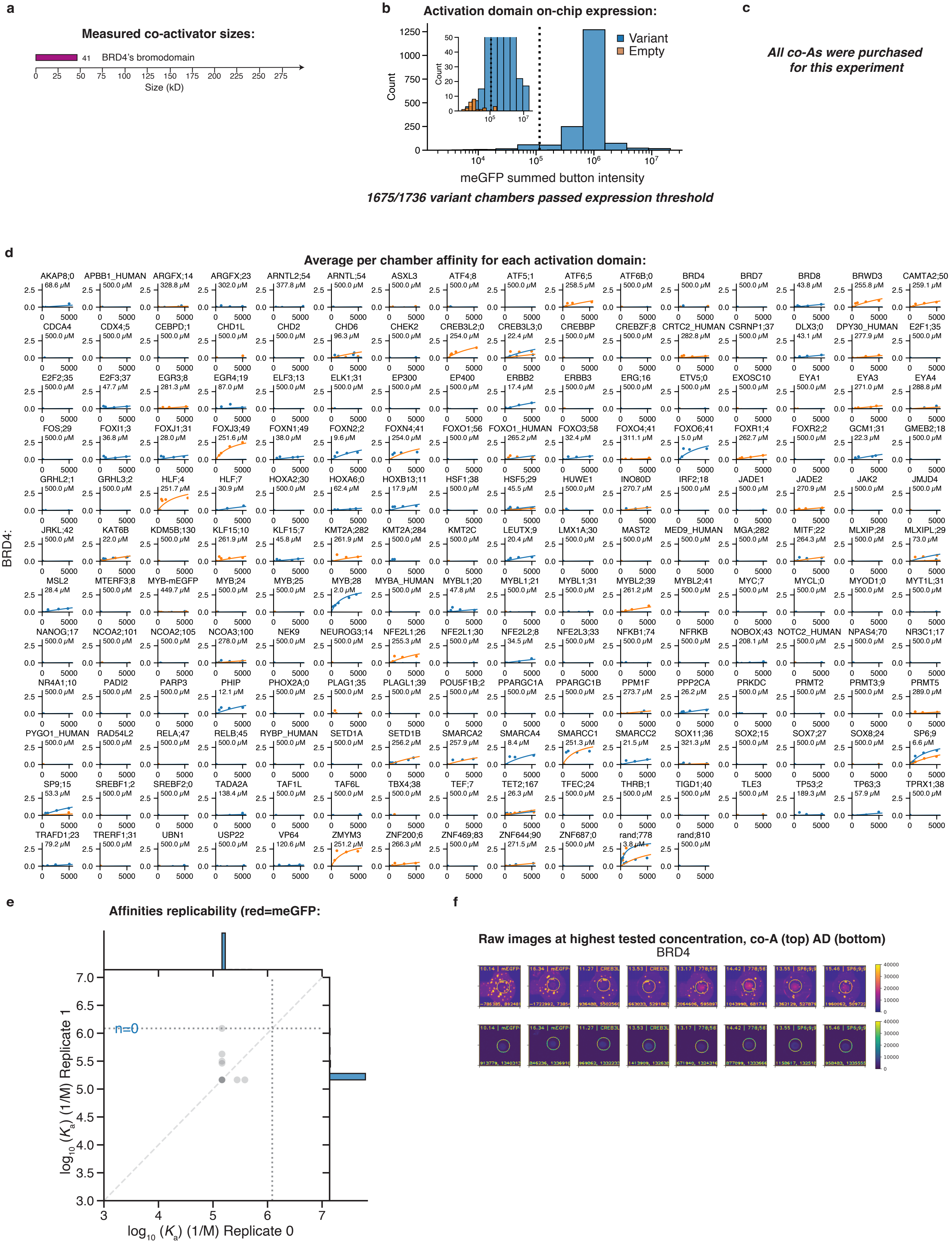
a
Measured co-activator sizes:
41 BRD4's bromodomain
Size (kD)
b
Activation domain on-chip expression:
Variant
Empty
Count
meGFP summed button intensity
1675/1736 variant chambers passed expression threshold
c
All co-As were purchased for this experiment
d
Average per chamber affinity for each activation domain:
BRD4:
e
Affinities replicability (red=meGFP:
n=0
$\log_{10}(K_a)$ (1/M) Replicate 1
$\log_{10}(K_a)$ (1/M) Replicate 0
f
Raw images at highest tested concentration, co-A (top) AD (bottom)
BRD4

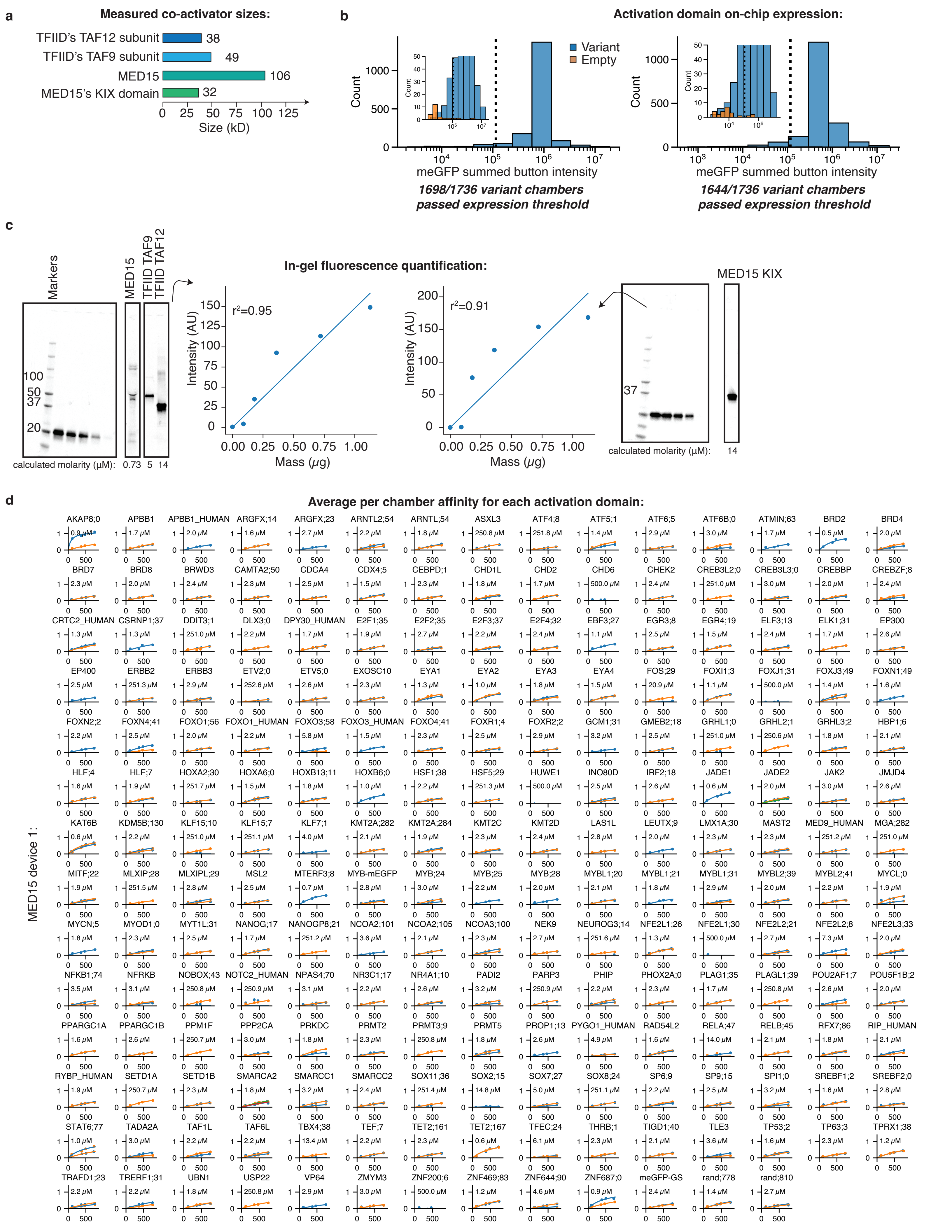

a
Measured co-activator sizes:
TFIID's TAF12 subunit 38
TFIID's TAF9 subunit 49
MED15 106
MED15's KIX domain 32
0 25 50 75 100 125
Size (kD)
b
Activation domain on-chip expression:
Variant
Empty
Count
meGFP summed button intensity
1698/1736 variant chambers passed expression threshold
1644/1736 variant chambers passed expression threshold
c
Markers
MED15
TFIID TAF9
TFIID TAF12
100
50
37
20
calculated molarity (μM): 0.73 5 14
In-gel fluorescence quantification:
r²=0.95
r²=0.91
Intensity (AU)
Mass (μg)
MED15 KIX
37
calculated molarity (μM): 14
d
Average per chamber affinity for each activation domain:
MED15 device 1:
AKAP8;0 0.9 μM
APBB1 1.7 μM
APBB1_HUMAN 2.0 μM
ARGFX;14 1.6 μM
ARGFX;23 2.7 μM
ARNTL2;54 2.2 μM
ARNTL;54 1.8 μM
ASXL3 250.8 μM
ATF4;8 251.8 μM
ATF5;1 1.4 μM
ATF6;5 2.9 μM
ATF6B;0 3.0 μM
ATMIN;63 1.7 μM
BRD2 0.5 μM
BRD4 2.0 μM
BRD7 2.3 μM
BRD8 2.0 μM
BRWD3 2.4 μM
CAMTA2;50 2.3 μM
CDCA4 2.5 μM
CDX4;5 1.5 μM
CEBPD;1 2.3 μM
CHD1L 1.8 μM
CHD2 1.7 μM
CHD6 500.0 μM
CHEK2 2.4 μM
CREB3L2;0 251.0 μM
CREB3L3;0 3.0 μM
CREBBP 2.0 μM
CREBZF;8 2.4 μM
CRTC2_HUMAN 1.3 μM
CSRNP1;37 1.3 μM
DDIT3;1 251.0 μM
DLX3;0 2.2 μM
DPY30_HUMAN 1.7 μM
E2F1;35 1.9 μM
E2F2;35 2.7 μM
E2F3;37 2.2 μM
E2F4;32 2.4 μM
EBF3;27 1.1 μM
EGR3;8 2.5 μM
EGR4;19 1.5 μM
ELF3;13 2.4 μM
ELK1;31 1.7 μM
EP300 2.6 μM
EP400 2.5 μM
ERBB2 251.3 μM
ERBB3 2.9 μM
ETV2;0 252.6 μM
ETV5;0 2.6 μM
EXOSC10 2.3 μM
EYA1 1.3 μM
EYA2 1.0 μM
EYA3 1.8 μM
EYA4 1.5 μM
FOS;29 20.9 μM
FOXI1;3 1.1 μM
FOXJ1;31 500.0 μM
FOXJ3;49 1.4 μM
FOXN1;49 1.6 μM
FOXN2;2 2.2 μM
FOXN4;41 2.5 μM
FOXO1;56 2.0 μM
FOXO1_HUMAN 2.2 μM
FOXO3;58 5.8 μM
FOXO3_HUMAN 1.5 μM
FOXO4;41 2.3 μM
FOXR1;4 2.5 μM
FOXR2;2 2.9 μM
GCM1;31 3.2 μM
GMEB2;18 2.5 μM
GRHL1;0 251.0 μM
GRHL2;1 250.6 μM
GRHL3;2 1.8 μM
HBP1;6 2.1 μM
HLF;4 1.6 μM
HLF;7 1.9 μM
HOXA2;30 251.7 μM
HOXA6;0 1.5 μM
HOXB13;11 1.8 μM
HOXB6;0 1.0 μM
HSF1;38 2.2 μM
HSF5;29 251.3 μM
HUWE1 500.0 μM
INO80D 2.5 μM
IRF2;18 2.6 μM
JADE1 0.6 μM
JADE2 2.0 μM
JAK2 3.0 μM
JMJD4 2.6 μM
KAT6B 0.6 μM
KDM5B;130 2.2 μM
KLF15;10 251.0 μM
KLF15;7 251.1 μM
KLF7;1 4.0 μM
KMT2A;282 2.1 μM
KMT2A;284 1.9 μM
KMT2C 2.3 μM
KMT2D 2.4 μM
LAS1L 2.8 μM
LEUTX;9 2.0 μM
LMX1A;30 2.3 μM
MAST2 2.3 μM
MED9_HUMAN 251.2 μM
MGA;282 251.0 μM
MITF;22 1.9 μM
MLXIP;28 251.5 μM
MLXIPL;29 2.8 μM
MSL2 2.5 μM
MTERF3;8 0.7 μM
MYB-mEGFP 2.8 μM
MYB;24 3.0 μM
MYB;25 2.2 μM
MYB;28 2.0 μM
MYBL1;20 2.1 μM
MYBL1;21 1.8 μM
MYBL1;31 2.9 μM
MYBL2;39 2.0 μM
MYBL2;41 2.2 μM
MYCL;0 1.9 μM
MYCN;5 1.8 μM
MYOD1;0 2.3 μM
MYT1L;31 2.5 μM
NANOG;17 1.7 μM
NANOGP8;21 251.2 μM
NCOA2;101 3.6 μM
NCOA2;105 2.1 μM
NCOA3;100 2.3 μM
NEK9 2.7 μM
NEUROG3;14 251.6 μM
NFE2L1;26 1.3 μM
NFE2L1;30 500.0 μM
NFE2L2;21 2.7 μM
NFE2L2;8 7.3 μM
NFE2L3;33 2.0 μM
NFKB1;74 3.5 μM
NFRKB 3.1 μM
NOBOX;43 250.8 μM
NOTC2_HUMAN 250.9 μM
NPAS4;70 3.1 μM
NR3C1;17 2.2 μM
NR4A1;10 2.6 μM
PADI2 3.2 μM
PARP3 250.9 μM
PHIP 2.2 μM
PHOX2A;0 2.3 μM
PLAG1;35 1.7 μM
PLAGL1;39 250.8 μM
POU2AF1;7 2.6 μM
POU5F1B;2 2.0 μM
PPARGC1A 1.6 μM
PPARGC1B 2.6 μM
PPM1F 250.7 μM
PPP2CA 3.0 μM
PRKDC 1.8 μM
PRMT2 2.3 μM
PRMT3;9 250.8 μM
PRMT5 1.8 μM
PROP1;13 2.6 μM
PYGO1_HUMAN 4.9 μM
RAD54L2 1.6 μM
RELA;47 14.0 μM
RELB;45 2.1 μM
RFX7;86 1.8 μM
RIP_HUMAN 2.1 μM
RYBP_HUMAN 1.9 μM
SETD1A 250.7 μM
SETD1B 2.3 μM
SMARCA2 1.8 μM
SMARCC1 3.2 μM
SMARCC2 2.4 μM
SOX11;36 251.4 μM
SOX2;15 14.8 μM
SOX7;27 5.0 μM
SOX8;24 251.1 μM
SP6;9 2.2 μM
SP9;15 2.5 μM
SPI1;0 3.2 μM
SREBF1;2 1.6 μM
SREBF2;0 2.8 μM
STAT6;77 1.0 μM
TADA2A 3.0 μM
TAF1L 2.2 μM
TAF6L 2.2 μM
TBX4;38 13.4 μM
TEF;7 2.2 μM
TET2;161 2.3 μM
TET2;167 0.6 μM
TFEC;24 6.1 μM
THRB;1 2.3 μM
TIGD1;40 2.1 μM
TLE3 3.6 μM
TP53;2 1.6 μM
TP63;3 2.3 μM
TPRX1;38 1.2 μM
TRAFD1;23 2.2 μM
TRERF1;31 2.2 μM
UBN1 1.8 μM
USP22 250.8 μM
VP64 2.9 μM
ZMYM3 3.0 μM
ZNF200;6 500.0 μM
ZNF469;83 1.2 μM
ZNF644;90 4.6 μM
ZNF687;0 0.9 μM
meGFP-GS 2.4 μM
rand;778 1.4 μM
rand;810 2.7 μM

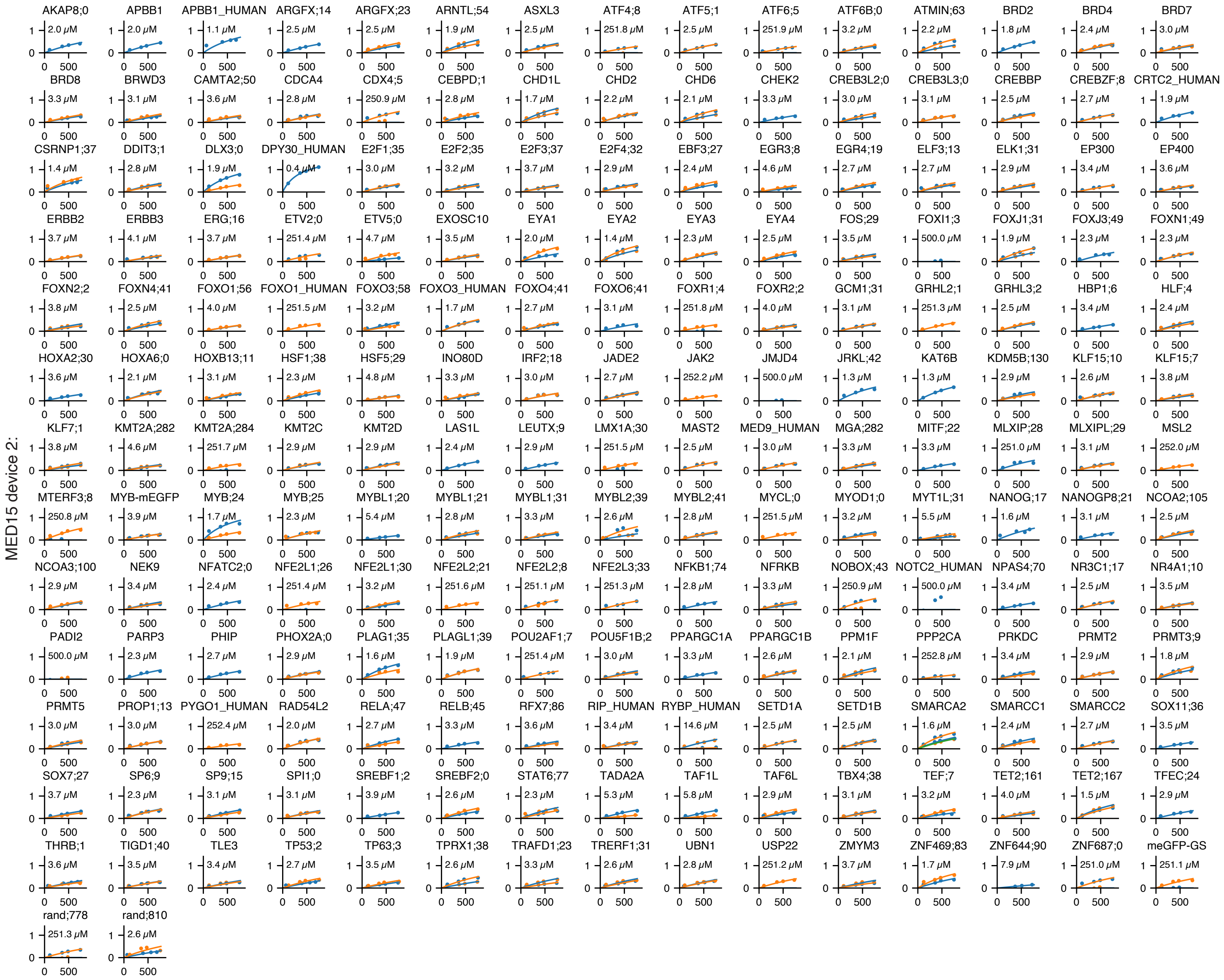
MED15 device 2:
AKAP8;0 2.0 μM
APBB1 2.0 μM
APBB1_HUMAN 1.1 μM
ARGFX;14 2.5 μM
ARGFX;23 2.5 μM
ARNTL;54 1.9 μM
ASXL3 2.5 μM
ATF4;8 251.8 μM
ATF5;1 2.5 μM
ATF6;5 251.9 μM
ATF6B;0 3.2 μM
ATMIN;63 2.2 μM
BRD2 1.8 μM
BRD4 2.4 μM
BRD7 3.0 μM
BRD8 3.3 μM
BRWD3 3.1 μM
CAMTA2;50 3.6 μM
CDCA4 2.8 μM
CDX4;5 250.9 μM
CEBPD;1 2.8 μM
CHD1L 1.7 μM
CHD2 2.2 μM
CHD6 2.1 μM
CHEK2 3.3 μM
CREB3L2;0 3.0 μM
CREB3L3;0 3.1 μM
CREBBP 2.5 μM
CREBZF;8 2.7 μM
CRTC2_HUMAN 1.9 μM
CSRNP1;37 1.4 μM
DDIT3;1 2.8 μM
DLX3;0 1.9 μM
DPY30_HUMAN 0.4 μM
E2F1;35 3.0 μM
E2F2;35 3.2 μM
E2F3;37 3.7 μM
E2F4;32 2.9 μM
EBF3;27 2.4 μM
EGR3;8 4.6 μM
EGR4;19 2.7 μM
ELF3;13 2.7 μM
ELK1;31 2.9 μM
EP300 3.4 μM
EP400 3.6 μM
ERBB2 3.7 μM
ERBB3 4.1 μM
ERG;16 3.7 μM
ETV2;0 251.4 μM
ETV5;0 4.7 μM
EXOSC10 3.5 μM
EYA1 2.0 μM
EYA2 1.4 μM
EYA3 2.3 μM
EYA4 2.5 μM
FOS;29 3.5 μM
FOXI1;3 500.0 μM
FOXJ1;31 1.9 μM
FOXJ3;49 2.3 μM
FOXN1;49 2.3 μM
FOXN2;2 3.8 μM
FOXN4;41 2.5 μM
FOXO1;56 4.0 μM
FOXO1_HUMAN 251.5 μM
FOXO3;58 3.2 μM
FOXO3_HUMAN 1.7 μM
FOXO4;41 2.7 μM
FOXO6;41 3.1 μM
FOXR1;4 251.8 μM
FOXR2;2 4.0 μM
GCM1;31 3.1 μM
GRHL2;1 251.3 μM
GRHL3;2 2.5 μM
HBP1;6 3.4 μM
HLF;4 2.4 μM
HOXA2;30 3.6 μM
HOXA6;0 2.1 μM
HOXB13;11 3.1 μM
HSF1;38 2.3 μM
HSF5;29 4.8 μM
INO80D 3.3 μM
IRF2;18 3.0 μM
JADE2 2.7 μM
JAK2 252.2 μM
JMJD4 500.0 μM
JRKL;42 1.3 μM
KAT6B 1.3 μM
KDM5B;130 2.9 μM
KLF15;10 2.6 μM
KLF15;7 3.8 μM
KLF7;1 3.8 μM
KMT2A;282 4.6 μM
KMT2A;284 251.7 μM
KMT2C 2.9 μM
KMT2D 2.9 μM
LAS1L 2.4 μM
LEUTX;9 2.9 μM
LMX1A;30 251.5 μM
MAST2 2.5 μM
MED9_HUMAN 3.0 μM
MGA;282 3.3 μM
MITF;22 3.3 μM
MLXIP;28 251.0 μM
MLXIPL;29 3.1 μM
MSL2 252.0 μM
MTERF3;8 250.8 μM
MYB-mEGFP 3.9 μM
MYB;24 1.7 μM
MYB;25 2.3 μM
MYBL1;20 5.4 μM
MYBL1;21 2.8 μM
MYBL1;31 3.3 μM
MYBL2;39 2.6 μM
MYBL2;41 2.8 μM
MYCL;0 251.5 μM
MYOD1;0 3.2 μM
MYT1L;31 5.5 μM
NANOG;17 1.6 μM
NANOGP8;21 3.1 μM
NCOA2;105 2.5 μM
NCOA3;100 2.9 μM
NEK9 3.4 μM
NFATC2;0 2.4 μM
NFE2L1;26 251.4 μM
NFE2L1;30 3.2 μM
NFE2L2;21 251.6 μM
NFE2L2;8 251.1 μM
NFE2L3;33 251.3 μM
NFKB1;74 2.8 μM
NFRKB 3.3 μM
NOBOX;43 250.9 μM
NOTC2_HUMAN 500.0 μM
NPAS4;70 3.4 μM
NR3C1;17 2.5 μM
NR4A1;10 3.5 μM
PADI2 500.0 μM
PARP3 2.3 μM
PHIP 2.7 μM
PHOX2A;0 2.9 μM
PLAG1;35 1.6 μM
PLAGL1;39 1.9 μM
POU2AF1;7 251.4 μM
POU5F1B;2 3.0 μM
PPARGC1A 3.3 μM
PPARGC1B 2.6 μM
PPM1F 2.1 μM
PPP2CA 252.8 μM
PRKDC 3.4 μM
PRMT2 2.9 μM
PRMT3;9 1.8 μM
PRMT5 3.0 μM
PROP1;13 3.0 μM
PYGO1_HUMAN 252.4 μM
RAD54L2 2.0 μM
RELA;47 2.7 μM
RELB;45 3.3 μM
RFX7;86 3.6 μM
RIP_HUMAN 3.4 μM
RYBP_HUMAN 14.6 μM
SETD1A 2.5 μM
SETD1B 2.5 μM
SMARCA2 1.6 μM
SMARCC1 2.4 μM
SMARCC2 2.7 μM
SOX11;36 3.5 μM
SOX7;27 3.7 μM
SP6;9 2.3 μM
SP9;15 3.1 μM
SPI1;0 3.1 μM
SREBF1;2 3.9 μM
SREBF2;0 2.6 μM
STAT6;77 2.3 μM
TADA2A 5.3 μM
TAF1L 5.8 μM
TAF6L 2.9 μM
TBX4;38 3.1 μM
TEF;7 3.2 μM
TET2;161 4.0 μM
TET2;167 1.5 μM
TFEC;24 2.9 μM
THRB;1 3.6 μM
TIGD1;40 3.5 μM
TLE3 3.4 μM
TP53;2 2.7 μM
TP63;3 3.5 μM
TPRX1;38 2.6 μM
TRAFD1;23 3.3 μM
TRERF1;31 2.6 μM
UBN1 2.8 μM
USP22 251.2 μM
ZMYM3 3.7 μM
ZNF469;83 1.7 μM
ZNF644;90 7.9 μM
ZNF687;0 251.0 μM
meGFP-GS 251.1 μM
rand;778 251.3 μM
rand;810 2.6 μM

MED15 KIX device 1:

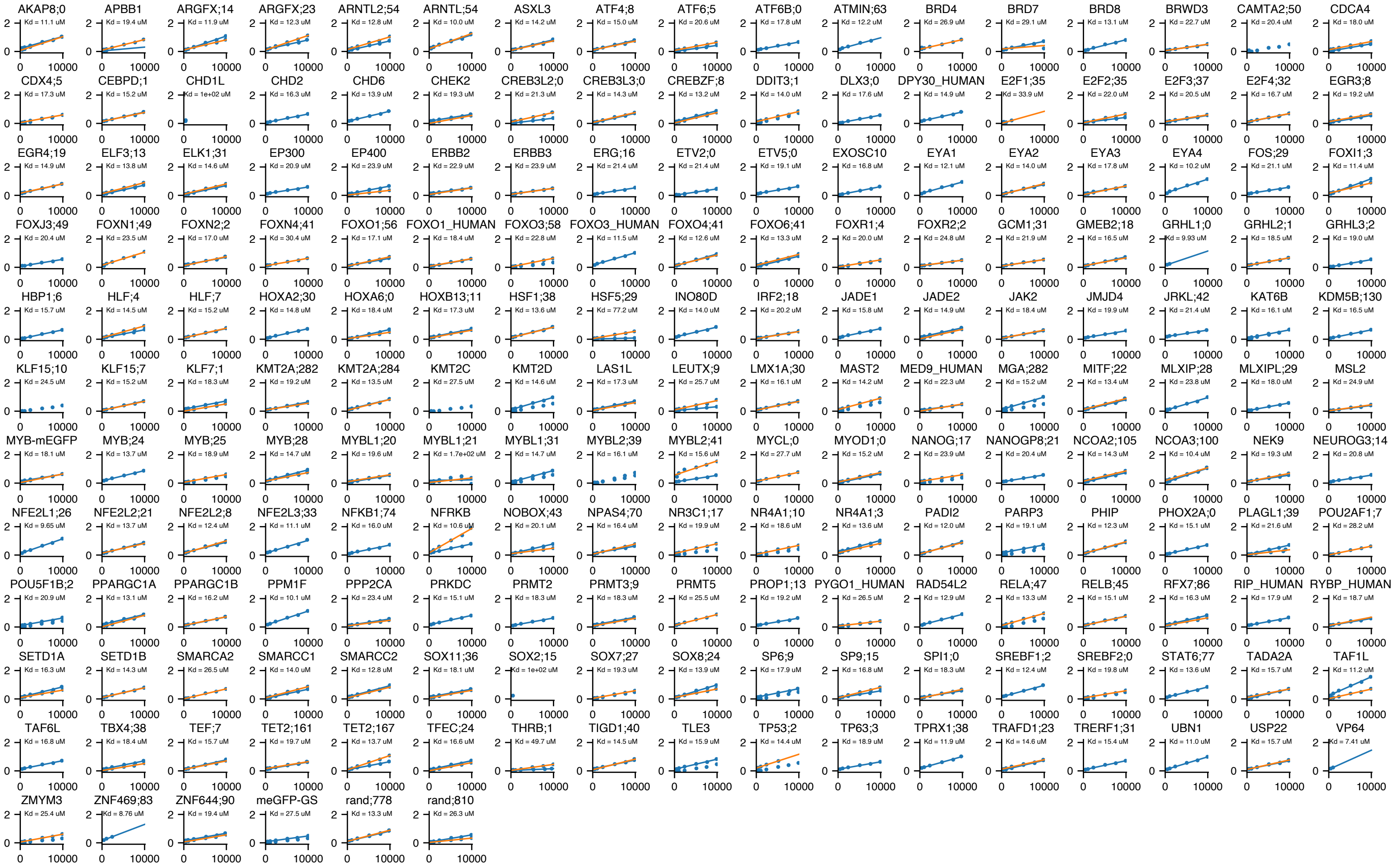


MED15 KIX device 2:

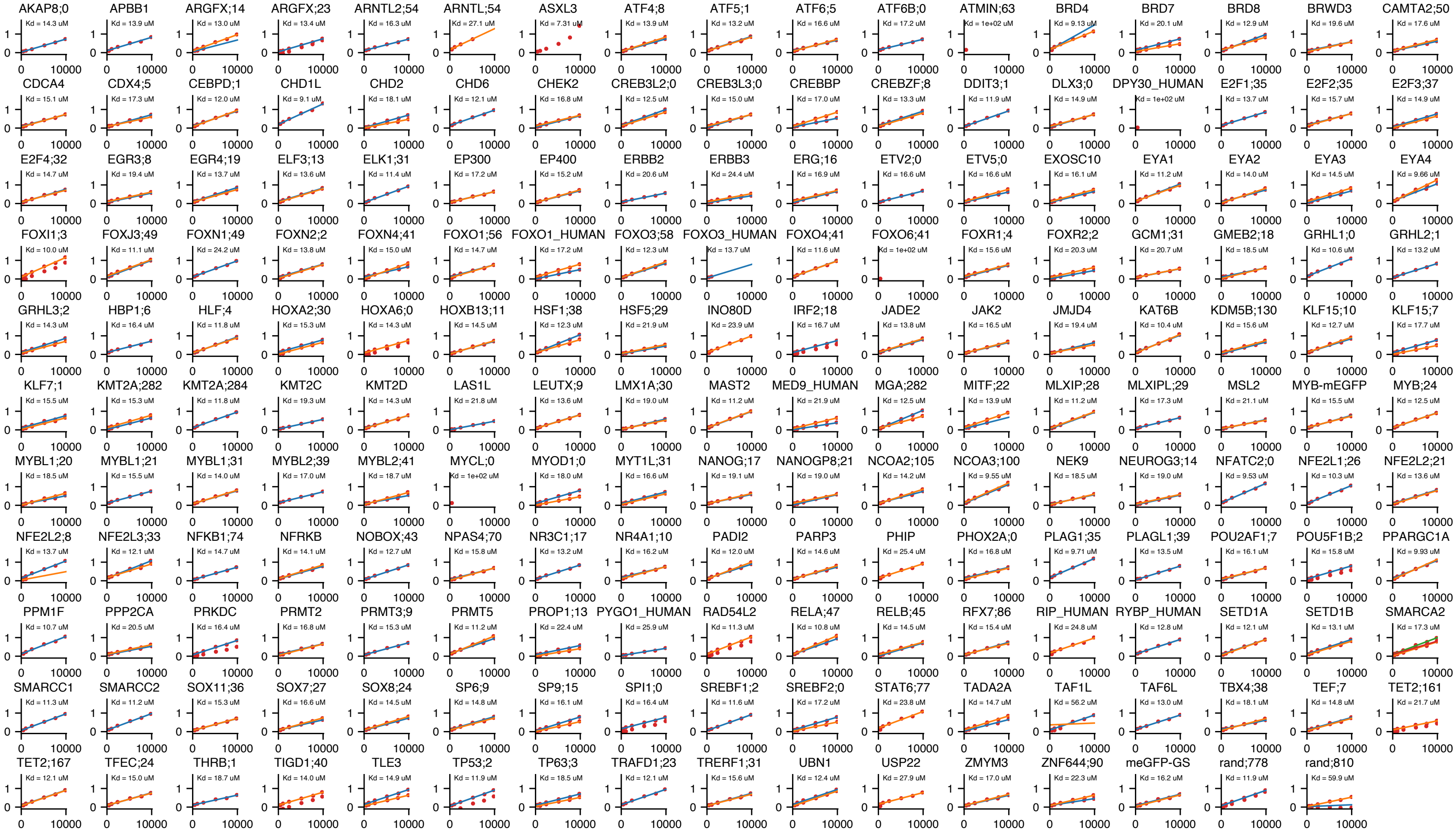

TFIID TAF12 device 1:

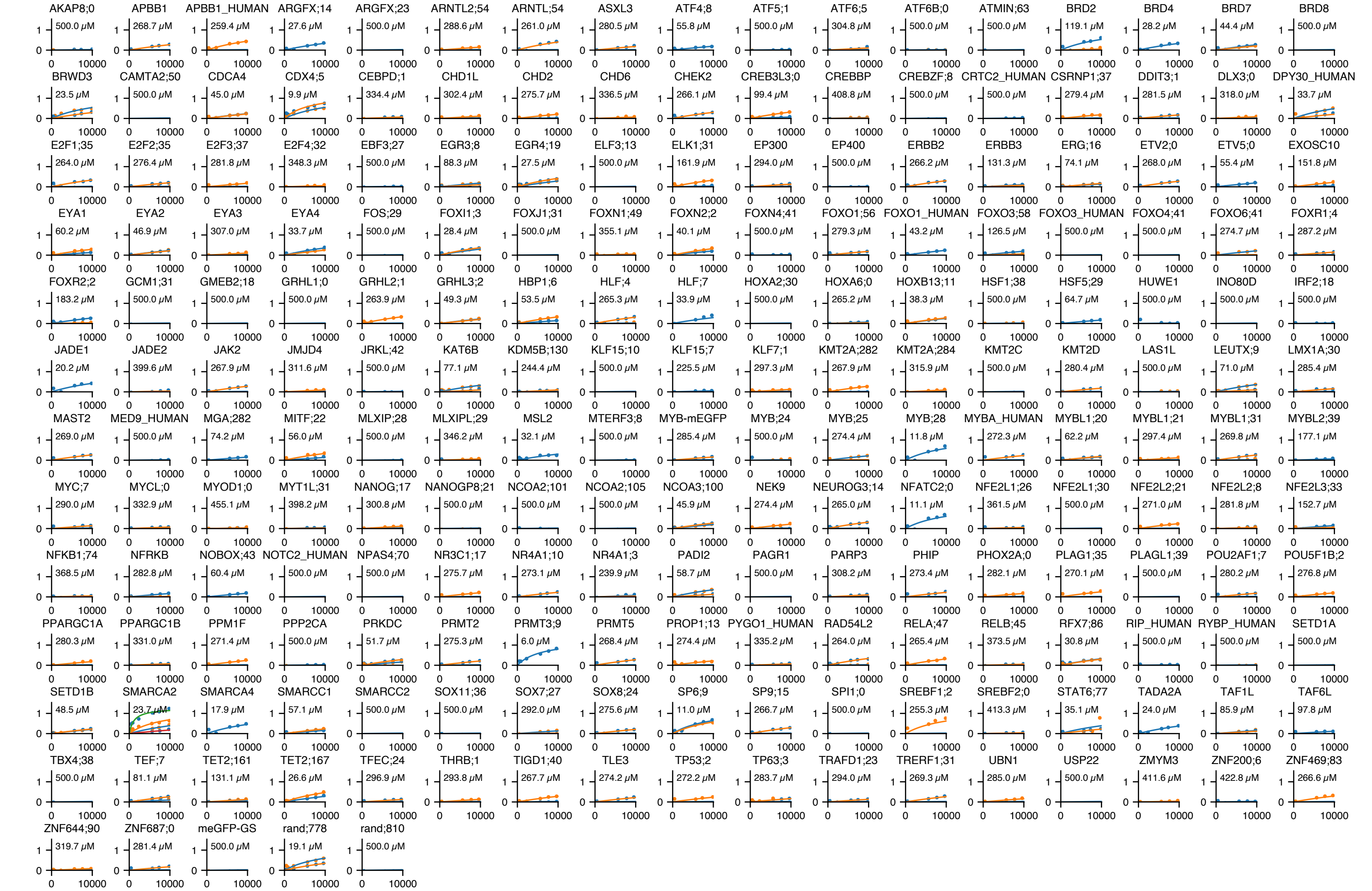


TFIID TAF12 device 2:

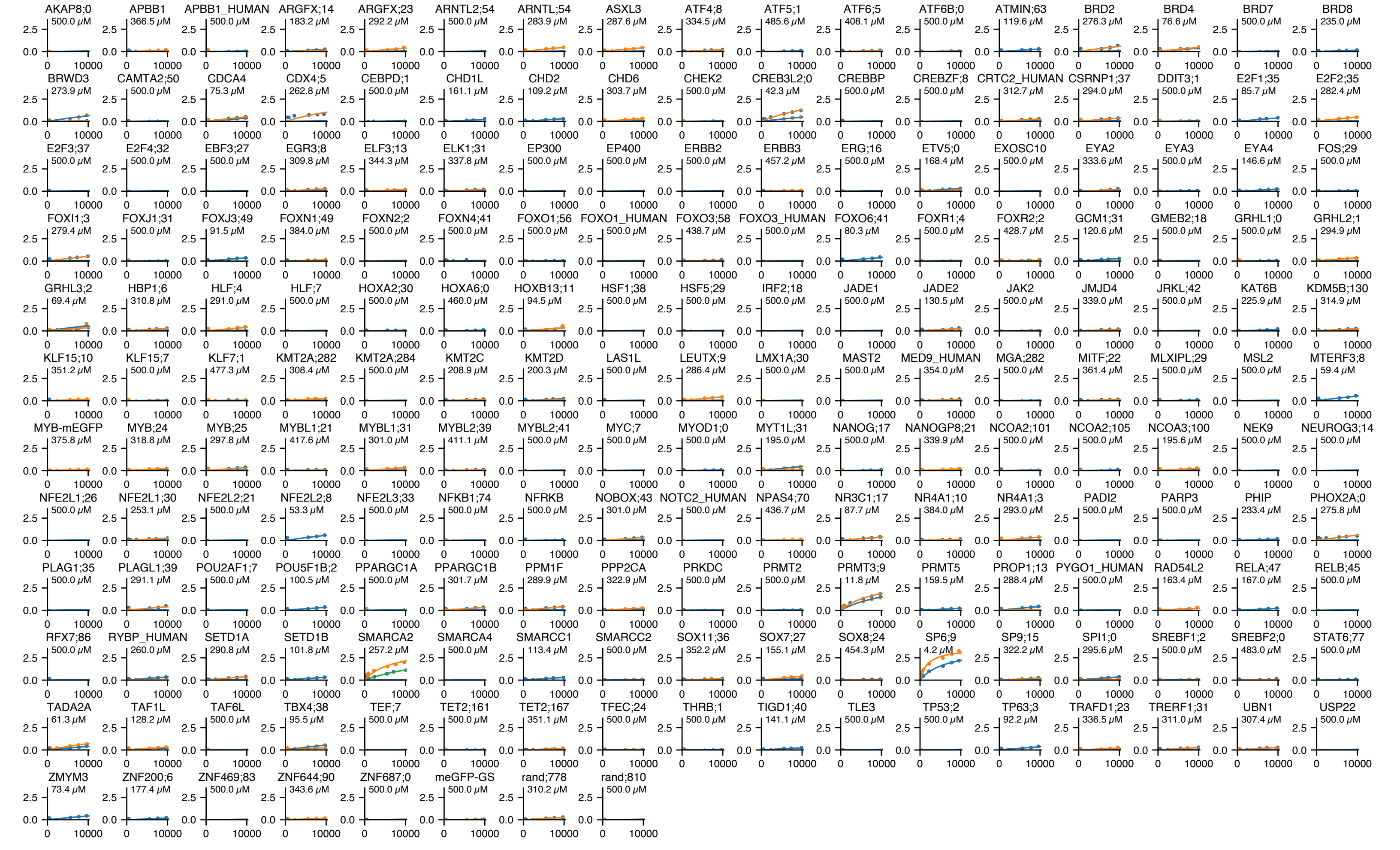

e

**Affinities replicability (red=meGFP, TFIID TAF12 is random control;778):**

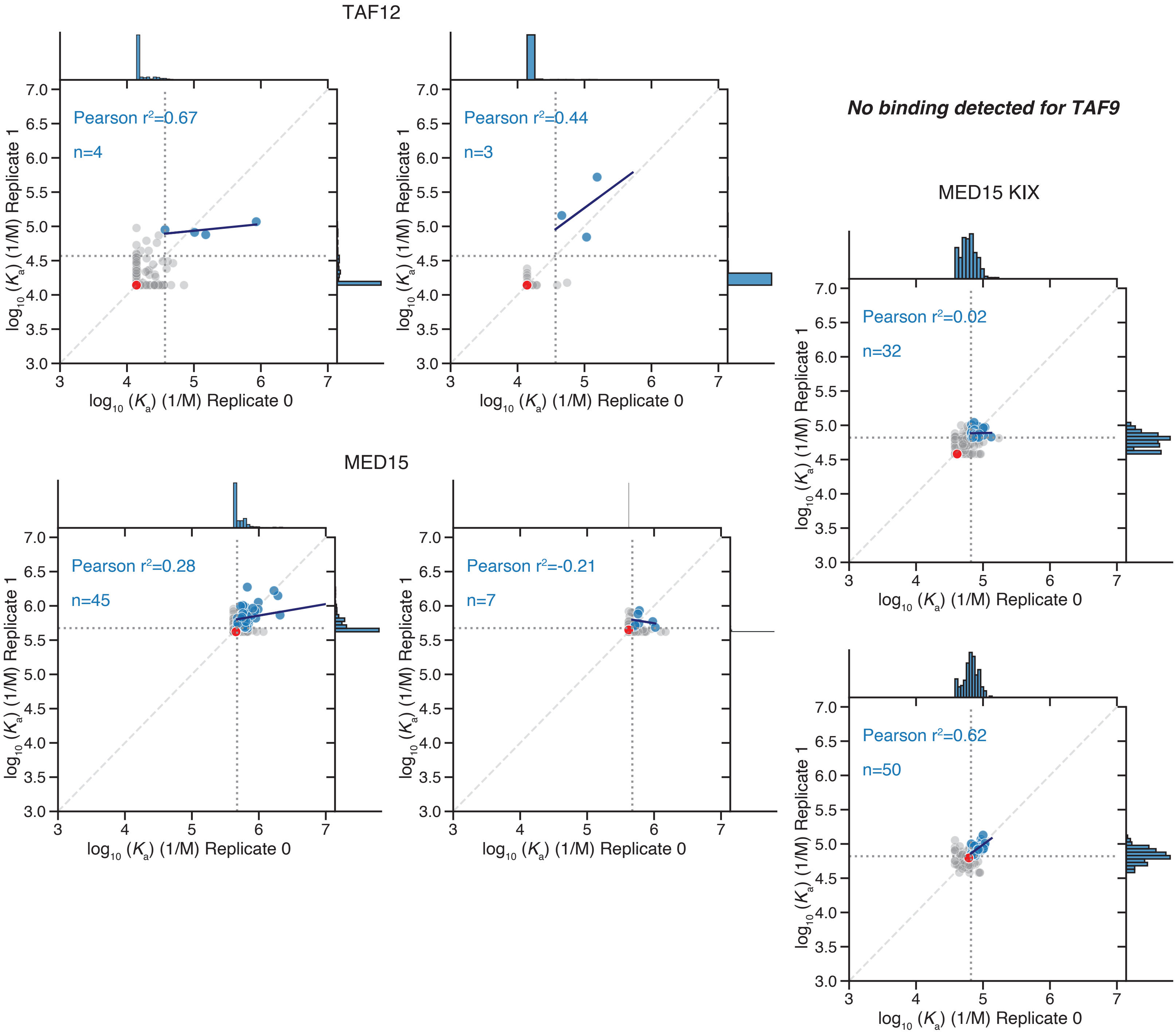


f

**Raw images at highest tested concentration, co-A (top) AD (bottom)**

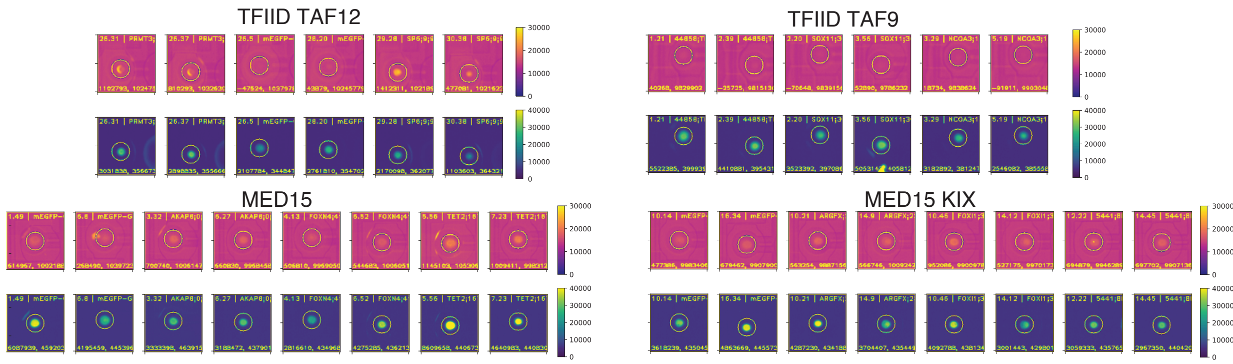

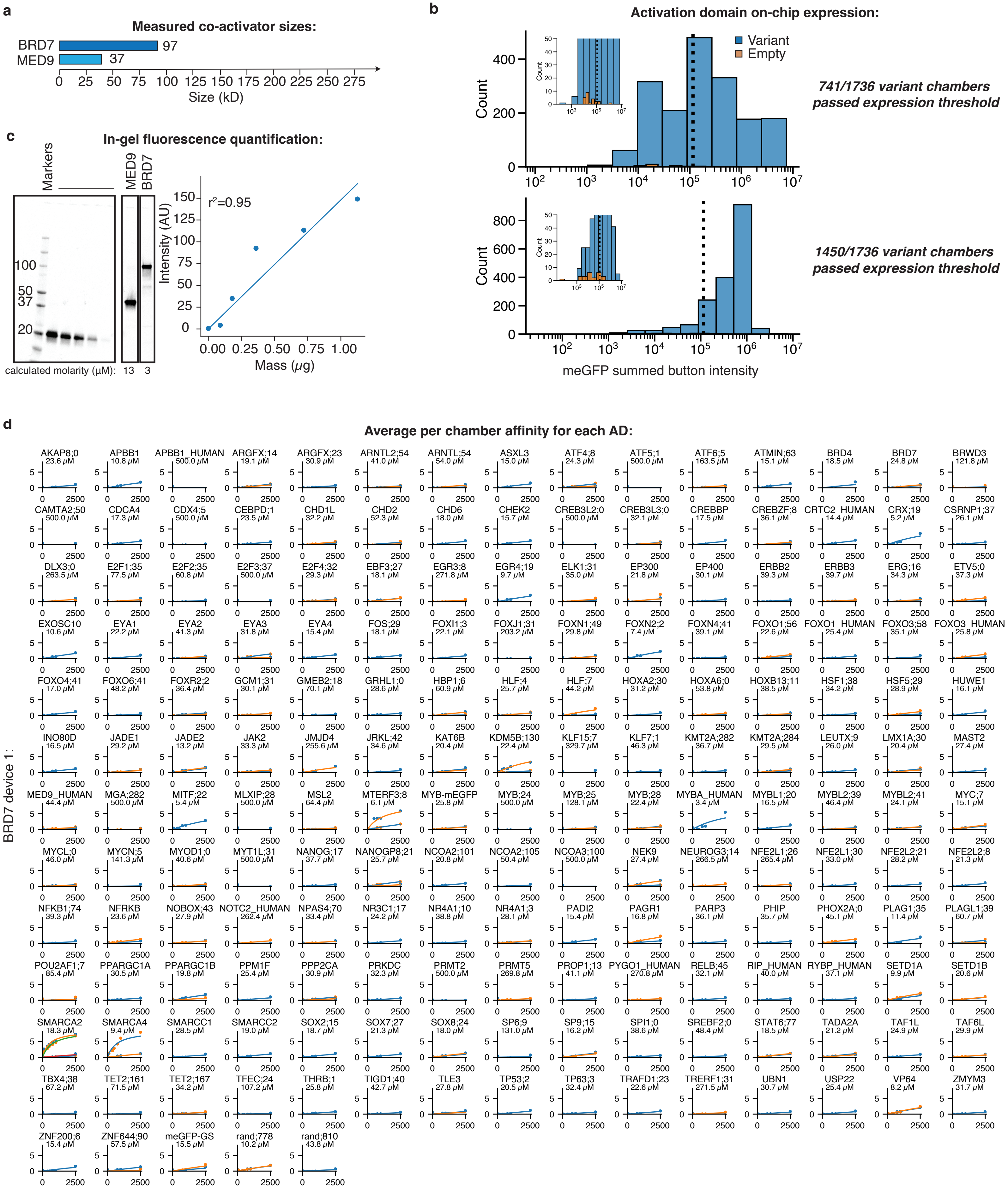

a
Measured co-activator sizes:
BRD7 97
MED9 37
Size (kD)
b
Activation domain on-chip expression:
Variant
Empty
Count
741/1736 variant chambers passed expression threshold
1450/1736 variant chambers passed expression threshold
meGFP summed button intensity
c
In-gel fluorescence quantification:
Markers
MED9
BRD7
100
50
37
20
calculated molarity (μM): 13 3
r²=0.95
Intensity (AU)
Mass (μg)
d
Average per chamber affinity for each AD:
BRD7 device 1:
AKAP8;0 23.6 μM
APBB1 10.8 μM
APBB1_HUMAN 500.0 μM
ARGFX;14 19.1 μM
ARGFX;23 30.9 μM
ARNTL2;54 41.0 μM
ARNTL;54 54.0 μM
ASXL3 15.0 μM
ATF4;8 24.3 μM
ATF5;1 500.0 μM
ATF6;5 163.5 μM
ATMIN;63 15.1 μM
BRD4 18.5 μM
BRD7 24.8 μM
BRWD3 121.8 μM
CAMTA2;50 500.0 μM
CDCA4 17.3 μM
CDX4;5 500.0 μM
CEBPD;1 23.5 μM
CHD1L 32.2 μM
CHD2 52.3 μM
CHD6 18.0 μM
CHEK2 15.7 μM
CREB3L2;0 500.0 μM
CREB3L3;0 32.1 μM
CREBBP 17.5 μM
CREBZF;8 36.1 μM
CRTC2_HUMAN 14.4 μM
CRX;19 5.2 μM
CSRNP1;37 26.1 μM
DLX3;0 263.5 μM
E2F1;35 77.5 μM
E2F2;35 60.8 μM
E2F3;37 500.0 μM
E2F4;32 29.3 μM
EBF3;27 18.1 μM
EGR3;8 271.8 μM
EGR4;19 9.7 μM
ELK1;31 35.0 μM
EP300 21.8 μM
EP400 30.1 μM
ERBB2 39.3 μM
ERBB3 39.7 μM
ERG;16 34.3 μM
ETV5;0 37.3 μM
EXOSC10 10.6 μM
EYA1 22.2 μM
EYA2 41.3 μM
EYA3 31.8 μM
EYA4 15.4 μM
FOS;29 18.1 μM
FOXI1;3 22.1 μM
FOXJ1;31 203.2 μM
FOXN1;49 29.8 μM
FOXN2;2 7.4 μM
FOXN4;41 39.1 μM
FOXO1;56 22.6 μM
FOXO1_HUMAN 25.4 μM
FOXO3;58 35.1 μM
FOXO3_HUMAN 25.8 μM
FOXO4;41 17.0 μM
FOXO6;41 48.2 μM
FOXR2;2 36.4 μM
GCM1;31 30.1 μM
GMEB2;18 70.1 μM
GRHL1;0 28.6 μM
HBP1;6 60.9 μM
HLF;4 25.7 μM
HLF;7 44.2 μM
HOXA2;30 31.2 μM
HOXA6;0 53.8 μM
HOXB13;11 38.5 μM
HSF1;38 34.2 μM
HSF5;29 28.9 μM
HUWE1 16.1 μM
INO80D 16.5 μM
JADE1 29.2 μM
JADE2 13.2 μM
JAK2 33.3 μM
JMJD4 255.6 μM
JRKL;42 34.6 μM
KAT6B 20.4 μM
KDM5B;130 22.4 μM
KLF15;7 329.7 μM
KLF7;1 46.3 μM
KMT2A;282 36.7 μM
KMT2A;284 29.5 μM
LEUTX;9 26.0 μM
LMX1A;30 20.4 μM
MAST2 27.4 μM
MED9_HUMAN 44.4 μM
MGA;282 500.0 μM
MITF;22 5.4 μM
MLXIP;28 500.0 μM
MSL2 64.4 μM
MTERF3;8 6.1 μM
MYB-mEGFP 25.8 μM
MYB;24 500.0 μM
MYB;25 128.1 μM
MYB;28 22.4 μM
MYBA_HUMAN 3.4 μM
MYBL1;20 16.5 μM
MYBL2;39 46.4 μM
MYBL2;41 24.1 μM
MYC;7 15.1 μM
MYCL;0 46.0 μM
MYCN;5 141.3 μM
MYOD1;0 40.6 μM
MYT1L;31 500.0 μM
NANOG;17 37.7 μM
NANOGP8;21 25.7 μM
NCOA2;101 20.8 μM
NCOA2;105 50.4 μM
NCOA3;100 500.0 μM
NEK9 27.4 μM
NEUROG3;14 266.5 μM
NFE2L1;26 265.4 μM
NFE2L1;30 33.0 μM
NFE2L2;21 28.2 μM
NFE2L2;8 21.3 μM
NFKB1;74 39.3 μM
NFRKB 23.6 μM
NOBOX;43 27.9 μM
NOTC2_HUMAN 262.4 μM
NPAS4;70 33.4 μM
NR3C1;17 24.2 μM
NR4A1;10 38.8 μM
NR4A1;3 28.1 μM
PADI2 15.4 μM
PAGR1 16.8 μM
PARP3 36.1 μM
PHIP 35.7 μM
PHOX2A;0 45.1 μM
PLAG1;35 11.4 μM
PLAGL1;39 60.7 μM
POU2AF1;7 85.4 μM
PPARGC1A 30.5 μM
PPARGC1B 19.8 μM
PPM1F 25.4 μM
PPP2CA 30.9 μM
PRKDC 32.3 μM
PRMT2 500.0 μM
PRMT5 269.8 μM
PROP1;13 41.1 μM
PYGO1_HUMAN 270.8 μM
RELB;45 32.1 μM
RIP_HUMAN 40.0 μM
RYBP_HUMAN 37.1 μM
SETD1A 9.9 μM
SETD1B 20.6 μM
SMARCA2 18.3 μM
SMARCA4 9.4 μM
SMARCC1 28.5 μM
SMARCC2 19.0 μM
SOX2;15 18.7 μM
SOX7;27 21.3 μM
SOX8;24 18.0 μM
SP6;9 131.0 μM
SP9;15 16.2 μM
SPI1;0 38.6 μM
SREBF2;0 48.4 μM
STAT6;77 18.5 μM
TADA2A 21.2 μM
TAF1L 24.9 μM
TAF6L 29.9 μM
TBX4;38 67.2 μM
TET2;161 71.5 μM
TET2;167 34.2 μM
TFEC;24 107.2 μM
THRB;1 25.8 μM
TIGD1;40 42.7 μM
TLE3 27.8 μM
TP53;2 20.5 μM
TP63;3 32.4 μM
TRAFD1;23 22.6 μM
TRERF1;31 271.5 μM
UBN1 30.7 μM
USP22 25.4 μM
VP64 8.2 μM
ZMYM3 31.7 μM
ZNF200;6 15.4 μM
ZNF644;90 57.5 μM
meGFP-GS 15.5 μM
rand;778 10.2 μM
rand;810 43.8 μM

MED9 device 1:

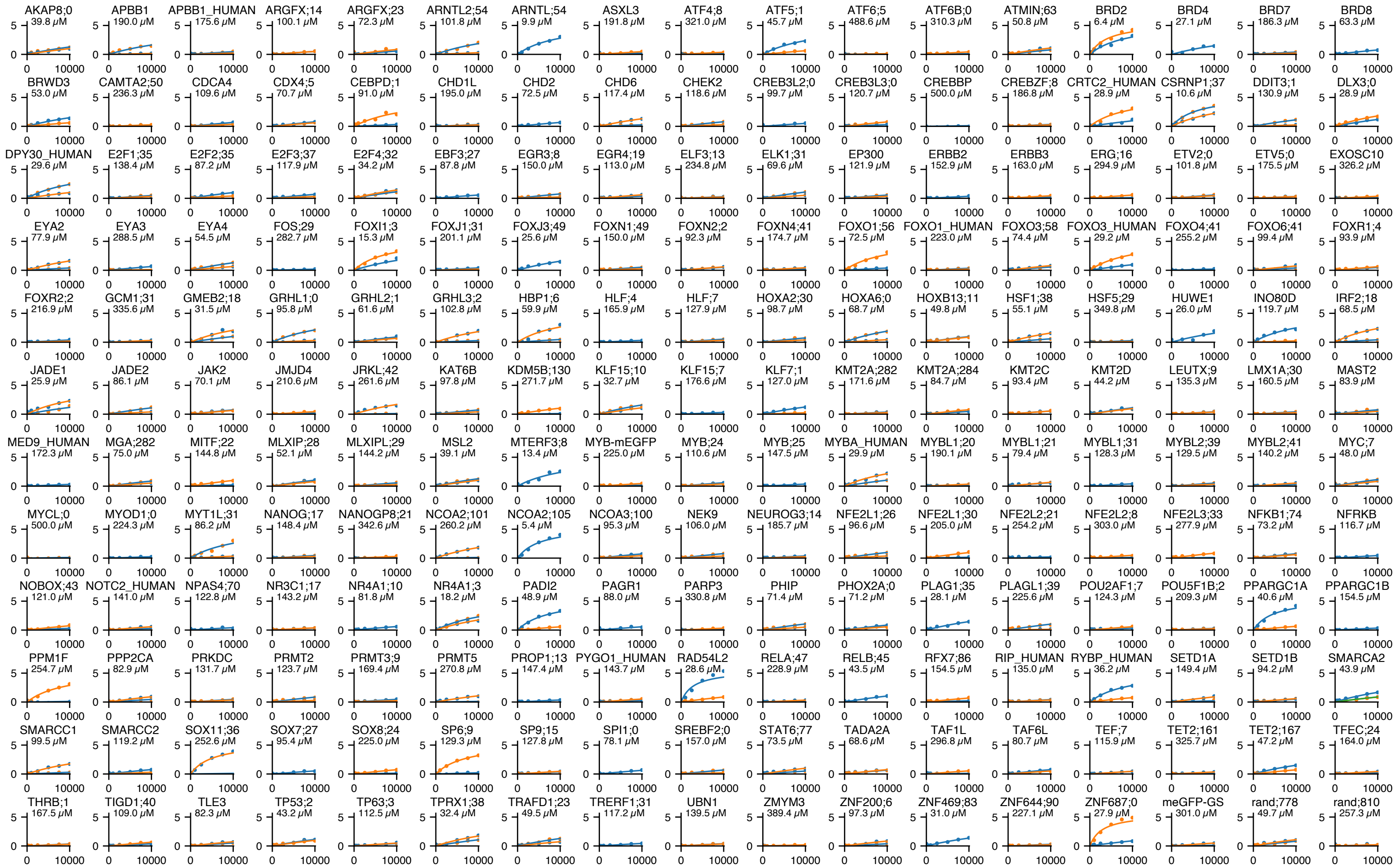


MED9 device 2:

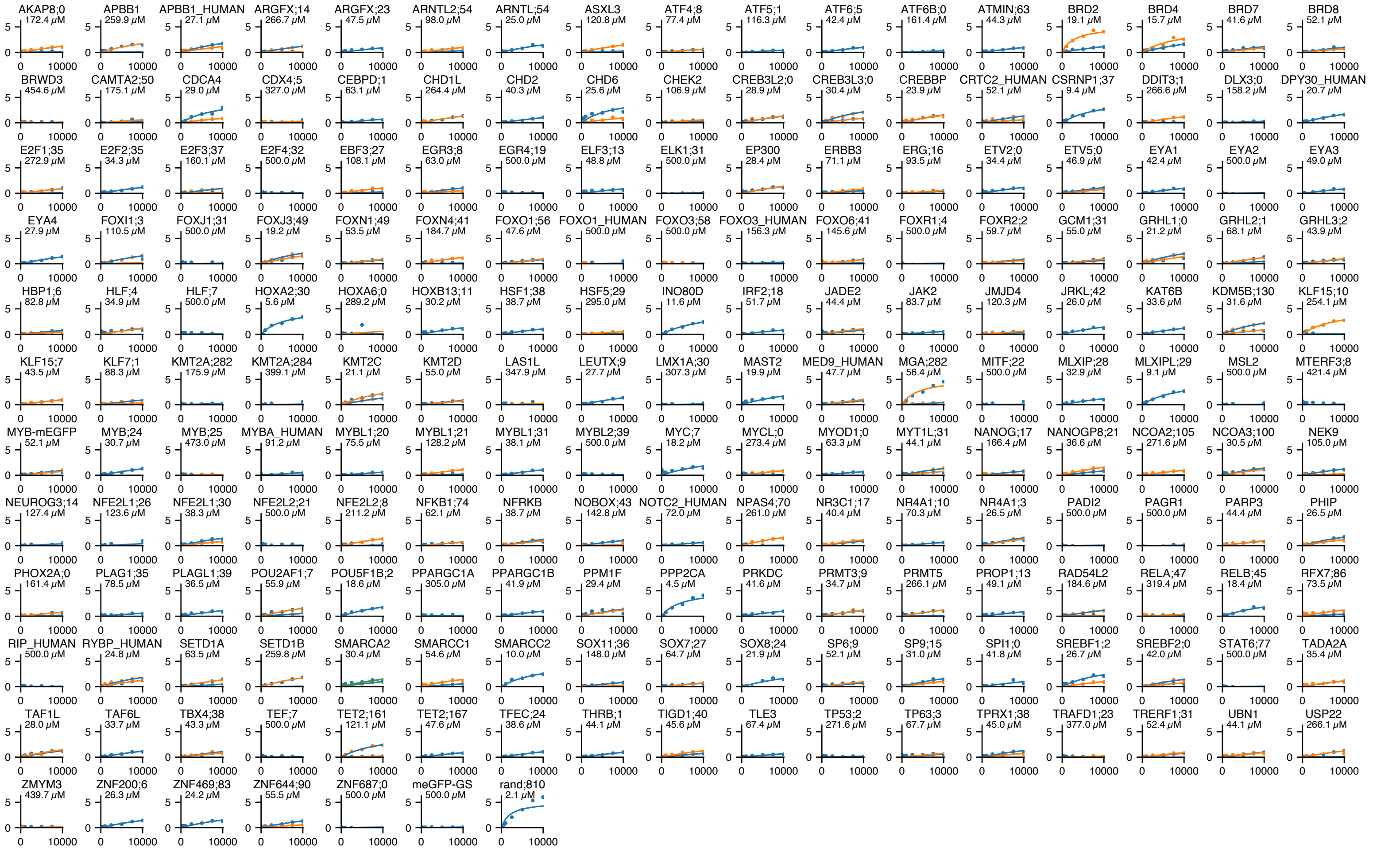

BRD7 device 2:

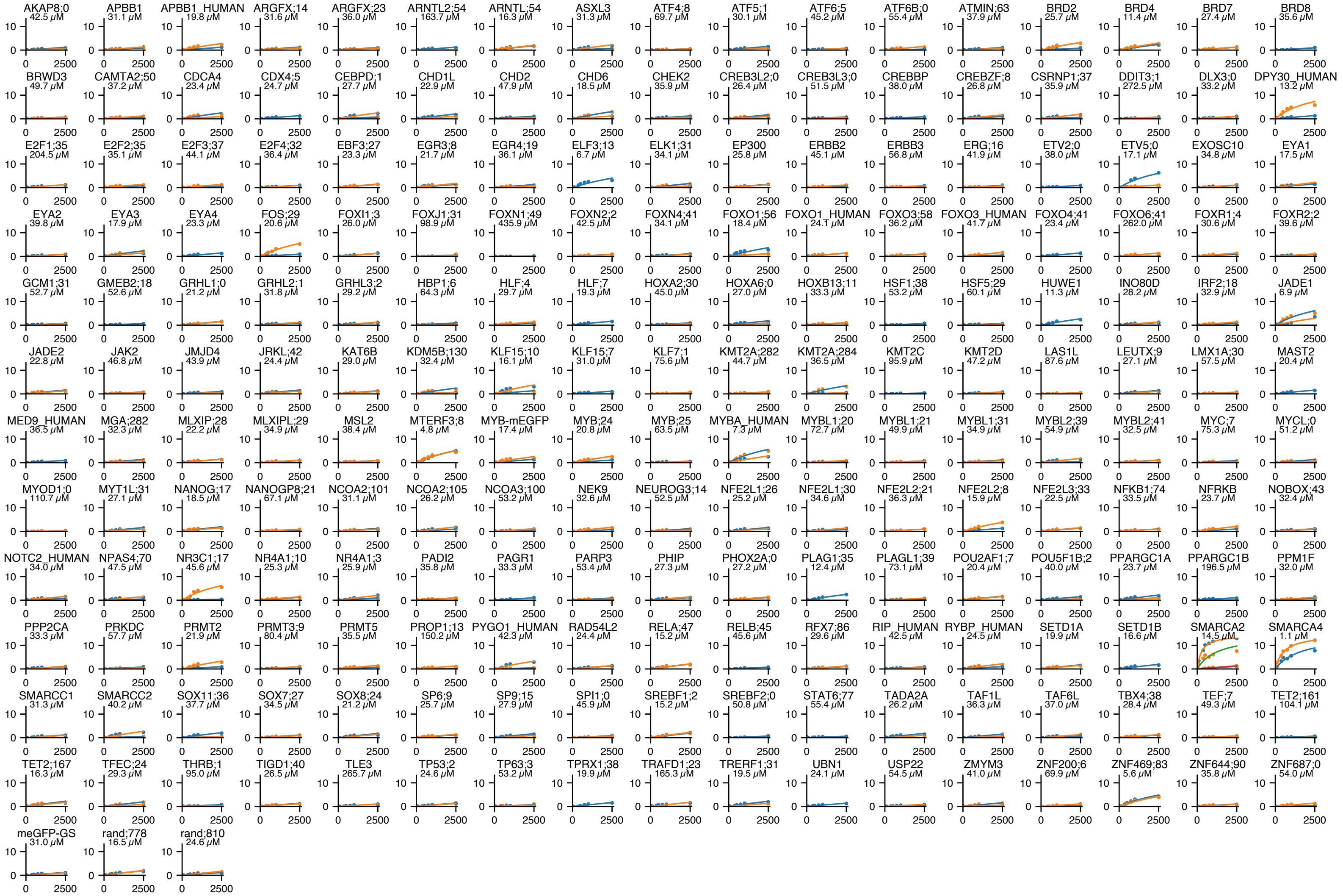


**e**

**Affinities replicability (red=meGFP):**

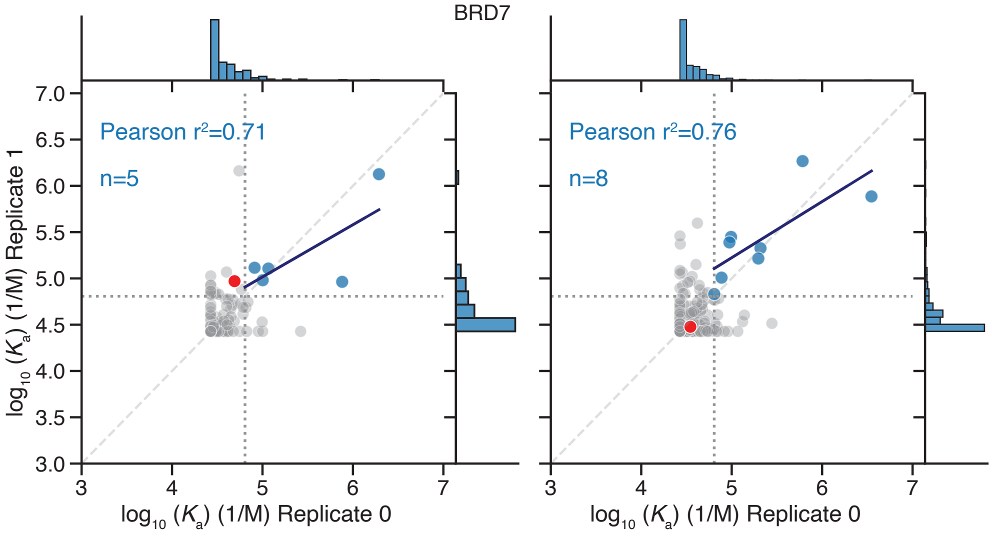

MED9

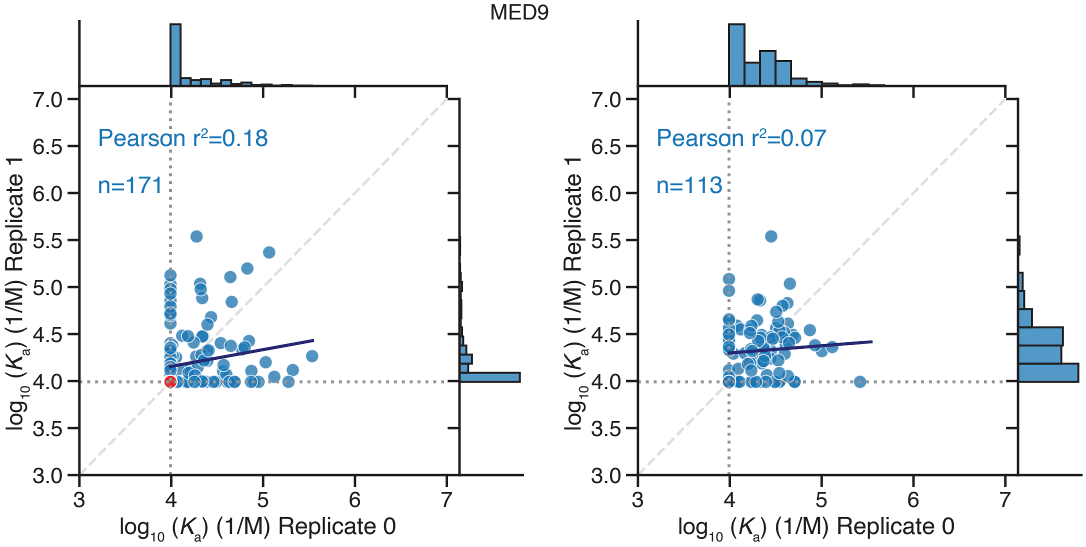


**f**

**Raw images at highest tested concentration, co-A (top) AD (bottom)**

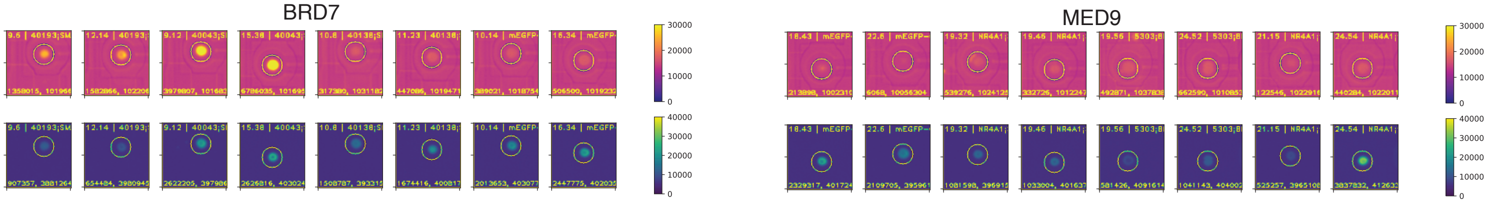
